# miss-SNF: a multimodal patient similarity network integration approach to handle completely missing data sources

**DOI:** 10.1101/2025.02.24.639805

**Authors:** Jessica Gliozzo, Mauricio A. Soto Gomez, Arturo Bonometti, Alex Patak, Elena Casiraghi, Giorgio Valentini

## Abstract

**Motivation:** Precision medicine leverages patient-specific multimodal data to improve prevention, diagnosis, prognosis and treatment of diseases. Advancing precision medicine requires the non-trivial integration of complex, heterogeneous and potentially high-dimensional data sources, such as multi-omics and clinical data. In literature several approaches have been proposed to manage missing data, but usually limited to the recovery of subsets of features for a subset of patients. A largely overlooked problem is the integration of multiple sources of data when one or more of them are completely missing for a subset of patients, a relatively common condition in clinical practice.

**Results:** We propose miss-Similarity Network Fusion (miss-SNF), a novel general-purpose data integration approach designed to manage completely missing data in the context of patient similarity networks. Miss-SNF integrates incomplete unimodal patient similarity networks by leveraging a non-linear message-passing strategy borrowed from the SNF algorithm. Miss-SNF is able to recover missing patient similarities and is “task agnostic”, in the sense that can integrate partial data for both unsupervised and supervised prediction tasks. Experimental analyses on nine cancer datasets from The Cancer Genome Atlas (TCGA) demonstrate that miss-SNF achieves state-of-the-art results in recovering similarities and in identifying patients subgroups enriched in clinically relevant variables and having differential survival. Moreover, amputation experiments show that miss-SNF supervised prediction of the overall survival and progression-free interval events with completely missing data achieves results comparable to those obtained when all the data are available.

**Availability and Implementation:** miss-SNF code, implemented in R, is available at https://github.com/AnacletoLAB/missSNF.

## 1. Introduction

Precision Medicine (PM) aims to improve disease prevention, diagnosis, prognosis, and management by enabling more effective stratification of patient sub-populations and the development of targeted therapies [Johansson et al., 2023]. These advancements are fueled by the increasing availability of large-scale, multimodal patient data (e.g., multi-omics, clinical, demographic, and wearable device data), which collectively provide a holistic view of disease states [Athieniti and Spyrou, 2022]. In particular, integrating multi-omics data enables the comprehensive biomolecular characterization of individuals, which is crucial to study complex diseases like cancer, cardiovascular disorders, and others [Karczewski and Snyder, 2018, Olivier et al., 2019].

Integrating multi-omics data presents significant challenges, addressed in the literature through four main strategies [Gliozzo et al., 2022]: (1) *Input data-fusion methods* apply factorization techniques to embed patients into an integrated patient space for clustering or classification. A notable example is MOFA+, which achieves impressive results using an efficient variational inference approach [Argelaguet et al., 2020]. (2) *Output-fusion approaches* independently process each omics view before combining results across views. However, such methods have been largely outperformed by more advanced techniques. (3) *Task-specific integrated patient embeddings*, such as graph neural networks (GNNs), showed promising results [Ryan et al., 2024], but they often lack generalization across diverse tasks, require large datasets – rare in the medical domain – and are affected by class imbalances inherent in medical data. (4) *Patient similarity network (PSN)-fusion techniques* construct unimodal PSNs for each omics view, where nodes represent patients and edges encode similarities between omics profiles. These unimodal PSNs are then fused into an integrated PSN space, supporting clustering and classification tasks. PSN-based methods, like NEMO [Rappoport and Shamir, 2019] and the widely used Similarity Network Fusion (SNF) [Wang et al., 2014], have demonstrated robust performance across multi-omics tasks, with additional benefits such as interpretability and privacy preservation [Gliozzo et al., 2020, Pai and Bader, 2018, Dai et al., 2020].

Several methods have been proposed for managing missing data [Liu et al., 2023, Ren et al., 2024], i.e. to recover subsets of features not available for subsets of patients, but in clinical practice it is quite common that some data sources are completely missing for subsets of patients, and this represents an open problem in the context of multi-omics and clinical data integration [Gliozzo et al., 2022]. Indeed, although state-of-the-art fusion methods are effective for disease subtype identification, biomarker discovery, and prognosis prediction, most cannot handle incomplete, or *“partial”*, datasets [Gliozzo et al., 2022, Flores et al., 2023] where one or more data sources are entirely missing for some patients (Supplementary Figure S6). While some methods, such as MOFA+ and NEMO, can process partial datasets, their application is often limited to clustering or classification tasks. On the other hand, despite its success in various clinical applications, SNF is not equipped to handle partial datasets. Supplementary Section S1 summarizes the state-of-the-art (SOTA) of partial data fusion techniques.

We introduce *miss-SNF*, a novel, task-agnostic data fusion approach specifically designed to integrate partial datasets. Miss-SNF extends the nonlinear message-passing strategy of SNF to reconstruct missing pairwise similarities within the fused PSNs. While several PSN-based data fusion algorithms have been designed as an extension of SNF, miss-SNF is the first to adapt the SNF diffusion process for integrating partial datasets. A preliminary version of the algorithm has been presented at an international conference [Gliozzo. et al., 2023].

Our experiments on nine diverse cancer datasets from the TCGA repository [Hutter and Zenklusen, 2018] show that miss-SNF not only excels at reconstructing integrated similarities but also produces an integrated PSN suitable for leveraging both unsupervised clustering and supervised classification tasks, outperforming SOTA methods, i.e. MOFA+ and NEMO, in various scenarios.

### Preliminaries: Similarity Network Fusion

SNF integrates multiple data sources via a cross-diffusion process that iteratively transfers information among the PSNs computed from each data source.

In more detail, SNF starts by using a scaled exponential affinity kernel to build a unimodal PSN for each view, **W**^(*s*)^ ∈ ℝ^*n*×*n*^, *s* ∈ {1, …, *m*}, where *m* in the number of views and *n* the number of patients. **W**^(*s*)^ is subsequently processed to build: (1) **P**^(*s*)^ ∈ ℝ^*n*×*n*^, a normalized PSN reflecting the “global” relationships among patients; (2) **S**^(*s*)^ ∈ ℝ^*n*×*n*^, a “local” PSN representing the topologies within data neighborhoods in *s*.

In the simplest case of two sources *s* ≠ *v*, the *t*-th iteration of the diffusion process uses the following formula to update 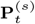 and 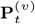:

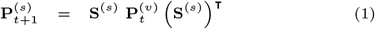

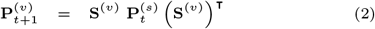

Expanding equation 1 to focus on the update of source *s* for two points *x*_*i*_ and *x*_*j*_, the update formula of **P**^(*s*)^(*i, j*) becomes:

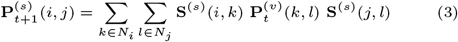

where *𝒩i* and *𝒩*_*j*_ are the *K*-nearest neighborhood of respectively *x*_*i*_ and *x*_*j*_ in *s*. The product within the sums is different from zero only when there exist points *x*_*k*_ ∈ 𝒩_*i*_ and *x*_*l*_ ∈ 𝒩_*j*_ that are also global neighbors in the other source *v* at the preceding iteration (i.e. 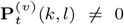) In other words, the similarity information is exchanged from source *v* to source *s* only through the common (global and local) neighborhoods they identify. This iterative message-passing process makes all the **P** global similarities progressively more similar, so that at convergence - or after *T* steps, the integrated (consensus) PSN, **P**^(*c*)^, is computed as the average of all the **P**^(*v*)^ ∀ *v* ∈ {1, …, *m*}. **P**^(*c*)^ captures shared and complementary, local and global, information from all the individual views (details in Supplementary Section S2 and Figure S7). SNF diffusion is easily adapted to *m >* 2 sources as:

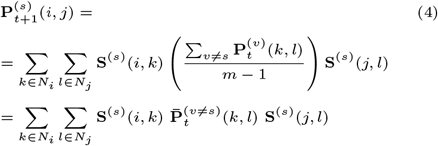

where 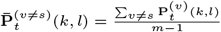

### 2. miss-SNF

Miss-SNF tweaks the three matrices **W**^(*s*)^, **P**^(*s*)^, and **S**^(*s*)^ to leverage SNF and handle partial patient samples *x*_*i*_ having completely missing information for some sources. We propose and comparatively evaluate different versions of *miss-SNF* that we named *one, zero, equidistant* and *random*.

**miss-SNF one** introduces a unique self-similarity (self-loop) for the partial sample *x*_*i*_ in all the similarity matrices of source *s* to allow message passing even when we have a completely missing source *s* for a patient *x*_*i*_. This is achieved by setting:

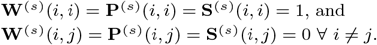

Under this setting, we have two possible different cases for computing the updated value for 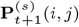 (eq. 3):

1. when *k* ≠ *i*

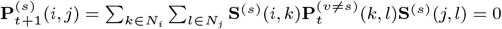
2. when *k* = *i*

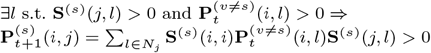

Hence eq. 3 becomes:

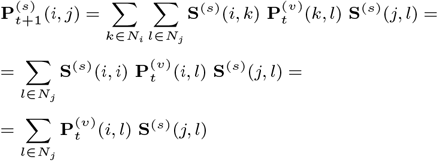

In other words, we have a contribution to the missing 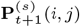 when *x*_*i*_ and *x*_*j*_ have a common neighbor *x*_*l*_ in, respectively, the “global” network **P**^(*v*)^ of the other source and in the “local” network **S**^(*s*)^ for the partial source *s*. This implies that we can populate **P**^(*s*)^(*i, j*) also when data are missing for source *s*. Figure 1 and Supplementary Figure S7 provide a pictorial description of the miss-SNF algorithm. If the dataset contains *m >* 2 sources, eq. 4 becomes:

**Fig. 1:**
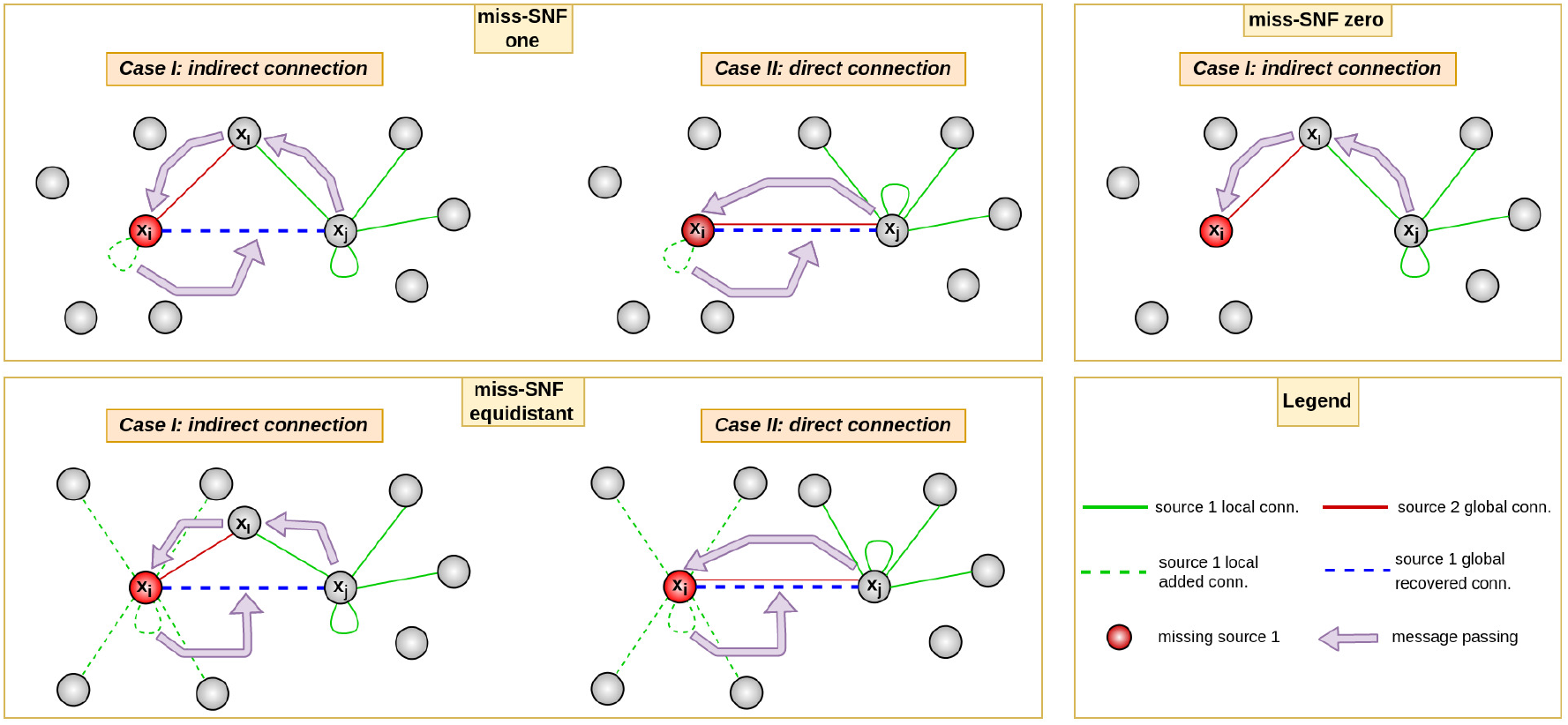
Pictorial representation of the edge recovery process in miss-SNF. Violet arrows represent the message passing mechanism across different nodes and data sources. *x*_*i*_ has completely missing data for source 1 (green edges) and a second data source 2 (red edges) is available for both *x*_*i*_ and *x*_*j*_. Miss-SNF *one* (upper left box) can recover a missing edge between *x*_*i*_ and *x*_*j*_ for source 1 (blue dotted line) if (I) there is a common neighbour *x*_*l*_ between *x*_*i*_ for data source 2 and *x*_*j*_ for the local similarity matrix of the missing data source 1, or (II) if *x*_*i*_ and *x*_*j*_ are directly connected through the source 2. In miss-SNF *equidistant* (lower left box), the patient *x*_*i*_ is initially set equally similar to all the other patients but more similar to himself and also in this case the algorithm can recover missing edges for the source 1 (blue dotted line) through indirect connections or exploiting a direct connection through source 2. Miss-SNF *zero* (upper right box), only preserves the information available for all the data sources, but does not allow for missing edge recovery, since no self-loop is added to *x*_*i*_.

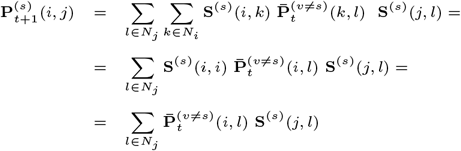

According to the above equation 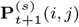 is reconstructed if *x*_*i*_ and *x*_*j*_ share common neighbours respectively in the global network 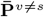 and in the local network **S**^(*s*)^.

**miss-SNF zero** simply ignores partial samples, *x*_*i*_, in the diffusion process by setting **W**^(*s*)^(*i, j*) = **P**^(*s*)^(*i, j*) = **S**^(*s*)^(*i, j*) = 0, ∀*j*. By applying computations similar to those shown for miss-SNF *one*, it is straightforward to see that 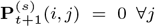 (Figure 1 and Supplementary Figure S7).

In this case, the integrated similarity **P**^(*c*)^ is computed as

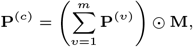

where ⊙ is a pointwise multiplication, and **M** is a matrix of the same dimension of **P** where **M**(*i, j*) is the reciprocal of the number of sources containing information for both *x*_*i*_ and *x*_*j*_. In this way we obtain a “consensus” *P* ^(*c*)^ that averages with respect to the actually available data source for each patient.

**miss-SNF equidistant** sets the self-loop on each partial *x*_*i*_ to **W**^(*s*)^(*i, i*) = 0.5 and assumes all the other samples are equally distant (Figure 1 and Supplementary Figure S7). This is achieved by setting **W**^(*s*)^(*i, j*) = 0.5*/*(*N* −1), ∀*j* ≠ *i*, where *N* is the number of patients. The same is done for **P**^(*s*)^ and **S**^(*s*)^. In this way, *x*_*i*_ is more similar to itself than to the other patients, and each patient is initially equally similar to the other patients, which is a fair assumption considering the complete lack of information about source *s* for patient *x*_*i*_.

**miss-SNF random** sets random uniformly distributed similarities for sample *x*_*i*_ in **W**^(*s*)^, **P**^(*s*)^ and **S**^(*s*)^. This method is a sort of baseline to test if miss-SNF performance is really influenced by the initial values setting of the partial sources.

## 3. Experimental Evaluation

### 3.1. Datasets

miss-SNF is evaluated on nine multi-omics datasets from TCGA [Hutter and Zenklusen, 2018]: BLadder urothelial Carcinoma (**BLCA**); BReast infiltrating ductal CArcinoma (**BRCA1**) and BReast infiltrating lobular CArcinoma (**BRCA2**); KIdney Renal Clear cell carcinoma (**KIRC**); LUng ADenocarcinoma (**LUAD**); LUng Squamous Cell carcinoma (**LUSC**); PRostate ADenocarcinoma (**PRAD**); OVarian serous cystadenocarcinoma (**OV**); SKin Cutaneous Melanoma (**SKCM**). Each dataset comprises four data sources: miRNA, mRNA, protein expression and DNA methylation. Clinical variables are downloaded and used to evaluate miss-SNF on unsupervised clustering.

The TCGA-CDR dataset [Liu et al., 2018] further provides standardized clinical features derived from the TCGA program, addressing the incomplete availability of outcome and treatment information and the relatively short follow-up durations. The curated information about the patients’ Overall Survival (OS) and Progression Free Interval (PFI) have been used to evaluate miss-SNF in supervised classification tasks.

Before any further analysis, the multi-omics datasets were pre-processed to filter variables mainly carrying noise or highly redundant information. After Z-score standardization, we reduced the effect of the small-sample-size problem by dimensionality reduction of each view. To this aim, according to Gliozzo et al. [2025], we chose the randomized principal component analysis [Erichson et al., 2019] and set the dimension of the latent space by computing the view intrinsic dimensionality (id), via the *two-nn* [Facco et al., 2017] algorithm, nowadays one of the most effective id estimators.

An exhaustive description of the TCGA datasets, their descriptive statistics, the clinical variables we downloaded, and the details of the data pre-processing can be found in Supplementary Section S3 and Table S5.

### 3.2. Experimental setup

We evaluated miss-SNF in three main experimental settings characterized by partial (completely missing) data:

a. data recovery capability, i.e. reconstructing global patient similarity
b. patients cluster recovery capability
c. recovery of phenotype/outcome prediction capabilities

To objectively assess and compare the information recovery reliability of the different miss-SNF algorithms, all the nine datasets were filtered to retain only their complete versions, excluding partial cases. This allowed the application of standard SNF, which served as a gold standard for comparison with miss-SNF applied to their randomly amputated counterparts.

We evaluated miss-SNF under varying partial-missingness conditions, by randomly amputating each dataset to remove x% (*x* ∈ {10, 20, 30, 40, 50}) of samples from each data source. The removal process was performed independently for each data source, but in such a way as to ensure that at least one data source remained available for each patient. To ensure statistical significance, *h* = 10 randomly amputated datasets were generated for each amputation percentage. Missing data patterns for all cancers are presented in Supplementary Figures S8-S16.

Regarding SNF and miss-SNF hyperparameters, all experiments were conducted with *T* = 100 iterations for all models except miss-SNF *one*, where *T* = 1000 was used. This adjustment was based on convergence analysis results, which indicated slower convergence for miss-SNF *one* (see Supplementary Section S4 for details).

In our experiments we compared miss-SNF with MOFA+ and NEMO algorithms, two state-of-the-art methods for the integration of partial datasets.

Statistical comparison was performed by computing win-tie-loss tables by paired Wilcoxon signed rank test (details in Supplementary Section S5).

A sketch of the evaluation schema is shown in Figure 2.

**Fig. 2:**
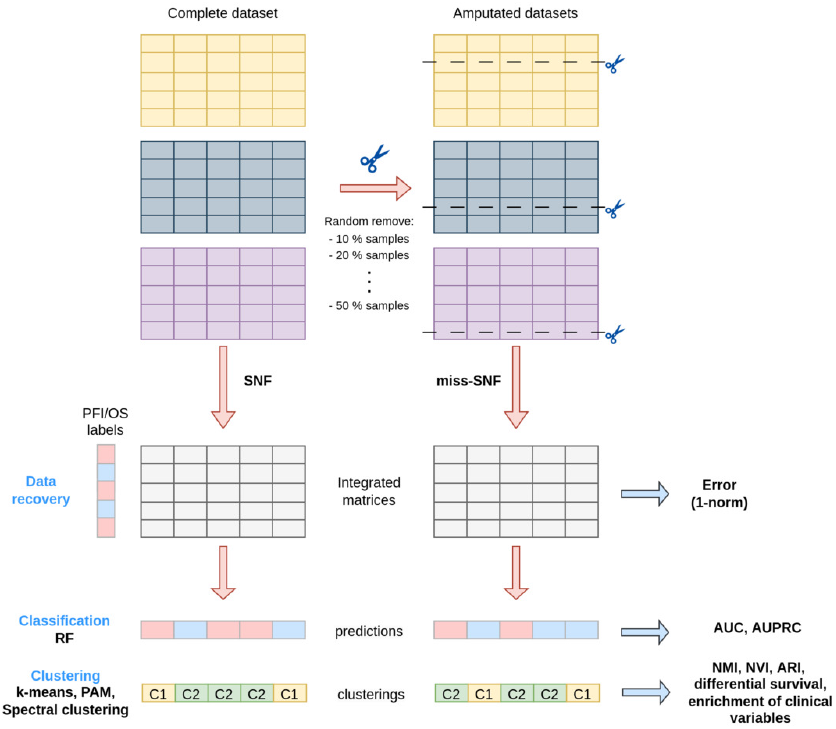
Experimental setup for miss-SNF evaluation

### 3.3. Results

#### Recovery of missing pairwise similarities

To evaluate the performance of miss-SNF in recovering information for partial samples, we applied SNF [Wang et al., 2014] to integrate the complete datasets and miss-SNF to integrate their amputated counterparts. Integration quality was assessed on each dataset by calculating the “reconstruction error” by means of the 1-norm distance between the fused PSN from the complete dataset (via SNF) and the 10 randomly generated amputated datasets (via miss-SNF) at each amputation percentage^1^. Figure 3 shows the distribution of the reconstruction errors obtained for increasing percentages of partial examples. As expected, all versions of miss-SNF, except miss-SNF *random*, show increasing difficulty (increasing reconstruction errors) in integrating data sources as the proportion of missingness rises. The behaviour of miss-SNF *random* can be attributed to its random similarity assignment for partial samples. This approach introduces bias into the integrated matrix by incorporating incorrect similarities during the diffusion process, resulting in noisy fused matrices that remain largely unaffected by the proportion of missing data.

**Fig. 3:**
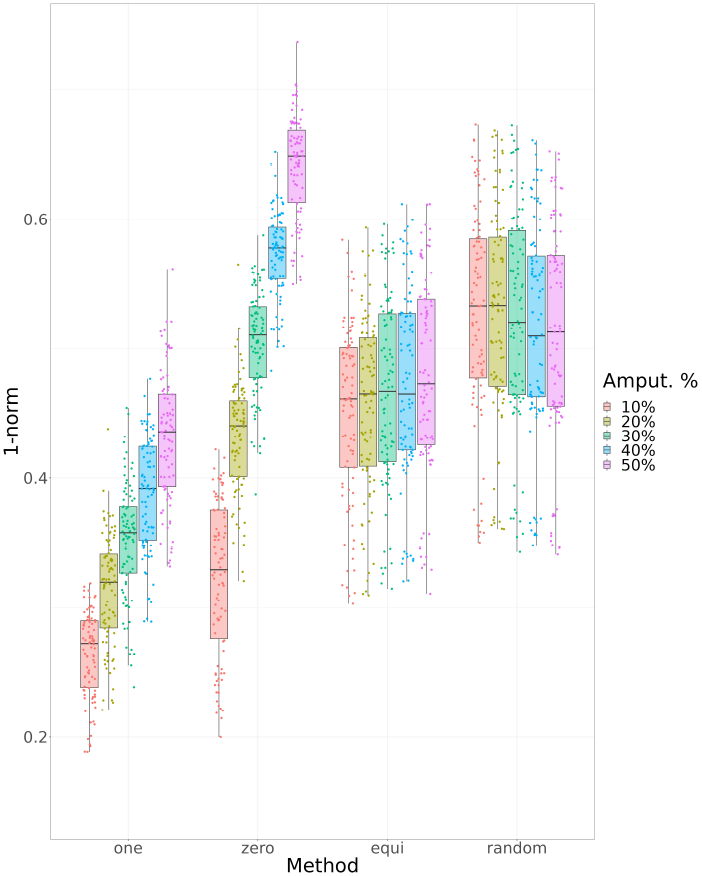
Distribution of the 1-norm difference between amputated and complete datasets for different miss-SNF algorithms (x-axis).

Overall, miss-SNF *one* and *equidistant* achieve the lowest reconstruction errors. In contrast, miss-SNF *zero* fails to recover information for partial samples, specially when the percentage of amputated data increases, as expected, since it completely disregards these samples during the diffusion process. Win-tie-loss plots in Supplementary Figure S4 show that miss-SNF *one* achieves the best results (one-sided paired Wilcoxon test, *α* = 0.05), followed by miss-SNF *equidistant*. Boxplots of the reconstruction error for individual cancers are available in Supplementary Figure S5.

Driven by these results, in the following we limit our analysis to miss-SNF *one* and miss-SNF *equidistant*.

#### Compared to SOTA methods, miss-SNF is more effective in similarity recovery

Using the same procedure, we computed the reconstruction error between the combined PSNs obtained from the complete and partial datasets via MOFA+^2^ or NEMO.

Figure 4 shows the reconstruction error distributions computed across all datasets by miss-SNF *one, equidistant*, MOFA+, and NEMO. Miss-SNF *one* and *equidistant* have the lowest reconstruction error across all amputation percentages with a slight increase for higher proportions of missing values. NEMO exhibits higher error values while the worst approach is MOFA+ with significant errors that consistently increase as the amount of partial samples increases. The win-tie-loss tables summarizing the results of one-sided paired Wilcoxon tests (Supplementary Figure S18 and Supplementary Table S6) confirm that miss-SNF (*one* and *equidistant*) outperforms NEMO and MOFA+.

**Fig. 4:**
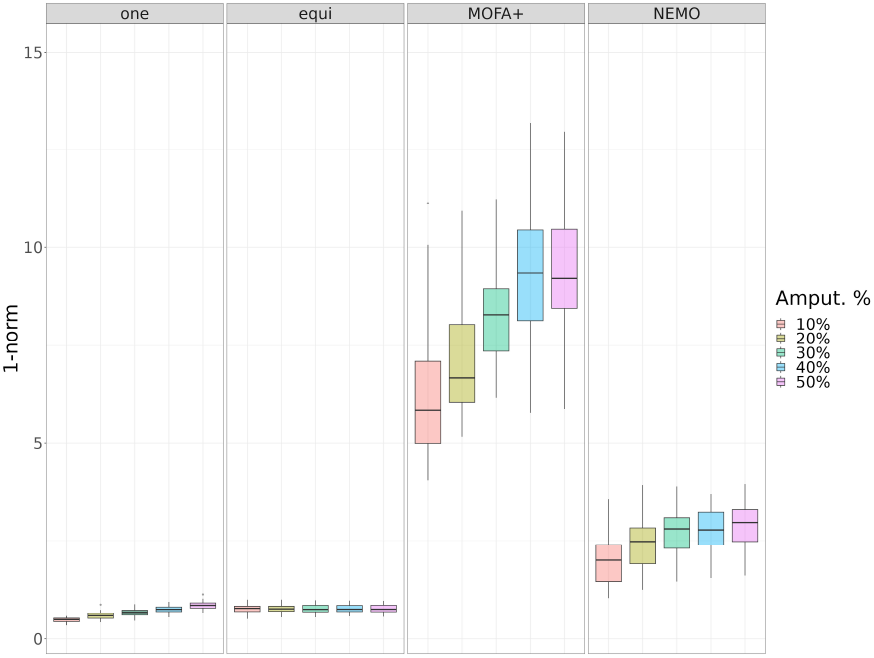
Comparison of the PSN recovery capabilities of miss-SNF *one* and *equidistant*, MOFA+ and NEMO across multiple cancer datasets. Boxplots show the distribution of the 1-norm distances among amputated and complete datasets for different data fusion methods and amputation percentages. To assure a fair comparison between different integration methods, PSN values are normalized by min-max scaling.

To further support these findings, we applied the 3-step Random Walk Kernel (3-RWK, [Sugiyama and Borgwardt, 2015]) to compare the topologies of the fused PSNs, after min-max normalization, obtained from complete and amputated datasets. The results, shown in Supplementary Figure S19, capture the impact of missing data on topological similarities. While the boxplots indicate that miss-SNF and MOFA+ maintain similar performance levels even as the percentage of partial samples increases, NEMO exhibits a decline in topological similarity between the fused PSNs from complete and amputated datasets as the proportion of partial data rises. Win-tie-loss tables summarizing the comparative statistical analysis (one-side paired Wilcoxon test) further demonstrate that miss-SNF outperforms MOFA+ and NEMO overall (Supplementary Table S7). Similar results were observed when experimenting with 5-step and 10-step RWK (data not shown).

#### Clustering experiments

We conducted clustering experiments using SNF (on complete datasets), miss-SNF (on amputated datasets), MOFA+, and NEMO (on both complete and amputated datasets). Three clustering methods were employed: (1) k-means [Hastie et al., 2009] algorithm applied both to the fused-PSN (from now on k-means (similarities)), and to the factors in the case of MOFA+ (from now on k-means (factors)); (2) Spectral Clustering (SP), using both the algorithm described in [Ng et al., 2001] (SP-sClust) and the version provided by NEMO’s authors (SP-SNFtool, applied exclusively to PSNs integrated by NEMO); (3) Partitioning Around Medoids (PAM) [Kaufman and Rousseeuw, 1990], applied to the fused-PSN (PAM (similarities)), and to the factors in the case of MOFA+ (PAM (factors)).

By combining each partial data fusion approach (miss-SNF *one* and *equidistant*, MOFA+, and NEMO) with one of the clustering methods, we derived a total of 15 distinct partial data fusion+clustering approaches (see the full list in Supplementary Table S8).

To avoid bias in determining the optimal number of clusters, as recommended by [Duan et al., 2021], we evaluated clustering results for a range from 2 to 10 clusters. Additional details regarding the experimental setup for clustering experiments are available in Supplementary Section S7.

#### Comparison between clusters obtained from complete and amputated data

Our initial analysis focused on comparing clusters derived from complete datasets with those generated from partial datasets.

Supplementary Figures S20, S21 and S22 show the computed Normalized Mutual Information (NMI), Normalized Variation of Information (NVI), and Adjusted Rand Index (ARI) [You, 2021]. Win-tie-loss results in Supplementary Tables S8, S9 and S10 summarize the statistical comparison of the computed NMI, NVI and ARI values. Independently from the considered metric, NEMO followed by SP (or k-means) is the best performing combination, followed by miss-SNF *one* (any clustering method). MOFA+ has the lowest results for all metrics. Overall, PAM algorithm decreases the performance of all the data integration methods. Focusing on the best two combinations of data fusion and clustering method (NEMO+SP SNFtool and miss-SNF *one*+k-means), we can see a steady and expected decrease in metric values (increase for NVI) as the percentage of partial data increases.

#### Cluster enrichment and differential survival analysis

As a further comparison, we assessed the clinical relevance of the identified clusters using a set of clinically relevant variables by counting the number of enriched variables per cluster (more details in Supplementary Section S7.1).

Miss-SNF *one* consistently produces clusterings enriched with more clinical variables, regardless of the clustering method employed. Conversely, MOFA+ demonstrates the lowest performance among the tested approaches (Supplementary Table S11 and Supplementary Figure S23). Among the top-performing methods, Supplementary Figure S23 reveals a gradual decline in the number of enriched variables as the percentage of partial samples increases for miss-SNF *one*, irrespective of the clustering technique used. For NEMO, however, this decline is less pronounced when spectral clustering is applied.

Furthermore, we assessed the ability of considered methods to identify clusters with differential survival, by applying the log-rank test with a 95% confidence level (*α* = 0.05). For each amputation percentage, all 10 randomly amputated versions of each cancer dataset were clustered, producing 10 log-rank p-values. These p-values were aggregated by calculating the proportion of significant tests (p-value *< α*) across the 10 repetitions. The results of the survival analysis are presented in Supplementary Figure S24 and summarized in the win-tie-loss Supplementary Table S12. Miss-SNF *one* consistently emerges as the best-performing method, followed by NEMO combined with spectral clustering and k-means clustering. In contrast, MOFA+ shows the lowest performance, regardless of the clustering technique applied.

The final evaluation aimed to further assess the ability of the partial data fusion+clustering approaches to identify clinically enriched clusters after selecting the optimal number of clusters using two methods: (I) the modified eigengap method [Rappoport and Shamir, 2019], a standard approach for determining the number of clusters based on spectral clustering properties; (II) the “Clin-Surv” strategy, an approach we propose that selects the optimal number of clusters as the configuration yielding the highest number of clinically enriched variables. In case of ties, the configuration with the lowest log-rank test p-value was chosen.

In addition to comparing clusters derived from complete and partial datasets across evaluation metrics – which yield results consistent with those obtained when all cluster numbers were considered (Supplementary Tables S13-S15) – we specifically examined the number of enriched clinical variables for clusters derived using the optimal number of clusters selected from the complete datasets. The analysis showed that miss-SNF *one* and NEMO, both combined with k-means, consistently achieved the largest proportions of clinically enriched variables (Supplementary Table S16). By contrast, all variants of MOFA+ exhibited the lowest performance. Supplementary Figure S28 shows that at low levels of missingness, clusters produced by *miss-SNF* generally exhibit a greater number of enriched clinical variables compared to clusters generated by NEMO. However, as the proportion of partial samples increases, the number of enriched variables in *miss-SNF* clusters progressively decreases, eventually converging toward the average values observed in NEMO clusters.

The results obtained from logrank tests are more difficult to interpret due to the high number of ties in the win-tie-loss (Supplementary Table S17). It is however clear that miss-SNF and NEMO, both followed by k-means are again the best performing models.

#### Classification experiments

We conducted classification experiments using miss-SNF *one* and *equidistant* (Section 3.3). Experiments were performed by applying Random Forest (RF) [Breiman, 2001] classifier on the integrated matrices. Following TCGA-CDR recommendations, supervised classification was applied to predict OS and PFI events. Datasets with high class imbalance (BRCA1, BRCA2, and PRAD) were not considered in the supervised classification experiments due to their low number of positive samples, which limits predictive accuracy and statistical reliability of models while complicating metric interpretation^3^. Unbiased classification performance was evaluated by using 10 random stratified holdouts (90% training set, 10% test set) and by computing the Area Under the ROC Curve (AUC) and Area Under the Precision-Recall Curve (AUPRC) as evaluation metrics. Feature selection and hyper-parameter tuning were performed using 10 internal holdouts (additional details in Supplementary Section S8). Results were summarized by computing the average and pooled standard deviation [Moore et al., 2021] across the 10 amputations randomly generated for each amputation percentage, folds and datasets. Supplementary Figure S17 compares the AUC and AUPRC obtained by miss-SNF. The classification metrics only show a slight decrease with the random amputation of increasing percentages of samples in both tasks, confirming the robustness and reliability of miss-SNF *one* and *equidistant*. Miss-SNF *one* exhibits a more gradual decline in performance compared to miss-SNF *equidistant*, highlighting its superior ability to reconstruct missing information.

## Discussion

In PSN multi-modal data integration is quite common that not all the sources of omics or clinical data are available for all patients. Miss-SNF provides a simple strategy to recover missing similarities for subsets of patients, allowing the construction of global similarity networks **P** even with patients having completely missing data for some sources.

Thorough evaluations on nine multi-omics cancer datasets with increasing levels of missing data (10%–50%) showed the following strengths of miss-SNF *one* and *equidistant*.

### Robust recovery and integration

when compared to SNF application on the complete datasets miss-SNF *one* and *equidistant* showed to excel at similarity recovery and integration, outperforming SOTA methods like MOFA+ and NEMO.

### Theoretically grounded strategy

unlike some other SOTA methods - e.g. NEMO, that apply a smart workaround to deal with missing data, miss-SNF directly leverages patient neighborhoods, making it theoretically grounded and versatile.

### Effectiveness in unsupervised clustering

clusters generated with miss-SNF *one* on partial datasets closely matched those from full datasets while identifying clinically relevant patient groups. miss-SNF *one* outperformed competing methods in enriching clinical variables and achieving significant log-rank tests, particularly when the optimal cluster count was selected via the proposed *clin-surv* method.

### Supervised learning potential

using RFs, we showed that the PSN computed by miss-SNF *one* and *equidistant* using partial data achieved perfomance very close to that obtained when the complete full data set is available, highlighting their suitability for inductive and also transductive learning tasks [Zhu and Ghahramani, 2002, Zhu et al., 2003, Valentini et al., 2014], crucial for multi-omics datasets with scarce manual annotations [Cappelletti et al., 2023].

### Advantages over SOTA methods

unlike techniques tailored to specific tasks (e.g., clustering with MaDDA [Vitali et al., 2018] or SUMO [Sienkiewicz et al., 2022], classification with DeepIMV [Lee and van der Schaar, 2021] or MOGDx [Ryan et al., 2024]), the miss-SNF algorithm produces a general-purpose integrated matrix suitable for both supervised and unsupervised tasks. It is particularly advantageous for biomedical datasets, usually characterized by low sample cardinality, where deep learning methods struggle due to sample size limitations.

In summary, miss-SNF *one* stands out as a robust, flexible solution for partial data fusion, excelling in recovery, clustering, and classification tasks while adapting to real-world challenges of incomplete data. To the best of our knowledge, this is the first algorithm to extend SNF message-passing process for partial dataset integration, setting a new benchmark for multi-omics data analysis [Gliozzo. et al., 2023].

While we performed our tests on multi-omics data, we note that miss-SNF inherits the advantages of SNF; it is therefore a versatile tool that may be used to integrate other data-types, whenever proper pairwise similarity metrics are used. Additionally, it does not rely on *a priori* biological knowledge, making it suitable for combining diverse patient data, such as clinical records or imaging features, and adaptable across various research fields beyond omics.

Although we showed the effectiveness of miss-SNF for both unsupervised and supervised tasks, this work has some limitations. Miss-SNF *one* is the best-performing version of our algorithm; however, it exhibits slower convergence when compared to both SNF and the other miss-SNF variants, which may limit its efficiency for large-scale or time-sensitive applications. Furthermore, while our tests on supervised classification showed that using partial data miss-SNF can obtain results close to that achievable when full data are available, the results were still suboptimal. This can be attributed to the inherent difficulty of the chosen tasks and the complexity of the used datasets. These limitations highlight the need for further optimization of the algorithm and the exploration of more sophisticated classification approaches.

## Supplementary data

Supplementary data are available at *Bioinformatics* online.

## Author contributions statement

J.G., M.S.G., E.C., G.V. conceptualized the work and methodology; J.G. and A.B. performed data curation; J.G. carried out formal analysis; G.V., E.C. and A.P. funding acquisition; J.G. and M.S.G. carried out experiments and visualization; J.G. and M.S.G. implemented the software; G.V., E.C. supervised this work; J.G., E.C. and G.V. wrote the paper. All the authors reviewed this work.

## Conflict of interest

The authors declare no competing interests.

## Funding

This work is supported by FAIR (Future Artificial Intelligence Research) project, funded by the NextGenerationEU program within the PNRR-PE-AI scheme (M4C2, Investment 1.3, Line on Artificial Intelligence). This work was realised with the collaboration of the EC JRC under the Collaborative Doctoral Partnership Agreement *N* ^°^35454. Computational resources were provided by the INDACO Core facility (University of MILAN).

## Data availability

The Cancer Genome Atlas (TCGA) legacy data are available to download from https://gdac.broadinstitute.org/runs/stddata2016_01_28/. TCGA-CDR dataset is included in the “Supplemental information” of Liu et al. [2018] manuscript.

## Code availability

miss-SNF is publicly available at: https://github.com/AnacletoLAB/missSNF.

## Footnotes

1 The reconstruction error is computed using the formula 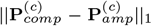, where 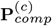 and 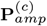 are the integrated matrices obtained from the complete and amputated datasets respectively.

2 To compute a PSN with MOFA+ where needed - e.g. when measuring the reconstruction error or when using the spectral clustering algorithm, we applied the scaled exponential Euclidean kernel [Wang et al., 2014].

3 Addressing such imbalance would require additional techniques (e.g., resampling, cost-sensitive approaches) that could complicate perfomance assessment in a real-world scenario.

## Notes

### Competing Interest Statement

The authors have declared no competing interest.

