## Supplementary Text for "miss-SNF: a multimodal patient similarity network integration approach to handle completely missing data sources"

[1]Dipartimento di Informatica "Giovanni Degli Antoni", Università degli Studi di Milano, Via Giovanni Celoria 18, 20133, Italy, [2]European Commission, Joint Research Centre (JRC), Ispra, Italy, [3]Department of Biomedical Sciences, Humanitas University, Via Rita Levi Montalcini 4, 20072, Pieve Emanuele (MI), Italy, [4]Department of Pathology, IRCCS Humanitas Clinical and Research Hospital, Via Alessandro Manzoni 56, 20089, Rozzano (MI), Italy, [5]Environmental Genomics and Systems Biology Division, Lawrence Berkeley National Laboratory, Berkeley, Ca, USA and [6]ELLIS - European Laboratory for Learning and Intelligent Systems, Milan, Italy

### Contents


S1 State-of-the-art data integration methods for "partial datasets" 1
S2 Similarity Network Fusion 3
S3 Dataset preparation 5
S3.1 High-pairwise correlation analysis and filtering 6
S3.2 Clinical variables for miss-SNF 7
S4 miss-SNF convergence experiments 14
S4.1 Convergence analysis 14
S4.2 Partial data reconstruction error 17
S5 Statistical analysis 19
S6 Pairwise similarity reconstruction in the fused PSNs: additional results 19
S7 Clustering experiments 19
S7.1 Experimental setup for clustering analysis 19
S7.2 Clustering algorithms 22
S8 Classification experiments - supervised feature selection and hyper-parameter tuning via internal-holdout validation 23
S9 Supplementary Tables 24


### S1. State-of-the-art data integration methods for "partial datasets"

This section collects representative state-of-the-art methods that can integrate partial datasets focusing on their used strategy to handle missing data (further methods are available in [Flores et al., 2023]). Tables S1 and S2 allow a quick comparison among these data fusion approaches.

In the category of input-data fusion methods that leverage matrix factorization (i.e. MF-based Gliozzo et al. [2022]) for the integration of different data views, there are some interesting approaches that can manage partial datasets. **DFMF** (Data Fusion by Matrix Factorization) Žitnik and Zupan [2014] exploits a penalized non-negative matrix tri-factorization where the different data views are modeled as non-negative relational matrices $\mathbf{R}_{ij}$ relating two object types $\epsilon_i$ and $\epsilon_j$ and, optional, constraint matrices that represent connections between the same type of objects. In the case of multi-omics data, our objects could be patients, genes, proteins, etc. The concurrent factorization of all constrained $\mathbf{R}_{ij}$ returns factors that are specific to each modality and factors that characterize each object, ultimately providing an integrated view of the data. This method is flexible in allowing to integrate the considered data sources even if some relational matrices are missing or sparse, it is only necessary that all relational matrices are connected. This possibility stems from how the objective function is designed. Moreover, this approach makes no assumption about the structural properties of the relational matrices. After training a model and computing the matrix factors, we have the possibility of using the trained model to reconstruct missing values in incomplete relational matrices. In this way, partial datasets can be imputed. Inspired by DFMF, **MaDDA** (Matrix trifactorization for Discovery of Data similarity and Association) Vitali et al. [2018] exploited matrix tri-factorization to compute integrated matrices having patients on the rows and factors on the columns, were factors are interpreted as clustering groups allowing to assign each patient to the group with maximum value. Multiple repetitions of the factorization with different initializations allow to obtain a consensus matrix through the averaging of the estimated matrices and an integrated PSN, where the connection between two samples represents how many times a pair of samples is assigned to the same group/cluster. **MOFA+** Argelaguet et al. [2020, 2018] (Multi-Omics Factor Analysis+) is the most notable and widely used matrix factorization approach developed for the integration of multi-omics data. It describes each view as the product of two matrices: a shared factor matrix and a view-specific matrix of weights. The factor matrix is shared across all views and it represents in a lower dimensional space the principal sources of variability shared among the different modalities, while a matrix of weights is generated for each view and encodes the description of the features in the view with respect to a set of factors. In particular, MOFA+ uses a Bayesian approach where the sparsity of feature/factor matrices is enforced by selecting a set of appropriate prior distributions that impose the regularization on the weights. Regularization leads to more interpretable decomposed matrices, where not all factors are active and not all features actually contribute to each factor. Inference of matrix coefficients is performed using a variational inference approach that seeks to minimize the (Kullback-Leibler) divergence from the actual posterior distribution. Missing data (including completely missing samples) are not considered in the likelihood and they are neglected in the update equations exploited to train the model by the use of a binary mask for each view, where 1 is set if a feature is observed for a specific sample and 0 if it is not. MOFA+ also supports the analysis of multi-view and multi-group data (e.g. multi-omics data where groups of samples were subjected to different experimental conditions).

Interesting embedding techniques are DeepIMV Lee and van der Schaar [2021] and MvNE Mitra et al. [2020]. **DeepIMV** (Deep Variational Information Bottleneck) Lee and van der Schaar [2021] is a deep learning approach having the objective of finding task-relevant embeddings of multi-omics data considering both the joint and marginal aspects of observations, where the tasks can be classification or regression tasks. Two fundamental principles are exploited by this approach. Firstly, the method adopts the *information bottleneck* (IB) principle Tishby et al. [2001] aiming to construct a task-specific representation balancing the trade-off between succinctness and predictive capabilities. Second, the method enforces an embedding leveraging both the marginal and joint elements of the different views. To this end, the proposed network architecture is composed of four main modules: (i) *view-specific encoders* maps observations from each single-view into a common latent space; (ii) the *product-of-experts (PoE)* integrates the single view marginal latent representations into a joint latent space; (iii) the *multi-view predictor* generates task-specific predictions from the joint representation computed by PoE; (iv) *view-specific predictors* generates prediction from the marginal single-view representation computed by the view-specific encoders. In case of missing views, the PoE module simply ignores missing views when finding the joint representation. The architecture of encoders and predictors is based on multi-layer perceptrons. The view-specific and joint embeddings are tailored to the considered predictive task but can be exploited for further data analysis (e.g. clustering) and to construct a task-specific PSNs. **MvNE** (Multi-view Neighbourhood Embedding) Mitra et al. [2020] attempts to use a combination of two techniques: a *probabilistic integration* of similarity between different views, and an *embedding* technique into a low-dimensional space in order to minimize the divergence with this combined similarity. The method starts by computing, in each view, the similarity matrix between samples using a scaled squared Euclidean distance and replacing the missing features with zeros. The resulting similarities are normalized and interpreted as a Gaussian probability distribution of two samples to be neighbours. The resulting distributions on the different views are thus integrated using their conflation, computed as the normalized product of the probability densities. Samples are embedded into a low-dimensional metric space using a stochastic neighbours embedding technique (t-SNE) Van der Maaten and Hinton [2008] assuming, in the target space, a Student t-distribution. Therefore, the embedding is obtained by minimizing the Kullback-Leibler divergence between the distributions computed in the former space and in the new low-dimensional spaces. Starting from an initial embedding, obtained through a stacked autoencoder composed of denoising autoecoders in each layer, the minimization task is performed using a stochastic gradient descent approach. The final embedding is exploited to obtain a clustering using an ad-hoc heuristic called AMOSA (Archived Multi-objective Simulated Annealing) Acharya et al. [2015]. In this heuristic, Simulated Annealing is used to find the clusters centers which minimize a bi-objective function accounting simultaneously for the cluster compactness and cluster separation. This method supports the integration of partial datasets in the computation of the integrated probability distribution. The integrated probability of samples having multiple views is computed by conflation considering all the available views, otherwise the single-view Gaussian probability distribution is exploited.

The computation of a lower-dimensional representation of the data is not the only strategy to integrate partial datasets. It is possible also to exploit the unimodal PSN for integration, as done by PSN-fusion methods. A method popular for its effectiveness and fast computation is **NEMO** (NEighborhood based Multi-Omics clustering) Rappoport and Shamir [2019]. Initially a PSN is constructed for each data view using a scaled Gaussian kernel of Euclidean distance, where the scaling factor is needed to normalize pairwise distances taking into account the neighbourhoods of samples. Of note, this normalization is used also by SNF Wang et al. [2014]. Then, a relative similarity matrix is computed from each unimodal PSN to capture the local relationships between observations. The (local) relative similarity matrix measures the similarity between two points relative to their neighbourhoods and it is designed to have values equal to zero for nodes that are not neighbors. The final integrated matrix is obtained by averaging the unimodal relative matrices. Spectral clustering is then applied on the integrated similarity matrix. Extensive experiments on both simulated and real datasets showed the effectiveness and efficiency of NEMO with respect to 9 state-of-the-art methods including one MKL-based method, a spectral clustering approach, k-means, and 6 clustering methods exploiting an input data-fusion strategy. NEMO can fuse also samples having completely missing views where the similarity between two observations is computed by considering the average with respect to the views where both observations are present. Thus, the main limitation of NEMO is that it requires for each pair of samples the presence of at least a common data view to allow the computation of a similarity in the final integrated matrix. This limitation is handled in the implementation of NEMO by setting the similarity of patients that do not share data views to the mean value of the available similarities in the lower triangular integrated matrix. This limitation is overcome by **MSNE** (Multiple Similarity Network Embedding) Xu et al. [2021], which allows the integration of partial datasets through a well-designed random walk strategy. Random walks capture the neighbor relationships among samples in different unimodal PSN. The PSNs, representing the local similarities among samples with respect to different data sources, are normalized into transition probability matrices, whose values are re-estimated through the propagation of the local similarities by a Markov process. The PSNs are then connected by common samples in different networks, allowing the random walker to move to other PSN without the constrain of requiring all samples to be present in all networks. Multiple random walks starts from each sample, and the resulting set of node sequences is embedded into an integrated low-dimensional matrix using a word2vec network with a Skip-gram Mikolov et al. [2013] architecture. The final embedding is then used for clustering by k-means. Following the state-of-the-art performance of Graph Convolutional Networks (GCNs) for classification tasks, **MOGDx** (Multi-Omics Graph Diagnosis) [Ryan et al., 2024] is a learning approach, recently published as the first GCN-based method able to integrate any number of modalities while handling patients having missing data sources. MOGDx exploits a GCN to solve classification tasks in heterogeneous diseases like cancer. In particular, SNF is used to fuse the unimodal PSNs, therefore obtaining an integrated PSN. On the other hand, a multi-modal encoder computes node features by pooling all the modalities into a reduced and integrated representation, where median imputation is exploited to retain missing patients in each modality. The PSN is then used to define homogeneous graph edges connecting nodes whose feature representation is given by the encoder output. Such graph is input to the GCN training. The multi-modal encoder and the GCN weights are jointly learned by backpropagation. SNF does not natively support missing data sources; therefore, MOGDx implicitly handles partial sources $s$ having the partial patient $x_i$, by setting $\mathbf{W}^{(s)}(i,j) = 0 \forall j$ (see Subsection "Preliminaries: Similarity Network Fusion" in the main file).

Another possibility is to analyse separately the PSN obtained from each data source and then integrate the obtained results. This is the strategy adopted by output-fusion techniques. An example of this kind of approaches is **SUMO** (Subtyping tool for multi-omic data) Sienkiewicz et al. [2022]. SUMO initially constructs a PSN for each data source where the similarity is computed considering only the common set of features present for each pair of samples if the common set has sufficient cardinality (by default 10%), otherwise the similarity among samples is missing. The method formulates a constrained non-negative tri-factorization of the similarity matrices, obtaining a shared sparse representation of the samples across all views in a lower-dimensional space. The dimension of the final space is set directly as the number of clusters, thus representing the membership of each sample to each factor/group. The procedure is repeated multiple times considering different subset of samples and initial conditions. The clusterings obtained are pooled exploiting consensus clustering. In particular, the factorization is formulated as an optimization problem, where the objective function minimizes the difference between the original data matrix and its approximation by matrix factorization, while using also a term to enforce sparsity on the matrix of basis components. Missing samples, which results in missing pairwise similarities in the unimodal PSN, are masked in the objective function through an indicator matrix encoding the presence/absence of an edge in the similarity network of each data source. Another approach that exploits consensus clustering is **MONET** (Multi Omic clustering by Non-Exhaustive Types) Rappoport et al. [2020], which is an integration method designed to perform clustering. The method constructs a family of subsets of samples called modules; modules are different from clusters since they are constructed based on only a subset of views, and they do not form a partition, thus some samples could not belong to any module. This property allows MONET to deal with missing information since not all omics must be considered when constructing the modules, thus two samples can be similar even if they are not comparable in all the views. In practice, partial samples are inserted in all omic networks, but they are considered as nodes without edges in the missing data sources. Nodes without edges have no effect in the decision of assigning a sample to a module. The method is built upon NEMO Rappoport and Shamir [2019] by creating, in each view, several cluster partitions using different random samples. A weight is assigned to each couple of samples in a view based on how often they appear together in a cluster with respect to average co-occurrence in the view. Module construction is then cast as an optimization problem with the aim of maximizing the module intra-sum of weights along the specific views considered in the module. Since the resulting problem is NP-hard, heavy modules are obtained using a greedy heuristic which, starting from an initial module construction, locally improves the solution applying a set of predefined module transformations (e.g. add/remove a sample to/from a module, split/merge modules, move a sample between modules, remove/add an omic to a module, etc.). Transformations are applied to maximize the marginal gain; thus the method converges to a local maximum.

### S2. Similarity Network Fusion

Similarity Network Fusion (SNF) [Wang et al., 2014] is a method that exploits a cross-diffusion process to pass information among PSN built from different data sources, until convergence to an integrated network. The first step of SNF is the computation of a PSN $\mathbf{W}$, expressing the pairwise similarity between the biomolecular profiles of individuals $x_i$ and $x_j$, from each data modality. A scaled exponential similarity kernel is exploited to compute similarity for continuous data[1]:

$$\mathbf{W}(i,j) = exp\left(-\frac{\rho(x_i,x_j)^2}{\mu\varepsilon_{ij}}\right) \tag{S1}$$

where $\rho(x_i,x_j)$ is the Euclidean distance between patients, $\mu$ is a hyperparameter related to the variance of the local model and $\varepsilon_{i,j}$ is a scaling factor taking into account the neighbourhoods $N_i$ and $N_j$ of the considered patients:

$$\varepsilon_{i,j} = \frac{mean(\rho(x_i,N_i)) + mean(\rho(x_j,N_j)) + \rho(x_i,x_j)}{3} \tag{S2}$$

From the initial PSN $\mathbf{W}$, other two matrices $\mathbf{P}$ and $\mathbf{S}$ are computed for each data modality. The "global" similarity matrix $\mathbf{P}$, which is essential to capture the overall relationships between patients, is computed through the following normalization:

$$\mathbf{P}(i,j) = \begin{cases} \frac{\mathbf{W}(i,j)}{2\sum_{k\neq i}\mathbf{W}(i,k)} & \text{, if } j \neq i \\ 1/2 & \text{, if } j = i \end{cases} \tag{S3}$$

where for equation S3 the property $\sum_j \mathbf{P}(i,j) = 1$ holds. Then a "local" similarity matrix is obtained as follows:

$$\mathbf{S}(i,j) = \begin{cases} \frac{\mathbf{W}(i,j)}{\sum_{k\in N_i}\mathbf{W}(i,k)} & \text{, if } j \in N_i \\ 0 & \text{, otherwise} \end{cases} \tag{S4}$$

where $N_i = \{x_k | x_k \in kNN(x_i) \cup \{x_i\}\}$. $\mathbf{S}$ is able to capture the local structure of the network because considers only local similarities in the neighbourhood of each individual, setting to zero all the others.

Given $m$ data modalities, $m$ different $\mathbf{W}$, $\mathbf{S}$ and $\mathbf{P}$ matrices are constructed and an iterative process is applied where similarities are diffused through the $\mathbf{P}$s until convergence, that is, until all the matrices $\mathbf{P}$ become similar. In the simplest case, when $m = 2$, we have $\mathbf{P}_t^{(v)}$ that refers to $\mathbf{P}$ matrices for data $v \in \{1,2\}$ at time $t$. In this case, the following recursive updating formulas describe the diffusion process:

$$\begin{aligned} \mathbf{P}_{t+1}^{(1)} &= \mathbf{S}^{(1)} \times \mathbf{P}_t^{(2)} \times \mathbf{S}^{(1)\intercal} \\ \mathbf{P}_{t+1}^{(2)} &= \mathbf{S}^{(2)} \times \mathbf{P}_t^{(1)} \times \mathbf{S}^{(2)\intercal} \end{aligned} \tag{S5}$$

In other words $\mathbf{P}^{(1)}$ is updated by using $\mathbf{S}^{(1)}$ from the same data source but $\mathbf{P}^{(2)}$ from a different view and vice-versa.

[1] As noted by SNF algorithm authors [Wang et al., 2014], the proposed distance measure is not appropriate for discrete/categorical variables, where they suggest using chi-squared distance, and for boolean (binary) variables, where agreement measures are better suited.

**Table S1.** Data integration methods for partial datasets. For each method, the table reports: the name/acronym with the corresponding reference paper; the dataset used to develop and evaluate the approach in the reference paper and the corresponding sample cardinality and data types composing the dataset; the exploited integration method; the application task and the code availability (with link to the repository and programming languages for which the code is available).

| Name | Dataset | Sample Cardinality | Data type | Integration approach | Task | Code |
|---|---|---|---|---|---|---|
| MOFA+ [1] Argelaguet et al. [2020] | 3 single-cell mouse datasets | 16152 cells<br>3069 cells<br>1828 cells | mRNA<br>methy<br>mRNA, methy, DNA accessibility | NMF | Data Analysis | Python, R |
| DFMF [2] Žitnik and Zupan [2014] | 11 data sources | | mRNA-conditions, genes-GO terms, PMIDs-gene, gene-pathways, MeSH-PMIDs, GO terms-gene products, pathways-GO terms associations | MTF | Sup. class. (gene function) | Python |
| | 6 data sources | | compounds-actions, compounds-PMIDs, compounds-depositor, depositor-depositor categories, compounds-substructure fingerprints | | Sup. class. (pharmacologic actions) | |
| MaDDA Vitali et al. [2018] | Integration of: TCGA AML, BioGRID, KEGG, Disease Ontology, DisGeNET. | 200 | mutations, mRNA, clinical data gene-gene, gene/disease-pathway, disease-disease, disease-gene relations | MTF | Unsup. Clust. (Pt's subtype) | MATLAB |
| | 25 synthetic datasets | 200 | same as previous dataset | | | |
| MSNE Xu et al. [2021] | pan-cancer | 2204 | mRNA<br>CNV<br>methy | Embedding of Random Walks | Unsup. Clust. (cancer type) | Python |
| | 10 TCGA datasets | 4941 | mRNA<br>miRNA<br>methy | | Unsup. Clust. (Pt's subtype) | |
| | image dataset | 2000 | pixels averages, Fourier coefficients | | Unsup. Clust. (digits clust.) | |
| NEMO Rappoport and Shamir [2019] | 10 TCGA datasets | 3168 across datasets | mRNA<br>miRNA<br>methy | Average of local PSN | Unsup. Clust. (Pt's subtype) | R |
| | images dataset | 500 | pixels averages, Fourier coefficients | | Unsup. Clust. (digits clust.) | |
| | Synthetic data | 300 | 2 omics | | Unsup. Clust. | |
| | Synthetic data | 300 | 3 omics | | Unsup. Clust. | |

[1] MOFA+ Argelaguet et al. [2020] has not been applied on patient's data but on single cell data. However, it is an extention of MOFA Argelaguet et al. [2018] which was tested on chronic lymphocytic leukaemia samples.

[2] DFMF Žitnik and Zupan [2014] has not been applied on patients' data. It could be however easily adapted to investigations of patients' data.

**Abbreviations**

**BioGRID**: Biological General Repository for Interaction Datasets; **CNV**: Copy Number Variation; **DisGeNET**: Disease-Gene Network; **GO**: Gene Ontology; **KEGG**: Kyoto Encyclopedia of Genes and Genomes; **MeSH**: Medical Subject Headings; **methy**: DNA methylation; **miRNA**: micro RNA; **mRNA**: messenger RNA; **MTF**: Matrix Tri-Factorization; **NMF**: Non-negative Matrix Factorization; **PMID**: PubMed IDentifier; **PSN**: Patient Similarity Network; **Pt**: Patient; **Sup. Class.**: Supervised Classification; **TCGA+*cancer code***: The Cancer Genome Atlas+ *link to complete cancer codes*; **Unsup. Clust.**: Unsupervised Clustering.

To better understand the process, the matrix multiplications can be rewritten to consider what happens at a single element $\mathbf{P}(i, j)$:

$$\mathbf{P}^{(1)}_{t+1}(i,j) = \sum_{k \in N_i} \sum_{l \in N_j} \mathbf{S}^{(1)}(i,k)\mathbf{P}^{(2)}_t(k,l)\mathbf{S}^{(1)}(j,l) \tag{S6}$$

that is similarity information is propagated across different networks only through common neighborhoods (Figure S7). In other words, $\mathbf{P}^{(1)}_{t+1}(i,j)$ is incremented only when there is an edge in $\mathbf{P}^{(2)}_t$ that connects the neighborhood of $x_i$ and $x_j$ (or that connects directly $x_i$ and $x_j$ since $N_i$ and $N_j$ include respectively $x_i$ and $x_j$).

SNF can be easily extended to $m > 2$ data sources:

$$\mathbf{P}^{(s)} = \mathbf{S}^{(s)} \frac{\sum_{k \neq s} \mathbf{P}^{(k)}}{m-1} \left(\mathbf{S}^{(s)\intercal}\right) \tag{S7}$$

and the final "consensus" matrix $\mathbf{P}^{(c)}$ is:

$$\mathbf{P}^{(c)} = \frac{1}{m} \sum_{k=1}^{m} \mathbf{P}^{(k)} \tag{S8}$$

**Table S2.** Data integration methods for partial datasets. For each method, the table reports: the name/acronym with the corresponding reference paper; the dataset used to develop and evaluate the approach in the reference paper and the corresponding sample cardinality and data types composing the dataset; the exploited integration method; the application task and the code availability (with link to the repository and programming languages for which the code is available).

| Name | Dataset | Sample Cardinality | Data type | Integration approach | Task | Code |
|---|---|---|---|---|---|---|
| SUMO Sienkiewicz et al. [2022] | 10 TCGA datasets | 3168 across datasets | mRNA miRNA methy | Cons. Clust. | Unsup. Clust. (Pt's subtype) | Python |
| | Synthetic data | 200 | 2 views | | Unsup. Clust. | |
| | METABRIC | | mRNA, CNV | | Unsup. Clust. | |
| | 34 TCGA datasets | | mRNA, methy, miRNA (32/34) | | Unsup. Clust. | |
| MONET Rappoport et al. [2020] | Synthetic data | 300 | 2 omics | Optimized Cons. clust. | Unsup. Clust. | Python |
| | Synthetic data | 150 | 3 omics | | Unsup. Clust. | |
| | images data | 400 | 6 views | | Unsup. Clust. (digits clust.) | |
| | 10 TCGA datasets | 3168 across datsets | mRNA miRNA methy | | Unsup. Clust. (Pt's subtype, gene clust.) | |
| | mouse embryos | 619 cells | mRNA methy | | Unsup. Clust. (cell clust.) | |
| DeepIMV Lee and van der Schaar [2021] | 38 TCGA datasets | 7295 | mRNA methy miRNA RPPA | Deep learning | Sup. Class. (1-year mortality) | Python |
| | CCLE | 504 | CNV methy mRNA miRNA RPPA metabolites | | Sup. Class. (drug sensitivity) | |
| MvNE Mitra et al. [2020] | 10 TCGA datasets | 3273 across datasets | mRNA miRNA methy | Probabilistic embedding | Unsup. Clust. (Pt's type and subtype, cancer stages) | None |
| | US Postal Service | 2000 images | Fourier coefficients of shapes, profile correlations | | Unsup. Clust. (digits) | |
| | Cora Dataset | 2708 documents | content, citations | | Unsup. Clust. (documents classes) | |
| | BBC corpora (synthetic) | 2012 documents | Segment representations | | Unsup. Clust. (topic classes) | |
| | VOC 2007 | 5619 images | GIST features tag features | | Unsup. Clust. (objects recognition) | |
| MOGDx Ryan et al. [2024] | TCGA BRCA TCGA LGG TCGA KIPAN | 1083 457 888 | mRNA, miRNA, methy, RPPA, CNV | GCN | Sup. class. (Pt's subtype, grade, type) | R+Python |

**Abbreviations**
**CCLE**: Cancer Cell Line Encyclopedia **CNV**: Copy Number Variation; **Cons. Clust.**: Consensus Clustering; **GCN**: Graph Convolutional Network; **METABRIC**: Molecular Taxonomy of Breast Cancer International Consortium; **methy**: DNA methylation; **miRNA**: micro RNA; **mRNA**: messenger RNA; **Pt**: Patient; **RPPA**: Reverse Phase Protein Array; **Sup. Class.**: Supervised Classification; **TCGA+*cancer code***: The Cancer Genome Atlas+ *link to complete cancer codes*; **Unsup. Clust.**: Unsupervised Clustering. **VOC 2007**: PASCAL Visual Object Classes Challenge 2007.

### S3. Dataset preparation

The TCGA repository groups cancers by their type (see [Tomczak et al., 2015] and the list of cancer types in the dedicated web page), but often clusters together samples having different histological types, which may add dispersive information biasing any analysis.

To achieve the highest possible biological homogeneity and ensure alignment with the latest WHO classifications, we carefully selected clinical-pathological features and filtered the histotypes included in TCGA for each cancer category.

Since the TCGA datasets were compiled prior to 2018, we retained only those histotypes from each dataset that hold clinical, histopathological, and biological significance according to the most recent WHO classifications. These classifications were published in 2018 for skin tumors (including SKCM), 2019 for breast tumors (including BRCA), 2020 for female genital tumors (including OV), 2021 for thoracic tumors (including LUAD and LUSC), and 2022 for urinary and male genital tumors (including BLCA, KIRC, and PRAD) [Elder et al., 2018, Tan et al., 2020, Lokuhetty et al., 2020, Tsao et al., 2022, Netto et al., 2022]

To improve the homogeneicity in the considered datasets, for the following cancers, we selected a subset of the available histological types:

- BLCA (Bladder Urothelial Carcinoma): only “muscle invasive urothelial carcinoma (pT2 or above)” samples were considered.
- BRCA (Breast invasive carcinoma): this dataset was split into two separated datasets (BRCA1 and BRCA2) comprising “infiltrating ductal carcinoma” and “infiltrating lobular carcinoma”, respectively.

- LUAD (Lung adenocarcinoma): only "lung acinar adenocarcinoma", "lung adenocarcinoma mixed subtype", "lung adenocarcinoma-not otherwise specified (NOS)", "lung bronchioloalveolar carcinoma mucinous", "lung bronchioloalveolar carcinoma nonmucinous", "lung micropapillary adenocarcinoma" and "lung papillary adenocarcinoma" were considered.
- LUSC (Lung squamous cell carcinoma): only "lung squamous cell carcinoma - not otherwise specified (NOS)" were considered.
- PRAD (Prostate adenocarcinoma): only "prostate adenocarcinoma acinar type" samples were considered.

The Kidney renal clear cell carcinoma (KIRC) dataset, the Ovarian serous cystadenocarcinoma (OV), and the Skin Cutaneous Melanoma dataset (SKCM) contained cancers belonging to the same histological type. We selectively excluded in each dataset the rare and unusual cancer histotypes, to only consider the most frequent and clinically relevant entities according to the WHO classifications and NIH's SEER database [Surveillance Research Program, 2023].

For Breast Invasive Carcinoma (BRCA), we considered only female patients, as male invasive breast cancer has recently emerged as a biologically and prognostically different tumor [Deb et al., 2012]. PRAD and OV datasets only included male and female patients respectively, for anatomical reasons. According to the same general principles of homogeneicity, only primary solid tumors are exploited for all cancer types except Skin Cutaneous Melanoma (SKCM). Most of the patients in SKCM dataset were metastatic (following the rapid progression of the disease), thus we retained both primary and metastatic samples, and when both are available we chosen the primary. We controlled for the absence of technical replicates in the datasets, and we further selected only samples not stored using "formalin-fixed paraffin-embedded" (FFPE). Indeed, this technique leads to lower quality nucleic acids (DNA and RNA) as they are fragmented and chemically modified by formalyn and lead to inferior outcomes in terms of quantity and quality of the molecular data compared to fresh samples [Hedegaard et al., 2014, Ottestad et al., 2022]. miRNA, mRNA, protein expression and DNA methylation data were downloaded for each cancer type (not including features having missing values). In particular, the following assays were exploited for each data view:

- *miRNA*: Gene-level log2 RPM expression values from RNA-Sequencing
- *mRNA*: RSEM TPM gene expression values from RNA-Sequencing
- *protein*: normalized protein expression values from Reverse Phase Protein Array (RPPA)
- *DNA methylation*: Probe-level methylation beta values from Methylation Array.

In the case of DNA methylation, we used data coming from "Infinium HumanMethylation 450K BeadChip" in all cases except for the Ovarian serous cystadenocarcinoma (OV) dataset, where only few samples where available for this array; thus, we opted for data with lower resolution coming from "Illumina HumanMethylation 27K BeadChip".

Only samples having a corresponding OS event and PFI event from TCGA-CDR dataset [Liu et al., 2018] were retained in each cancer dataset.

After downloading all the datasets we generated each view by applying the following filters, which allow to prune variables carrying practically no information.

1. **Remove variables mainly due to noise by filtering near-zero variance features**: Features with near zero variance were defined as: (1) features with very few unique values with respect to sample cardinality (in our experiments we removed features that had less than $\frac{N}{10}$ unique values, being $N$ the number of cases in the dataset); (2) features for which the frequency of the most common value is twenty times larger than the frequency of the second most common value.[2]
2. **Remove high pairwise-feature correlations**. Pairs of features showing a high pairwise correlation (i.e. Pearson correlation $> 0.75$) were filtered to remove the features having the highest mean correlation with all the other features in the dataset[3]. The parallel algorithm we implemented to perform this task for data views having more that 50000 features is outlined in Supplementary Section S3.1, otherwise the non parallelized algorithm provided by R function *caret::findCorrelation* was applied.

For the experiments involving the evaluation of clustering results by enrichment of clinical variables, we downloaded a wider set of relevant clinical variables, whose details are provided in Section S3.2.

### S3.1. High-pairwise correlation analysis and filtering

We considered as highly-correlated pairs of features with a Pearson correlation coefficient $corr_P > 0.75$. For each pair of correlated variables, we discarded the one whose mean absolute pairwise-correlation with respect to all the other features is higher. Since the application of pairwise-correlation filtering is computationally demanding, we implemented a parallel-distributed algorithm that applies the following consecutive steps.

1. The feature-set is split into non-intersecting feature sub-subsets; in other words, given the input dataset $\mathbf{X} \in \Re^{N \times D}$, where $N$ is the number of cases and $D$ is the number of features, we create $M$ non intersecting $d$-dimensional feature-subsets $\mathbf{X}^j_{sub} \in \Re^{N \times d}$, $\bigcap_{j=1,\dots,M} \mathbf{X}^j_{sub} = \emptyset$. The value of $d$ was set to $d = 5000$ to obtain practical computational costs.

[2] The function "nearZeroVar" from R package caret was used with default arguments.

[3] From a biological perspective, the correlation between omics variables could be meaningful. However, from a statistical point of view, highly correlated variables lead to inflated estimates, therefore affecting the reliability of statistical estimators. Thus, their removal is often advisable.

2. Each $\mathbf{X}^j_{sub}$ is assigned to a core, where it is processed to remove high-pairwise correlations, therefore obtaining a filtered subset $\mathbf{X}^j_{subF}$.
3. All the filtered subsets are recollected by the main thread and are recomposed by concatenation along the feature-dimension to obtain a new (partially filtered) dataset $\mathbf{X}_F = \left[\mathbf{X}^1_{subF}, \ldots, \mathbf{X}^M_{subF}\right]$.
4. The position of features in $\mathbf{X}_F$ is shuffled and the process (points 1 to 3) is iterated on $\mathbf{X}_F$ until a maximum number of iterations (set to 10) has been reached or no high-correlated features have been removed in any of the subsets.

### S3.2. Clinical variables for miss-SNF

For each cancer category we considered a set of clinical pathological data, including:

1. epidemiological data (e.g. sex, age at diagnosis, ethnicity);
2. data concerning risk factors or predictive markers of a certain type of cancers (e.g. menopause status at diagnosis for BRCA, tobacco smoking history for LUAD and LUSC, PSA values for PRAD [Bandi et al., 2021]);
3. histopathological data indicating the extent or clinical aggressiveness of cancer (e.g. TMN staging system [Amin et al., 2016], biological profile for BRCA [Harbeck et al., 2019], and Gleason score for PRAD [Netto et al., 2022]);
4. follow-up intervals and outcomes (OS and PFI, in days from diagnosis to last follow-up or patients' death) [Liu et al., 2018].

We provide the complete list of clinical variables that were selected as particularly relevant for our studies, while Table S3 shows useful descriptive statistics:

- **BLCA**: gender, age at initial diagnosis, M stage, N stage, T stage, pathologic stage, race, ethnicity, radiation therapy, tobacco smoking history
- **BRCA1, BRCA2**: age at initial diagnosis, M stage, N stage, T stage, pathologic stage, race, ethnicity, radiation therapy, margin status, estrogen receptor status, progesterone receptor status, HER2/neu receptor status, menopause status
- **KIRC**: gender, age at initial diagnosis, M stage, N stage, T stage, pathologic stage, race, ethnicity, history of prior malignancy
- **LUAD**: gender, age at initial diagnosis, M stage, N stage, T stage, pathologic stage, race, radiation therapy, tobacco smoking history, history of prior malignancy
- **LUSC**: gender, age at initial diagnosis, M stage, N stage, T stage, pathologic stage, race, radiation therapy, number pack years smoked, tobacco smoking history, history of prior malignancy, residual tumor
- **OV**: age at initial diagnosis, race, clinical stage
- **PRAD**: age at initial diagnosis, N stage, T stage, radiation therapy, history of prior malignancy, residual tumor, gleason score, psa value, psa result preop
- **SKCM**: gender, age at initial diagnosis, M stage, N stage, T stage, pathologic stage, race, ethnicity, radiation therapy

Of note, the choice of the selected clinical variables is driven by their clinical relevance with respect to the considered tumor. However, not all interesting variables are available for each tumor type or they can be available but having many missing values. In particular, we selected variables having less than 20% of missing values.

Table S3: Statistics for clinical variables. For each cancer we report the clinical variables with their total number and percentage (column “Total N (%)”). Column “Levels” reports variable categories if the variable is categorical. Column “Count (%)” shows the cardinality of each category and its percentage for categorical variables, while for numeric variables we report their median and interquartile range. (continue...)

| | Total N (%) | Levels | Count (%) |
|---|---|---|---|
| **BLCA** | 335 (100.0) | | |
| **gender** | 335 (100.0) | female | 84 (25.1) |
| | | male | 251 (74.9) |
| **age at initial diagnosis** | 335 (100.0) | Median (IQR) | 69.0 (60.0 to 76.0) |
| **pathology T stage** | 310 (92.5) | t0 | 1 (0.3) |
| | | t2 | 29 (9.4) |
| | | t2a | 22 (7.1) |
| | | t2b | 45 (14.5) |
| | | t3 | 39 (12.6) |
| | | t3a | 56 (18.1) |
| | | t3b | 70 (22.6) |
| | | t4 | 9 (2.9) |
| | | t4a | 34 (11.0) |
| | | t4b | 4 (1.3) |
| | | tx | 1 (0.3) |
| **pathology N stage** | 332 (99.1) | n0 | 197 (59.3) |
| | | n1 | 37 (11.1) |
| | | n2 | 65 (19.6) |
| | | n3 | 7 (2.1) |
| | | nx | 26 (7.8) |
| **pathology M stage** | 333 (99.4) | m0 | 160 (48.0) |
| | | m1 | 10 (3.0) |
| | | mx | 163 (48.9) |
| **pathologic stage** | 333 (99.4) | stage ii | 103 (30.9) |
| | | stage iii | 117 (35.1) |
| | | stage iv | 113 (33.9) |
| **race** | 320 (95.5) | asian | 39 (12.2) |
| | | black or african american | 20 (6.2) |
| | | white | 261 (81.6) |
| **ethnicity** | 304 (90.7) | hispanic or latino | 7 (2.3) |
| | | not hispanic or latino | 297 (97.7) |
| **radiation therapy** | 311 (92.8) | no | 297 (95.5) |
| | | yes | 14 (4.5) |
| **tobacco smoking history** | 322 (96.1) | current reformed smoker for <or = 15 years | 59 (18.3) |
| | | current reformed smoker for >15 years | 84 (26.1) |
| | | current reformed smoker, duration not specified | 11 (3.4) |
| | | current smoker | 74 (23.0) |
| | | lifelong non-smoker | 94 (29.2) |
| **OS** | 335 (100.0) | 0 | 184 (54.9) |
| | | 1 | 151 (45.1) |
| **OS time** | 334 (99.7) | Median (IQR) | 537.5 (334.0 to 935.2) |

Table S3: Statistics for clinical variables. For each cancer we report the clinical variables with their total number and percentage (column "Total N (%)"). Column "Levels" reports variable categories if the variable is categorical. Column "Count (%)" shows the cardinality of each category and its percentage for categorical variables, while for numeric variables we report their median and interquartile range. (continued)

| | Total N (%) | Levels | Count (%) |
|---|---|---|---|
| **BRCA1** | | | |
| **age at initial diagnosis** | 317 (100.0) | Median (IQR) | 56.0 (47.0 to 65.0) |
| **pathology T stage** | 317 (100.0) | t1 | 21 (6.6) |
| | | t1a | 1 (0.3) |
| | | t1b | 3 (0.9) |
| | | t1c | 61 (19.2) |
| | | t2 | 196 (61.8) |
| | | t3 | 25 (7.9) |
| | | t4 | 1 (0.3) |
| | | t4b | 7 (2.2) |
| | | t4d | 2 (0.6) |
| **pathology N stage** | 317 (100.0) | n0 | 110 (34.7) |
| | | n0 (i+) | 4 (1.3) |
| | | n0 (i-) | 29 (9.1) |
| | | n0 (mol+) | 1 (0.3) |
| | | n1 | 38 (12.0) |
| | | n1a | 50 (15.8) |
| | | n1b | 11 (3.5) |
| | | n1c | 2 (0.6) |
| | | n1mi | 11 (3.5) |
| | | n2 | 17 (5.4) |
| | | n2a | 23 (7.3) |
| | | n3 | 4 (1.3) |
| | | n3a | 9 (2.8) |
| | | n3b | 3 (0.9) |
| | | nx | 5 (1.6) |
| **pathology M stage** | 317 (100.0) | cm0 (i+) | 4 (1.3) |
| | | m0 | 243 (76.7) |
| | | m1 | 4 (1.3) |
| | | mx | 66 (20.8) |
| **pathologic stage** | 315 (99.4) | stage i | 24 (7.6) |
| | | stage ia | 27 (8.6) |
| | | stage ib | 1 (0.3) |
| | | stage ii | 2 (0.6) |
| | | stage iia | 110 (34.9) |
| | | stage iib | 78 (24.8) |
| | | stage iiia | 49 (15.6) |
| | | stage iiib | 5 (1.6) |
| | | stage iiic | 15 (4.8) |
| | | stage iv | 4 (1.3) |
| **race** | 316 (99.7) | asian | 21 (6.6) |
| | | black or african american | 91 (28.8) |
| | | white | 204 (64.6) |
| **ethnicity** | 295 (93.1) | hispanic or latino | 11 (3.7) |
| | | not hispanic or latino | 284 (96.3) |
| **radiation therapy** | 287 (90.5) | no | 124 (43.2) |
| | | yes | 163 (56.8) |
| **margin status** | 281 (88.6) | close | 6 (2.1) |
| | | negative | 259 (92.2) |
| | | positive | 16 (5.7) |
| **estrogen receptor status** | 287 (90.5) | negative | 88 (30.7) |
| | | positive | 199 (69.3) |
| **progesterone receptor status** | 286 (90.2) | indeterminate | 1 (0.3) |
| | | negative | 109 (38.1) |
| | | positive | 176 (61.5) |
| **HER2/neu receptor status** | 255 (80.4) | equivocal | 57 (22.4) |
| | | indeterminate | 6 (2.4) |
| | | negative | 151 (59.2) |
| | | positive | 41 (16.1) |
| **menopause status** | 289 (91.2) | peri (6-12 months since last menstrual period) | 12 (4.2) |
| | | post (prior bilateral ovariectomy or >12 mo since lmp with no prior hysterectomy) | 201 (69.6) |
| | | pre (<6 months since lmp and no prior bilateral ovariectomy and not on estrogen replacement) | 76 (26.3) |
| **OS** | 317 (100.0) | 0 | 275 (86.8) |
| | | 1 | 42 (13.2) |
| **OS time** | 317 (100.0) | Median (IQR) | 752.0 (470.0 to 1611.0) |

Table S3: Statistics for clinical variables. For each cancer we report the clinical variables with their total number and percentage (column "Total N (%)"). Column "Levels" reports variable categories if the variable is categorical. Column "Count (%)" shows the cardinality of each category and its percentage for categorical variables, while for numeric variables we report their median and interquartile range. (continued)

| | Total N (%) | Levels | Count (%) |
|---|---|---|---|
| **BRCA2** | 128 (100.0) | | |
| **age at initial diagnosis** | 128 (100.0) | Median (IQR) | 61.5 (50.0 to 70.0) |
| **pathology T stage** | 128 (100.0) | t1 | 4 (3.1) |
| | | t1b | 1 (0.8) |
| | | t1c | 14 (10.9) |
| | | t2 | 71 (55.5) |
| | | t2a | 1 (0.8) |
| | | t3 | 36 (28.1) |
| | | t4b | 1 (0.8) |
| **pathology N stage** | 128 (100.0) | n0 | 39 (30.5) |
| | | n0 (i+) | 6 (4.7) |
| | | n0 (i-) | 10 (7.8) |
| | | n1 | 14 (10.9) |
| | | n1a | 19 (14.8) |
| | | n1mi | 6 (4.7) |
| | | n2 | 4 (3.1) |
| | | n2a | 9 (7.0) |
| | | n3 | 9 (7.0) |
| | | n3a | 11 (8.6) |
| | | nx | 1 (0.8) |
| **pathology M stage** | 128 (100.0) | m0 | 88 (68.8) |
| | | m1 | 1 (0.8) |
| | | mx | 39 (30.5) |
| **pathologic stage** | 127 (99.2) | stage i | 6 (4.7) |
| | | stage ia | 6 (4.7) |
| | | stage ii | 1 (0.8) |
| | | stage iia | 33 (26.0) |
| | | stage iib | 35 (27.6) |
| | | stage iii | 2 (1.6) |
| | | stage iiia | 23 (18.1) |
| | | stage iiib | 1 (0.8) |
| | | stage iiic | 19 (15.0) |
| | | stage iv | 1 (0.8) |
| **race** | 126 (98.4) | asian | 8 (6.3) |
| | | black or african american | 10 (7.9) |
| | | white | 108 (85.7) |
| **ethnicity** | 124 (96.9) | hispanic or latino | 6 (4.8) |
| | | not hispanic or latino | 118 (95.2) |
| **radiation therapy** | 126 (98.4) | no | 52 (41.3) |
| | | yes | 74 (58.7) |
| **margin status** | 122 (95.3) | close | 2 (1.6) |
| | | negative | 111 (91.0) |
| | | positive | 9 (7.4) |
| **estrogen receptor status** | 124 (96.9) | negative | 5 (4.0) |
| | | positive | 119 (96.0) |
| **progesterone receptor status** | 124 (96.9) | indeterminate | 1 (0.8) |
| | | negative | 17 (13.7) |
| | | positive | 106 (85.5) |
| **HER2/neu receptor status** | 107 (83.6) | equivocal | 26 (24.3) |
| | | indeterminate | 3 (2.8) |
| | | negative | 60 (56.1) |
| | | positive | 18 (16.8) |
| **menopause status** | 120 (93.8) | indeterminate (neither pre or postmenopausal) | 1 (0.8) |
| | | peri (6-12 months since last menstrual period) | 5 (4.2) |
| | | post (prior bilateral ovariectomy or >12 mo since lmp with no prior hysterectomy) | 92 (76.7) |
| | | pre (<6 months since lmp and no prior bilateral ovariectomy and not on estrogen replacement) | 22 (18.3) |
| **OS** | 128 (100.0) | 0 | 114 (89.1) |
| | | 1 | 14 (10.9) |
| **OS time** | 128 (100.0) | Median (IQR) | 787.5 (502.0 to 1554.5) |

Table S3: Statistics for clinical variables. For each cancer we report the clinical variables with their total number and percentage (column "Total N (%)"). Column "Levels" reports variable categories if the variable is categorical. Column "Count (%)" shows the cardinality of each category and its percentage for categorical variables, while for numeric variables we report their median and interquartile range. (continued)

| | Total N (%) | Levels | Count (%) |
|---|---|---|---|
| **KIRC** | 169 (100.0) | | |
| **gender** | 169 (100.0) | female | 60 (35.5) |
| | | male | 109 (64.5) |
| **age at initial diagnosis** | 169 (100.0) | Median (IQR) | 61.0 (52.0 to 69.0) |
| **pathology T stage** | 169 (100.0) | t1 | 10 (5.9) |
| | | t1a | 36 (21.3) |
| | | t1b | 39 (23.1) |
| | | t2 | 18 (10.7) |
| | | t2a | 6 (3.6) |
| | | t2b | 1 (0.6) |
| | | t3 | 4 (2.4) |
| | | t3a | 30 (17.8) |
| | | t3b | 21 (12.4) |
| | | t4 | 4 (2.4) |
| **pathology N stage** | 169 (100.0) | n0 | 71 (42.0) |
| | | n1 | 4 (2.4) |
| | | nx | 94 (55.6) |
| **pathology M stage** | 167 (98.8) | m0 | 121 (72.5) |
| | | m1 | 27 (16.2) |
| | | mx | 19 (11.4) |
| **pathologic stage** | 168 (99.4) | stage i | 82 (48.8) |
| | | stage ii | 21 (12.5) |
| | | stage iii | 32 (19.0) |
| | | stage iv | 33 (19.6) |
| **race** | 166 (98.2) | asian | 1 (0.6) |
| | | black or african american | 33 (19.9) |
| | | white | 132 (79.5) |
| **ethnicity** | 146 (86.4) | hispanic or latino | 6 (4.1) |
| | | not hispanic or latino | 140 (95.9) |
| **history of prior malignancy** | 164 (97.0) | no | 145 (88.4) |
| | | yes, history of prior malignancy | 17 (10.4) |
| | | yes, history of synchronous and or bilateral malignancy | 2 (1.2) |
| **OS** | 169 (100.0) | 0 | 121 (71.6) |
| | | 1 | 48 (28.4) |
| **OS time** | 169 (100.0) | Median (IQR) | 1130.0 (469.0 to 2271.0) |
| **LUAD** | 300 (100.0) | | |
| **gender** | 300 (100.0) | female | 164 (54.7) |
| | | male | 136 (45.3) |
| **age at initial diagnosis** | 286 (95.3) | Median (IQR) | 65.0 (59.0 to 72.0) |
| **pathology T stage** | 300 (100.0) | t1 | 42 (14.0) |
| | | t1a | 27 (9.0) |
| | | t1b | 30 (10.0) |
| | | t2 | 105 (35.0) |
| | | t2a | 43 (14.3) |
| | | t2b | 18 (6.0) |
| | | t3 | 20 (6.7) |
| | | t4 | 14 (4.7) |
| | | tx | 1 (0.3) |
| **pathology N stage** | 300 (100.0) | n0 | 188 (62.7) |
| | | n1 | 57 (19.0) |
| | | n2 | 48 (16.0) |
| | | nx | 7 (2.3) |
| **pathology M stage** | 297 (99.0) | m0 | 201 (67.7) |
| | | m1 | 8 (2.7) |
| | | m1a | 1 (0.3) |
| | | m1b | 3 (1.0) |
| | | mx | 84 (28.3) |
| **pathologic stage** | 297 (99.0) | stage i | 2 (0.7) |
| | | stage ia | 78 (26.3) |
| | | stage ib | 79 (26.6) |
| | | stage ii | 1 (0.3) |
| | | stage iia | 31 (10.4) |
| | | stage iib | 41 (13.8) |
| | | stage iiia | 46 (15.5) |
| | | stage iiib | 7 (2.4) |
| | | stage iv | 12 (4.0) |
| **race** | 268 (89.3) | asian | 4 (1.5) |
| | | black or african american | 36 (13.4) |
| | | white | 228 (85.1) |
| **radiation therapy** | 279 (93.0) | no | 237 (84.9) |
| | | yes | 42 (15.1) |
| **tobacco smoking history** | 290 (96.7) | current reformed smoker for <or = 15 years | 100 (34.5) |
| | | current reformed smoker for >15 years | 80 (27.6) |
| | | current reformed smoker, duration not specified | 1 (0.3) |
| | | current smoker | 64 (22.1) |
| | | lifelong non-smoker | 45 (15.5) |
| **history of prior malignancy** | 298 (99.3) | no | 252 (84.6) |
| | | yes, history of prior malignancy | 36 (12.1) |
| | | yes, history of synchronous and or bilateral malignancy | 10 (3.4) |
| **OS** | 300 (100.0) | 0 | 180 (60.0) |
| | | 1 | 120 (40.0) |
| **OS time** | 292 (97.3) | Median (IQR) | 661.0 (412.8 to 1159.5) |

Table S3: Statistics for clinical variables. For each cancer we report the clinical variables with their total number and percentage (column “Total N (%)”). Column “Levels” reports variable categories if the variable is categorical. Column “Count (%)” shows the cardinality of each category and its percentage for categorical variables, while for numeric variables we report their median and interquartile range. (continued)

| | Total N (%) | Levels | Count (%) |
|---|---|---|---|
| **LUSC** | 228 (100.0) | | |
| **gender** | 228 (100.0) | female | 53 (23.2) |
| | | male | 175 (76.8) |
| **age at initial diagnosis** | 223 (97.8) | Median (IQR) | 68.0 (62.0 to 74.0) |
| **pathology T stage** | 228 (100.0) | t1 | 18 (7.9) |
| | | t1a | 9 (3.9) |
| | | t1b | 25 (11.0) |
| | | t2 | 53 (23.2) |
| | | t2a | 54 (23.7) |
| | | t2b | 20 (8.8) |
| | | t3 | 41 (18.0) |
| | | t4 | 8 (3.5) |
| **pathology N stage** | 228 (100.0) | n0 | 143 (62.7) |
| | | n1 | 63 (27.6) |
| | | n2 | 18 (7.9) |
| | | nx | 4 (1.8) |
| **pathology M stage** | 228 (100.0) | m0 | 183 (80.3) |
| | | m1 | 1 (0.4) |
| | | m1a | 1 (0.4) |
| | | m1b | 1 (0.4) |
| | | mx | 42 (18.4) |
| **pathologic stage** | 226 (99.1) | stage i | 2 (0.9) |
| | | stage ia | 41 (18.1) |
| | | stage ib | 60 (26.5) |
| | | stage ii | 3 (1.3) |
| | | stage iia | 36 (15.9) |
| | | stage iib | 46 (20.4) |
| | | stage iii | 2 (0.9) |
| | | stage iiia | 29 (12.8) |
| | | stage iiib | 4 (1.8) |
| | | stage iv | 3 (1.3) |
| **race** | 194 (85.1) | asian | 4 (2.1) |
| | | black or african american | 15 (7.7) |
| | | white | 175 (90.2) |
| **radiation therapy** | 218 (95.6) | no | 190 (87.2) |
| | | yes | 28 (12.8) |
| **tobacco smoking history** | 219 (96.1) | current reformed smoker for <or = 15 years | 101 (46.1) |
| | | current reformed smoker for >15 years | 35 (16.0) |
| | | current reformed smoker, duration not specified | 3 (1.4) |
| | | current smoker | 72 (32.9) |
| | | lifelong non-smoker | 8 (3.7) |
| **history of prior malignancy** | 228 (100.0) | no | 204 (89.5) |
| | | yes, history of prior malignancy | 21 (9.2) |
| | | yes, history of synchronous and or bilateral malignancy | 3 (1.3) |
| **number pack years smoked** | 191 (83.8) | Median (IQR) | 44.0 (30.0 to 55.9) |
| **residual tumor** | 185 (81.1) | r0 | 170 (91.9) |
| | | r1 | 5 (2.7) |
| | | r2 | 2 (1.1) |
| | | rx | 8 (4.3) |
| **OS** | 228 (100.0) | 0 | 135 (59.2) |
| | | 1 | 93 (40.8) |
| **OS time** | 225 (98.7) | Median (IQR) | 688.0 (377.0 to 1336.0) |
| **OV** | 226 (100.0) | | |
| **age at initial diagnosis** | 226 (100.0) | Median (IQR) | 58.0 (51.0 to 67.0) |
| **race** | 216 (95.6) | american indian or alaska native | 2 (0.9) |
| | | asian | 10 (4.6) |
| | | black or african american | 13 (6.0) |
| | | white | 191 (88.4) |
| **clinical stage** | 225 (99.6) | stage iia | 1 (0.4) |
| | | stage iib | 3 (1.3) |
| | | stage iic | 10 (4.4) |
| | | stage iiia | 6 (2.7) |
| | | stage iiib | 10 (4.4) |
| | | stage iiic | 169 (75.1) |
| | | stage iv | 26 (11.6) |
| **OS** | 226 (100.0) | 0 | 83 (36.7) |
| | | 1 | 143 (63.3) |
| **OS time** | 225 (99.6) | Median (IQR) | 1039.0 (518.0 to 1650.0) |

Table S3: Statistics for clinical variables. For each cancer we report the clinical variables with their total number and percentage (column "Total N (%)"). Column "Levels" reports variable categories if the variable is categorical. Column "Count (%)" shows the cardinality of each category and its percentage for categorical variables, while for numeric variables we report their median and interquartile range. (continued)

| | Total N (%) | Levels | Count (%) |
|---|---|---|---|
| **PRAD** | 337 (100.0) | | |
| **age at initial diagnosis** | 337 (100.0) | Median (IQR) | 62.0 (57.0 to 66.0) |
| **pathology T stage** | 332 (98.5) | t2a | 6 (1.8) |
| | | t2b | 7 (2.1) |
| | | t2c | 95 (28.6) |
| | | t3a | 118 (35.5) |
| | | t3b | 97 (29.2) |
| | | t4 | 9 (2.7) |
| **pathology N stage** | 296 (87.8) | n0 | 235 (79.4) |
| | | n1 | 61 (20.6) |
| **radiation therapy** | 311 (92.3) | no | 272 (87.5) |
| | | yes | 39 (12.5) |
| **history of prior malignancy** | 337 (100.0) | no | 315 (93.5) |
| | | yes, history of prior malignancy | 20 (5.9) |
| | | yes, history of synchronous and or bilateral malignancy | 2 (0.6) |
| **residual tumor** | 327 (97.0) | r0 | 210 (64.2) |
| | | r1 | 104 (31.8) |
| | | r2 | 3 (0.9) |
| | | rx | 10 (3.1) |
| **Gleason score** | 337 (100.0) | Median (IQR) | 7.0 (7.0 to 9.0) |
| **psa value** | 304 (90.2) | Median (IQR) | 0.1 (0.0 to 0.1) |
| **psa result preop** | 335 (99.4) | Median (IQR) | 7.7 (5.3 to 12.0) |
| **OS** | 337 (100.0) | 0 | 331 (98.2) |
| | | 1 | 6 (1.8) |
| **OS time** | 337 (100.0) | Median (IQR) | 992.0 (526.0 to 1518.0) |
| **SKCM** | 334 (100.0) | | |
| **gender** | 334 (100.0) | female | 138 (41.3) |
| | | male | 196 (58.7) |
| **age at initial diagnosis** | 328 (98.2) | Median (IQR) | 57.0 (48.0 to 71.2) |
| **pathology T stage** | 309 (92.5) | t0 | 19 (6.1) |
| | | t1 | 8 (2.6) |
| | | t1a | 17 (5.5) |
| | | t1b | 6 (1.9) |
| | | t2 | 18 (5.8) |
| | | t2a | 21 (6.8) |
| | | t2b | 10 (3.2) |
| | | t3 | 5 (1.6) |
| | | t3a | 23 (7.4) |
| | | t3b | 26 (8.4) |
| | | t4 | 12 (3.9) |
| | | t4a | 15 (4.9) |
| | | t4b | 89 (28.8) |
| | | tis | 1 (0.3) |
| | | tx | 39 (12.6) |
| **pathology N stage** | 317 (94.9) | n0 | 153 (48.3) |
| | | n1 | 12 (3.8) |
| | | n1a | 10 (3.2) |
| | | n1b | 28 (8.8) |
| | | n2 | 5 (1.6) |
| | | n2a | 10 (3.2) |
| | | n2b | 19 (6.0) |
| | | n2c | 9 (2.8) |
| | | n3 | 45 (14.2) |
| | | nx | 26 (8.2) |
| **pathology M stage** | 312 (93.4) | m0 | 290 (92.9) |
| | | m1 | 5 (1.6) |
| | | m1a | 4 (1.3) |
| | | m1b | 5 (1.6) |
| | | m1c | 8 (2.6) |
| **pathologic stage** | 299 (89.5) | i/ii nos | 12 (4.0) |
| | | stage i | 18 (6.0) |
| | | stage ia | 13 (4.3) |
| | | stage ib | 19 (6.4) |
| | | stage ii | 13 (4.3) |
| | | stage iia | 12 (4.0) |
| | | stage iib | 16 (5.4) |
| | | stage iic | 46 (15.4) |
| | | stage iii | 29 (9.7) |
| | | stage iiia | 9 (3.0) |
| | | stage iiib | 37 (12.4) |
| | | stage iiic | 54 (18.1) |
| | | stage iv | 21 (7.0) |
| **race** | 326 (97.6) | asian | 11 (3.4) |
| | | white | 315 (96.6) |
| **ethnicity** | 326 (97.6) | hispanic or latino | 7 (2.1) |
| | | not hispanic or latino | 319 (97.9) |
| **radiation therapy** | 334 (100.0) | no | 303 (90.7) |
| | | yes | 31 (9.3) |
| **OS** | 334 (100.0) | 0 | 184 (55.1) |
| | | 1 | 150 (44.9) |
| **OS time** | 327 (97.9) | Median (IQR) | 938.0 (445.5 to 2036.5) |

### S4. miss-SNF convergence experiments

The data integration performed by miss-SNF is based on a cross-diffusion process on the local similarities of several observations across different sources (Section 2). The cross-diffusion process iteratively passes information across different networks exploiting the presence of (local) neighbourhoods connected in the different data sources, which makes the "global" similarity matrices representing each data source progressively more similar until convergence. The stopping criterion implemented in SNF and miss-SNF regards the number of iterations to be performed, which is set *a priori*. Thus, it is important to find the appropriate value for the number of iterations $T$ for the diffusion process. SNF's authors recommend setting the number of iterations $T \in (10, 20)$ based on convergence experiments [Wang et al., 2014].

In this section, we experimentally assess the convergence and the reconstruction capabilities of miss-SNF with respect to the choice of the initialization approach for the missing information during the initial phase (i.e. one, zero, equidistant and random initializations; see Section 2). We confront the results of the method on several datasets when different proportions of data are amputated from the original dataset leading to "partial datasets".

Tests are carried out on nine datasets (BLCA, BRCA1, BRCA2, KIRC, LUAD, LUSC, OV, PRAD, SKCM) comprising four omics (miRNA, mRNA, proteins and DNA methylation). For each of them, we consider three (partial) versions that contain increasing proportions of missing entries (10%, 30% and 50%). Moreover, to capture the potential variability of the results, we generate ten versions of the partial datasets for each percentage of amputation where random patients are removed (see Section 3.1 for details about datasets).

For the convergence analysis (Section S4.1), we compare the *stability* of the integration process by measuring the variation of the integrated matrix between two consecutive interactions of the diffusion process. We also evaluate the convergence *velocity* by comparing the integrated matrix after each iteration with the final one, obtained at the end of the integration process (and considered as the best approximation of the true solution). The *reconstruction capability* (Section S4.2) is quantified by measuring the distance between the integration matrices obtained using partial and complete data respectively.

These analyses guide the choice of the parameters in the subsequent experimental evaluations reported in Section 3.3.

#### S4.1. Convergence analysis

To evaluate the convergence of the method, we execute miss-SNF on all the complete and amputated cancer datasets for a fixed number of iterations. For each execution, we assess the stability of the process by computing the Frobenius distance between the resulting integration matrices obtained in two consecutive iterations of the cross-diffusion. Additionally, we measure the velocity of the convergence by computing the Frobenius distance between the integration matrix after each iteration and the one obtained at the end of the process.

Figure S1 and Figure S2 depict the results when 100 iterations are considered for stability and velocity respectively. Each curve represents the mean value computed on the ten random amputations for each percentage of missingness, while shaded regions depict the min-max interval of the measured distance.

Miss-SNF *random* reaches convergence faster with respect to *zero* and *equi*, while *one* has the slowest convergence in terms of both stability and velocity. The proportion of missing information has diverse impacts on the different methods: for the *random* strategy there is not a noticeable effect, for *equi* and *zero* the convergence is quicker and more stable for higher percentages of missing data, while for *one*, higher percentages of missing data appear to be associated to less stable and slower convergence.

From a quantitative point of view, *random* strategy reaches convergence stability after 20 iterations and independently of the percentage of missing data; the *equi* strategy stabilizes after circa 20 and 30 iterations depending on the proportion of missing information; *zero* reaches convergence stability between 20 and 40 iterations, and *one* did not reach a stable regime after 100 iterations in almost all the datasets.

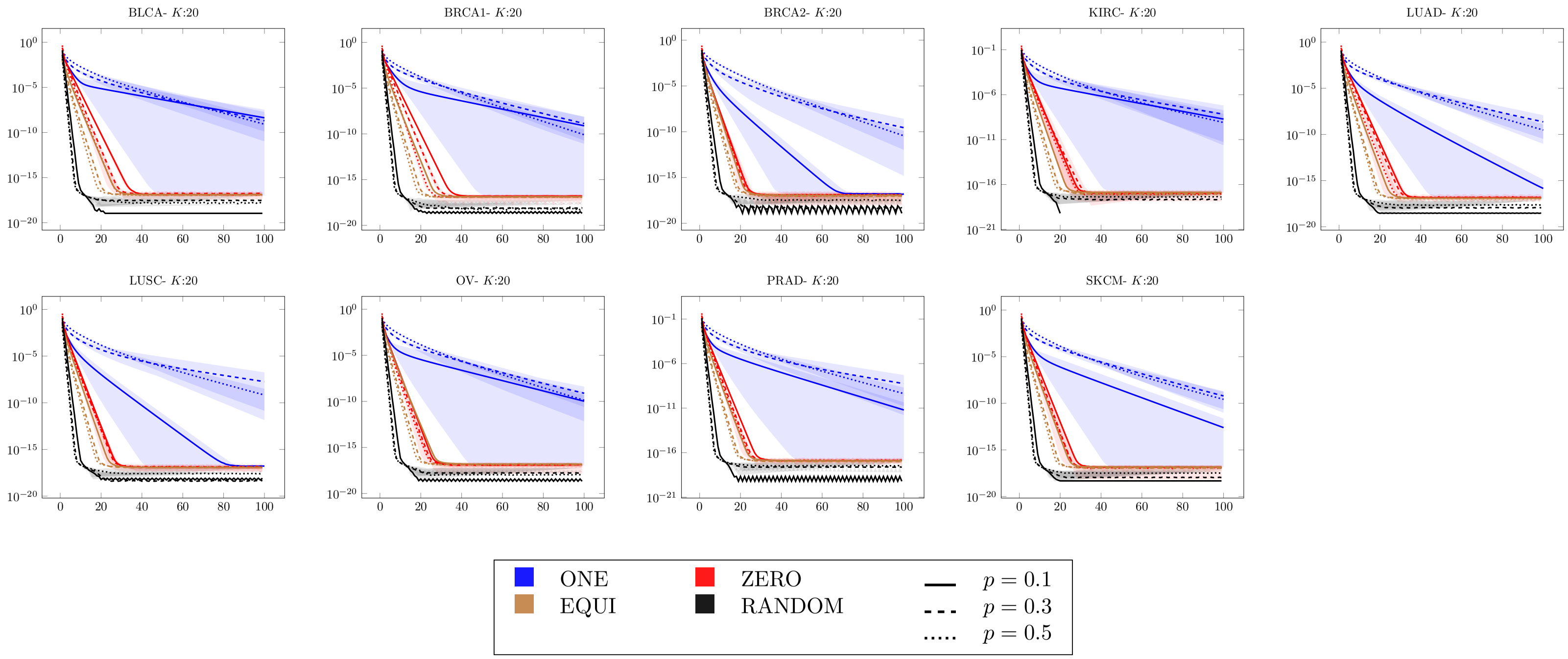


Fig. S1: Convergence stability evaluated using the Frobenius norm between the integrated matrices in consecutive steps of the algorithm for 100 iterations. Each panel shows a different dataset using a fixed neighbourhood size ($K = 20$). Lines depict the mean distance on ten random amputations for each dataset, where a more dashed line indicates a higher percentage of missing samples and the color is related to the different initializations of miss-SNF. Shadows represent the minimum and maximum values.

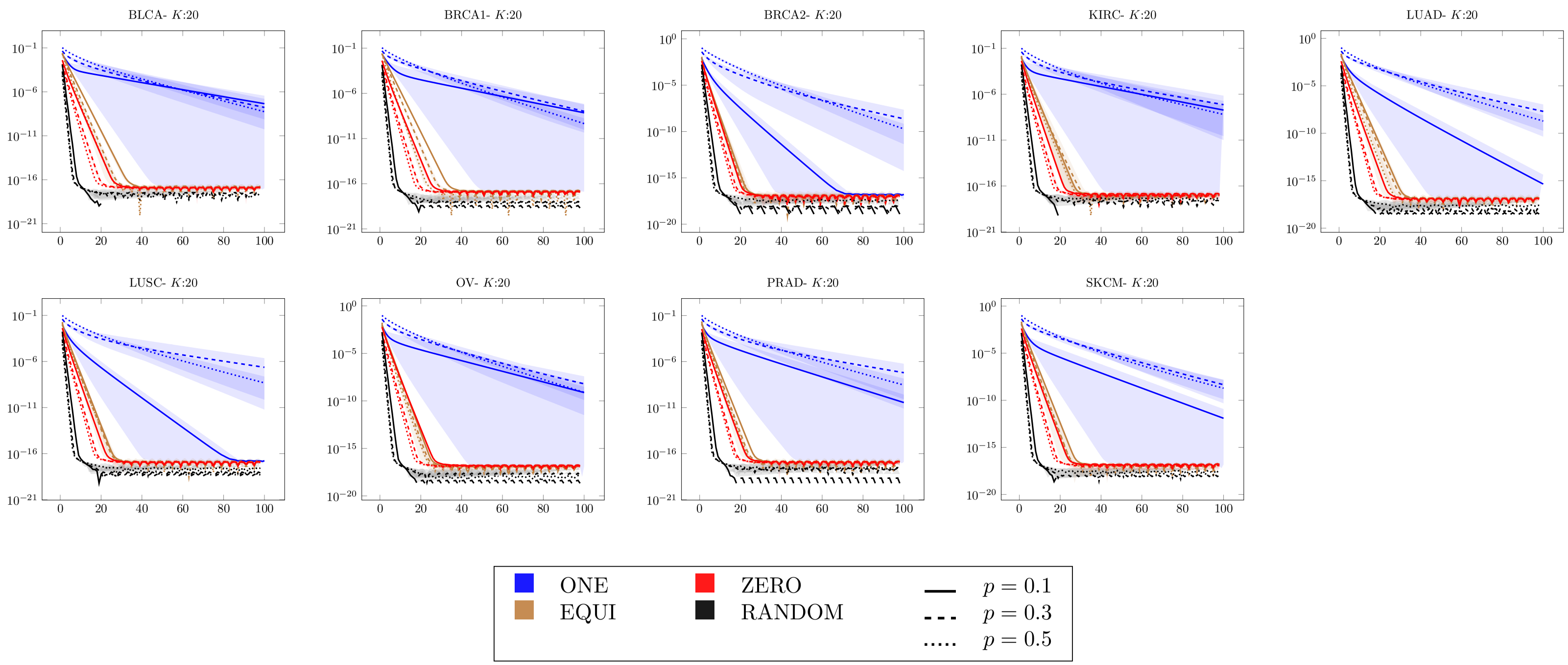


Fig. S2: Convergence velocity evaluated using the Frobenius norm between the integrated matrices in each step of the algorithm and the final one for 100 iterations. Each panel shows a different dataset using a fixed neighbourhood size ($K = 20$). Lines depict the mean distance on ten random amputations for each dataset, where a more dashed line indicates a higher percentage of missing samples and the color is related to the different initialization of miss-SNF. Shadows represent the minimum and maximum values.

#### S4.2. Partial data reconstruction error

To analyse the effect of missing information on miss-SNF, we execute the method on datasets with different proportions of missing data, and we compare the matrix generated by the method when complete and partial datasets are used. Specifically, for each cancer dataset we randomly generate ten amputated versions using increasing proportions of completely missing samples ranging from 10% to 50%. For each execution, we measure the reconstruction error as the 1-norm distance between the resulting integrated matrices obtained with the full and the amputated dataset.

Figure S3 shows the error distributions for the different initialization strategies. Among the different initialization options, *one* and *equidistant* are the ones having smaller reconstruction errors, while *random* results are similar independently of the missing data proportion. Moreover, *one* and *equidistant* have a less steep increase in error w.r.t. *zero* strategy. This could be considered a foreseeable behavior if we interpret *random* strategy as the injection of random noise in the data. On the other hand, *one* and *equidistant* are the methods that enable the partial reconstruction of missing pairwise similarities coming from the presence of partial observations by assigning a larger value to the self-similarity, differently from the *zero* strategy. Thus, *one* and *equidistant* show in general a lower reconstruction error across the different percentage of data amputations. In this set of experiments we used 100 iterations for all the methods but, as shown in Section S4.1, *one* did not reach convergence. Thus, in the main experiments presented in Section 3.3 we used 1000 iterations for miss-SNF *one*.

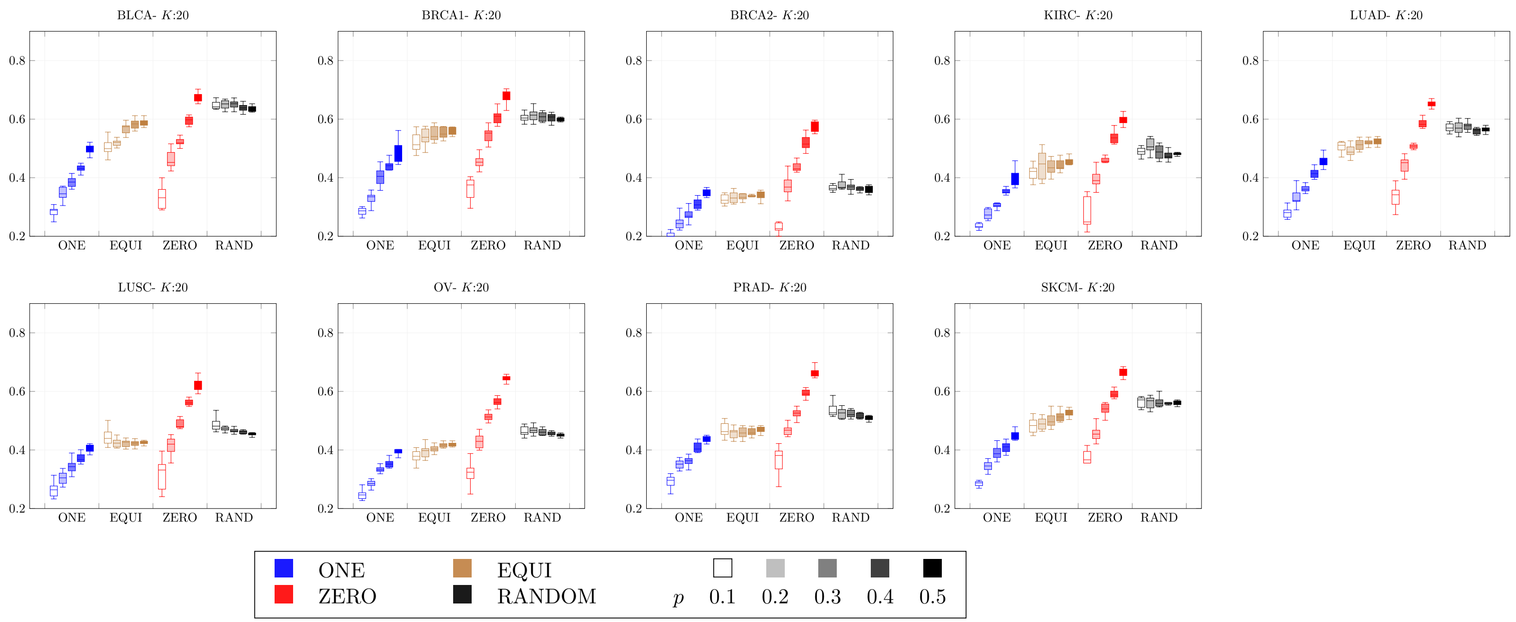


Fig. S3: Evaluation of miss-SNF data recovery in terms of 1-norm. Boxplots represent the distribution of the distance between the generated matrix using full and partial information. Each panel depicts a different dataset using a fixed neighbourhood size ($K = 20$). For each dataset, 10 random amputations were generated with increasing percentages of amputation ($p$=10%, $p$=20%, etc.) represented by the intensity of the color while the color itself is assigned to the different miss-SNF initialization strategies.

### S5. Statistical analysis

Paired samples Wilcoxon test, alias Wilcoxon signed-rank test, at the 95% of confidence (i.e. $\alpha = 0.05$) was used for comparison. If not specified, the test was performed by pooling the results obtained on all the nine TCGA datasets; for each comparison, and having defined a proper metric for evaluation, we exploited win-tie-loss tables to summarize the statistical comparison between each method against all the others. When using sided-hypothesis tests to compare the performance of two methods $A$ and $B$, where higher values of the chosen metric indicate better performance, a win is assigned if the $p - value < \alpha$ for the "greater" alternative, a loss is assigned if $p - value < \alpha$ for the "less" alternative, and a tie is assigned otherwise. When lower values of the metric indicate better performance (i.e., Normalized Variation of Information), a win is assigned if the $p - value < \alpha$ for the "less" alternative, a loss is assigned if the $p - value < \alpha$ for the "greater" alternative, and a tie is assigned otherwise. When assessing multiple methods, all pairwise comparisons are performed, and a three-column table is computed that lists, for each method, the number of wins, ties, and losses. In particular, when comparing data fusion+clustering algorithms, we paired the results obtained across the nine TCGA datasets, the missingness percentage and the $h = 10$ random repetitions at that percentage. When, instead, we performed more generic comparisons across data fusion experiments, we paired the results obtained across the nine different datasets, the missingness percentage, the $h = 10$ random repetitions at that percentage, and the clustering/classification algorithm. Win-tie-loss tables that summarize the results of enriched clinical variables and differential survival use only the results obtained from the amputated datasets since we compare miss-SNF against other data fusion approaches for the integration of partial datasets, while SNF is used to fuse complete datasets.

Wilcoxon signed-rank tests summarize and compare the performance of different pipelines across multiple settings. Therefore, pipelines that achieve the highest/lowest number of wins/losses can be regarded as being, on the average of all the experimented settings, the top-performing and most robust.

### S6. Pairwise similarity reconstruction in the fused PSNs: additional results

In this section we present additional results that complement the analysis reported in Section 3.3. Table S4 shows the win-tie-losses for each miss-SNF initialization option considering all cancers together, while Figure S4 presents the results for cancers separately. The win-tie-loss table confirms miss-SNF one as the best option to initialize missing similarities for partial samples, followed by equidistant method. Unsurprisingly, these are the two approaches designed to recover missing pairwise similarities during the integration process. On the other hand, ignoring partial samples in the data fusion process penalizes miss-SNF zero, which obtains the worst performance.

**Table S4.** Win-tie-loss table for miss-SNF initializations. One-sided paired Wilcoxon test ($\alpha = 0.05$) is exploited to compare the different miss-SNF versions in terms of 1-norm distance, considering all amputation percentages. A win is assigned when the reconstruction error is significantly less than another method, a loss is assigned in the opposite case, otherwise the methods tie.

| Method | wins | ties | losses |
|---|---|---|---|
| one | 3 | 0 | 0 |
| equi | 2 | 0 | 1 |
| random | 1 | 0 | 2 |
| zero | 0 | 0 | 3 |

For the sake of completeness, we provide the boxplots showing the 1-norm distribution at different amputation percentages for each cancer in Figure S5.

### S7. Clustering experiments

#### S7.1. Experimental setup for clustering analysis

miss-SNF is compared against representative state-of-the-art data fusion approaches able to integrate partial datasets: an input data-fusion approach based on matrix factorization, MOFA+ [Argelaguet et al., 2020], and a PSN-fusion method called NEMO [Rappoport and Shamir, 2019]. The integrated matrices computed by miss-SNF *one* and *equidistant*[4], NEMO and MOFA+ are used to integrate complete and amputated cancer datasets, where for each amputation percentage (i.e. 10%, 20%, 30%, 40%, 50%) we have 10 different randomly amputated datasets (details on datasets creation are available in Section 3.1). The hyper-parameters of MOFA+ and NEMO are set to their default values[5].

Once the matrices integrated using miss-SNF, MOFA+ and NEMO are computed for complete and amputated datasets, they are used as input to perform patients clustering by means of the following classic approaches: k-means [Hastie et al., 2009], Spectral Clustering

[4] All the experiments set $K = 20$ and $T = 100$ iterations for miss-SNF (except for miss-SNF one where $T = 1000$ due to its slow convergence, see Supplementary Section S4 for convergence experiments). We used the same values of $K$ and $T$ when comparing SNF and miss-SNF results.

[5] The only hyper-parameter required by NEMO is the number of neighbors to use for each omic, which is set by default as $k = NA$. This means that NEMO automatically selects the number of neighbours to be the number of samples available for each omic divided by 6, where 6 is an estimate of the number of clusters observed in cancer dataset [Rappoport and Shamir, 2019].

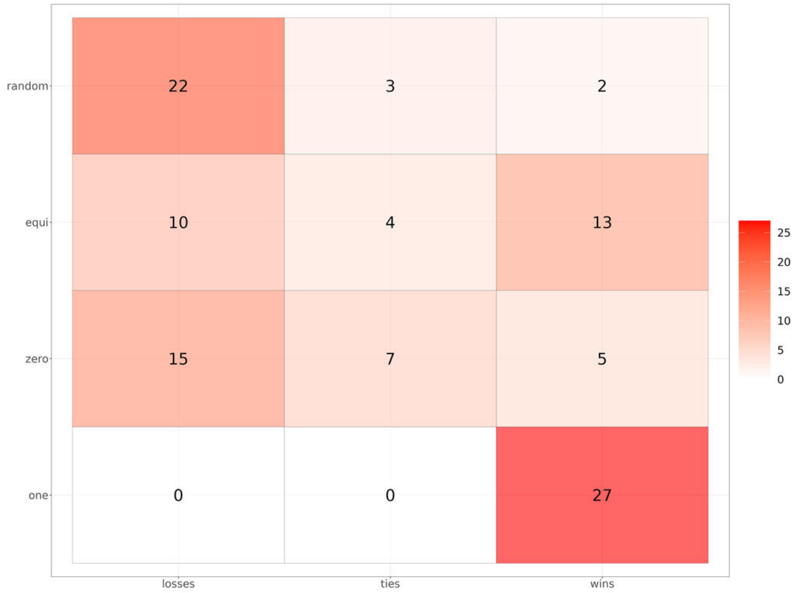


Fig. S4: Number of win-tie-losses (x-axis) for miss-SNF initializations (y-axis). Tests are computed using one-sided paired Wilcoxon test ($\alpha = 0.05$) considering the 1-norm values. Cancers are considered separately leading to 27 comparisons for each variant.

(SP) [Von Luxburg, 2007] and Partitioning Around Medoids (PAM) [Kaufman and Rousseeuw, 1990] (see Supplementary Section S7.2 for further details).

Clustering approaches are applied on the integrated matrices obtained by miss-SNF, NEMO and MOFA+. miss-SNF and NEMO works on PSN and they return a fused patient similarity network as output, which we used to run k-means, PAM and spectral clustering algorithms. Since k-means cannot take as input a similarity/distance matrix, the algorithm interprets the PSN as a conventional feature matrix (features are the similarities of a patient with all the others) and internally exploits it to compute distances by means of Euclidean distance. On the other hand, PAM can directly leverage a dissimilarity matrix, which is easily obtained from the integrated and normalized PSN as $\mathbf{D} = 1 - \mathbf{W}$. Finally, the integrated PSN obtained from miss-SNF and NEMO is directly used by SP approaches. SP algorithm has different variants and in our experiments we decided to evaluate the Ng et al. algorithm [Ng et al., 2001], as implemented in the *sClust* package. For the experiments with NEMO, we additionally evaluated the algorithm used in SNF and NEMO reference papers [Wang et al., 2014, Rappoport and Shamir, 2019], which is implemented in the R package *SNFtool*. MOFA+ does not exploit PSN during the integration process and it returns an integrated matrix $\mathbf{M} \in \mathbb{R}^{n \times f}$ having the set of samples on the rows and the computed factors on the columns. To perform experiments using MOFA+ as data integration method, we run k-means and PAM on the matrix of factors returned by MOFA+ where both the clustering approaches internally exploits Euclidean distance. Moreover, we computed from the continuous factors matrix a corresponding similarity matrix leveraging the *scaled exponential Euclidean kernel* [Wang et al., 2014] and we used this similarity matrix as input for k-means, PAM and SP, following the same experimental setup explained for miss-SNF. All similarity matrices were normalized using min-max graph normalization before using them to perform clustering. Considering all the experimented combinations of data integration approaches (i.e. miss-SNF *one*, miss-SNF*equidistant*, NEMO and MOFA+), clustering methods (i.e. k-means, PAM, Spectral clustering - two versions evaluated for NEMO) and the use of both factor matrices and similarity matrices for MOFA+ with k-means and PAM, we performed 15 different experiments for each of the 9 cancer datasets (for a total of 135 tests).

All the considered clustering approaches require to define *a priori* the number of clusters to look for in the dataset. In our experiments, we run all the clustering algorithms considering a number of clusters $nc = [2 .. 10]$. The selection of the optimal number of clusters is still an open problem and many methods are proposed in literature to tackle this issue (e.g. Elbow method [Bholowalia and Kumar, 2014], Silhouette method [Rousseeuw, 1987], Gap statistic [Tibshirani et al., 2001]). Moreover, data integration approaches that are evaluated on clustering tasks often suggest methods to select the best number of clusters [Duan et al., 2021] and the diversity in the proposed approaches remarks the difficulty of this problem. Following the work in [Duan et al., 2021], we initially evaluate clustering results considering all numbers of clusters to make our experiments exhaustive and less influenced by the choice of the method to detect the number of clusters. We compare clusterings from complete and amputated datasets, obtained with miss-SNF and its competitors, considering two different aspects:

1. *Similarity between clusterings* is evaluated through the use of state-of-the-art metrics comprising adjusted Rand Index, Normalized Mutual Information (NMI) and Normalized Variation of Information (NVI) [Wagner and Wagner, 2007, Lancichinetti et al., 2009, You, 2021].

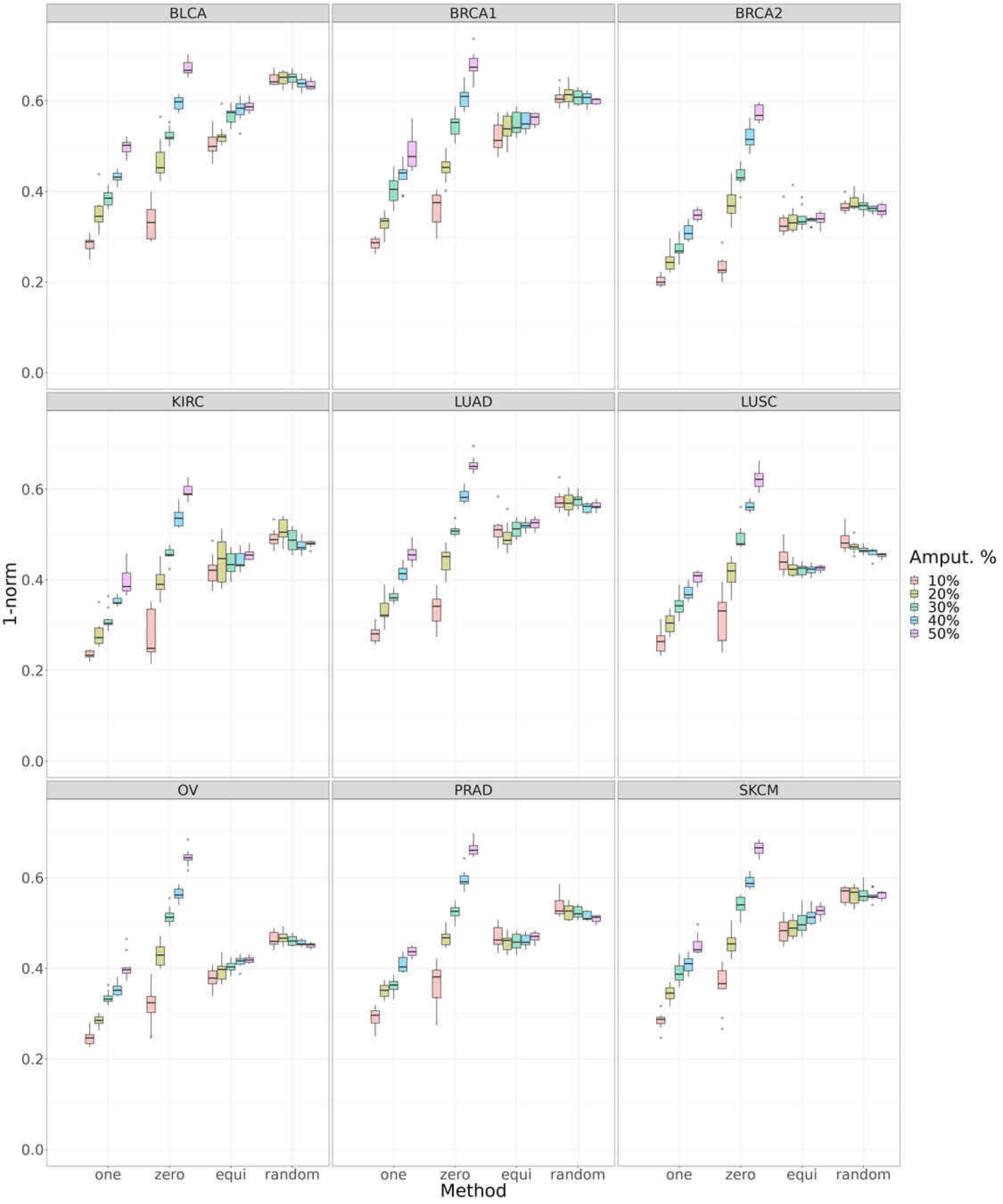


Fig. S5: Comparison of 1-norm recovery error among miss-SNF initialization approaches for each cancer. Each panel represents a different cancer dataset where the distribution of the 1-norm among amputated and complete datasets for different miss-SNF initializations (x-axis) and amputation percentages (represented by different colors) is depicted by means of boxplots.

2. *Enrichment of clinical variables and differential survival in the clusters* are considered to assess if clusterings are different in a biologically relevant way. We exploited the logrank test to evaluate if the identified clusters are statistically different in terms of survival. The enrichment of clinical variables in the clusters is performed considering a carefully selected set of meaningful clinical variables for each cancer (the complete list is available in Supplementary Section S3.2), where we considered the $\chi^2$ test for categorical variables and the Kruskal-Wallis test for numerical ones. As recommended in the work of Rappoport et al [Rappoport and Shamir, 2018], we exploited permutation-based tests in our analysis because all the above-mentioned hypothesis tests assume a $\chi^2$ distribution

of the statistic, which is hardly respected due to small sample size and imbalanced cluster sizes typical of TCGA datasets[6]. P-values obtained for the clinical variables are adjusted for multiple comparisons using Bonferroni correction. For both logrank test and enrichment of clinical variables the significance threshold is 0.05.

After the evaluation considering all numbers of clusters $nc = [2 .. 10]$, we also compare clustering results after the selection of the optimal $nc$. We consider two different approaches applied on the complete dataset:

1. A *modified eigengap method* that is proposed by NEMO [Rappoport and Shamir, 2019] as a modification of the *eigengap method* [Von Luxburg, 2007] used by SNF [Wang et al., 2014]. The *modified eigengap method* forces the selection of a higher number of clusters with respect to the classic eigengap approach, which the authors consider a desirable behaviour that improves the prognostic quality of cancer data.
2. The selection of the optimal number of clusters based on the enrichment of clinical variables and logrank tests. The best number of clusters is defined as the one leading to the highest number of enriched clinical variables and, in case of ties, to the lowest logrank test p-value on the complete dataset [7]. The idea behind this proposed approach is that a clustering enriched in many clinical variables should have been captured a biologically relevant grouping of the samples, thus being a good value for the choice of the optimal number of clusters.

Of note, MOFA+ [Argelaguet et al., 2020] is not presented as a clustering approach but it is a data integration method that outputs matrices that can be used for a wide range of downstream analysis, including clustering and classification tasks [Argelaguet et al., 2018]. Thus, any clustering approach and method to select the number of clusters could in principle be exploited.

Since the modified eigengap method works using a similarity matrix as input, we exploited the similarity matrix computed using the scaled exponential Euclidean kernel [Wang et al., 2014] for MOFA+.

Paired Wilcoxon signed rank test is used to compute win-tie-loss tables (see Section S5) to compare the results of different combinations of integration methods and clustering approaches considering (I) the different evaluated quality metrics for the reconstruction of pairwise similarities and (II) the agreement between clusterings obtained from complete and amputated datasets.

The comparison among miss-SNF, NEMO, and MOFA+ in terms of (I) pairwise similarity reconstruction and related preservation of graph topology in amputated datasets and (II) clustering results, is reported in Section 3.3.

### S7.2. Clustering algorithms

The following state-of-the-art clustering algorithms are used in our experiments to perform patients clustering:

- *k-means* Hastie et al. [2009] is a popular iterative clustering algorithm that minimizes the within-cluster sum of squares, thus that the average distance between points assigned to a cluster and the cluster mean is minimized. In particular, after an initial random selection of clusters centers, two phases are iteratively repeated until convergence: (I) assignment of points to the closest center, (II) recomputation of the centers. Convergence is reached when assignments do not change or, as implemented in the Hartigan and Wong algorithm Hartigan and Wong [1979] (default implementation in the used R package *stats* R Core Team [2023]), no switch of objects from one cluster to another is able to further reduce the objective function.
- *Spectral clustering (SP)* Von Luxburg [2007] processes a similarity network, $\mathbf{W} \in \Re^{n \times n}$ (being $n$ the number of points), and partitions it into $k$ clusters, so that points falling in the same clusters are highly similar, while points in different clusters are maximally dissimilar. The clustering problem is seen as a graph-partition problem and normalized spectral clustering (see below) solves a relaxed version of the Normalized cut problem Von Luxburg [2007]. Spectral clustering has the advantage of dealing with non-convex clusters, i.e. complex topological cluster structures, while k-means can not Hastie et al. [2009]. Among the most popular spectral clustering algorithms the one proposed by Ng et al. Ng et al. [2001] computes the normalized graph Laplacian as $\mathbf{L} = \mathbf{I} - \mathbf{D}^{-1/2}\mathbf{W}\mathbf{D}^{-1/2}$, where $\mathbf{I}$ is the identity matrix, $\mathbf{W}$ the affinity matrix, and $\mathbf{D}$ is the degree matrix having the degree of each point on the diagonal. Then, if $k$ is the number of clusters to be found, the first $k$ eigenvectors having the smallest eigenvalues are computed obtaining a lower-dimensional matrix $\mathbf{U} \in \mathbb{R}^{n \times k}$, which is normalized and then used as the new k-dimensional representation of the $n$ points. Finally k-means algorithm is used to cluster points. However, another variant of the algorithm is exploited by SNF and NEMO, and available in the package *SNFtool*. In this implementation, instead of leveraging the k-means algorithm for clustering, each sample (on rows of $\mathbf{U}$) is assigned to the cluster whose corresponding column has the maximum value. The algorithm from Ng et al. Ng et al. [2001] is implemented in the R package *sClust* Poisson-Caillault and Vincent [2021].

[6] We used the implementation of permutation tests from Rappoport et al [Rappoport and Shamir, 2018] work, available at `https://github.com/Shamir-Lab/Multi-Omics-Cancer-Benchmark/`. For the logrank test, the number of performed permutations is set to $min(max(\frac{10}{original\ p-value}, 10^3), 10^6)$ and $10^5$ additional permutations are performed until the stopping criterion is met, which consists in having the 95% confidence interval not crossing the significance threshold of 0.05 and a maximum number of iterations of $5 \times 10^6$. For the enrichment of clinical variables, batches of $10^3$ permutations are performed until the 95% confidence interval is not crossing the significance threshold of 0.05, with $10^5$ maximum number of iterations.

[7] In case of ties for both the number of enriched clinical variables and logrank test p-values, we select the number of clusters nearest to 6, which is a rough estimate of the number of clusters in cancer datasets [Rappoport and Shamir, 2019]. This situation happened in just 1 experiment out of 135.

- *Partitioning Around Medoids (PAM)* Kaufman and Rousseeuw [1990] is an alternative to k-means that is more robust to outliers and noise because, instead of using the mean of each cluster as a centroid, it exploits the concept of "medoid" (i.e. the most central point in the cluster Jin and Han [2010] defined by the point having minimum sum of dissimilarities with the other objects in the cluster).

  The final goal is to obtain a set of clusters where the average distances of objects belonging to the cluster and the cluster representative is minimized (equivalently the sum of the distances can be minimized). Briefly, the algorithm can be decomposed in an initial *build* phase and in the *swap* phase. In the build phase a starting set of $k$ representative points is selected, where the first point is the most central to all the dataset and the subsequent points are iteratively selected as medoids if they have a high number of close unselected objects while being far to already chosen representatives. Points that are not selected are assigned to the closest medoid. In the swap phase, the goal is to improve the set of medoids, and consequently the quality of the clusters, by swapping already selected points with objects that are not medoids. The swap that is most effective in terms of minimization of the cost function (i.e. sum of the dissimilarities between objects of a cluster and the corresponding medoid) is maintained and swaps are performed till the cost function stops to decrease.

### S8. Classification experiments - supervised feature selection and hyper-parameter tuning via internal-holdout validation

Random Forest (RF) generalization performance are evaluated using 10 stratified external holdouts (90 : 10, training:test set), while 10 stratified internal holdouts (90 : 10, training:test set) are considered to perform feature selection and hyper-parameters tuning. Each internal holdout is exploited to train a RF and the average feature importance (i.e. Mean Decrease in Accuracy) across all internal holdouts is used for feature selection. The most discriminative variables are the ones with a cumulative sum lower than 0.95 (average feature importance values are sorted in decreasing order and normalized to unitary sum) and we force the selection of at least 3 variables. Hyper-parameter tuning is performed using only the subset of discriminative features $f_{sel}$ where the tuned hyper-parameters are the number of trees *ntree* and the number of variables *mtry* randomly picked to be considered at each split. In particular, the hyper-parameter grid to evaluate is defined as follows.

If $f_{sel} > 10$, then:

- $ntree = \{500, 750, 1000\}$
- $mtry = \left\{ \left\lfloor \frac{f_{sel}}{10} \right\rfloor, \left\lfloor \frac{f_{sel}}{5} \right\rfloor, \left\lfloor \sqrt{f_{sel}} \right\rfloor \right\}$

else:

- $ntree = \{100, 500\}$
- $mtry = \left\lfloor \sqrt{f_{sel}} \right\rfloor$

Note that only unique values of *mtry* in the range $0 < mtry < f_{sel}$ are considered. The combination of hyper-parameters that reaches the highest average AUC across the internal holdouts is selected.

### S9. Supplementary Tables

Table S5: Descriptive statistics for cancer datasets. Column $N$ reports the number of cases; column $\frac{N_{pos}^{OS}}{N}$ reports the balance ratio for OS label, measured as the ratio between the number of OS positive cases (OS = 1) and all the cases in the dataset; column $\frac{N_{pos}^{PFI}}{N}$ reports the balance ratio for PFI label, measured as the ratio between the number of PFI positive cases (PFI = 1) and all the cases in the dataset. Column *View* indicates the different data sources and for each one of them the following information are provided: $D$ (raw) reports the original dimension of each view and $D$ (RPCA) shows the dimensionality after dimensionality reduction using RPCA guided by id-estimation. (continue...)

| **Dataset** | **N** | $\frac{N_{pos}^{OS}}{N}$ | $\frac{N_{pos}^{PFI}}{N}$ | **View** | **D (raw)** | **D (RPCA)** |
|---|---|---|---|---|---|---|
| **BLCA** | 335 | 0.45 | 0.43 | miRNA | 469 | 47 |
| | | | | mRNA | 12276 | 42 |
| | | | | proteins | 183 | 24 |
| | | | | methylation | 315551 | 44 |
| **BRCA1** | 317 | 0.13 | 0.11 | miRNA | 496 | 49 |
| | | | | mRNA | 12242 | 40 |
| | | | | proteins | 202 | 24 |
| | | | | methylation | 289962 | 39 |
| **BRCA2** | 128 | 0.11 | 0.13 | miRNA | 502 | 34 |
| | | | | mRNA | 8128 | 37 |
| | | | | proteins | 192 | 18 |
| | | | | methylation | 278099 | 39 |
| **KIRC** | 169 | 0.28 | 0.34 | miRNA | 364 | 51 |
| | | | | mRNA | 7942 | 37 |
| | | | | proteins | 186 | 21 |
| | | | | methylation | 319740 | 50 |
| **LUAD** | 300 | 0.40 | 0.41 | miRNA | 465 | 47 |
| | | | | mRNA | 11131 | 44 |
| | | | | proteins | 179 | 24 |
| | | | | methylation | 331828 | 43 |
| **LUSC** | 228 | 0.41 | 0.37 | miRNA | 491 | 47 |
| | | | | mRNA | 11473 | 40 |
| | | | | proteins | 178 | 25 |
| | | | | methylation | 273884 | 40 |
| **OV** | 226 | 0.63 | 0.71 | miRNA | 308 | 24 |
| | | | | mRNA | 11731 | 47 |
| | | | | proteins | 186 | 27 |
| | | | | methylation | 13296 | 51 |
| **PRAD** | 337 | 0.02 | 0.18 | miRNA | 457 | 56 |
| | | | | mRNA | 8887 | 41 |
| | | | | proteins | 169 | 20 |
| | | | | methylation | 301920 | 47 |
| **SKCM** | 334 | 0.45 | 0.63 | miRNA | 523 | 45 |
| | | | | mRNA | 13050 | 39 |
| | | | | proteins | 186 | 24 |
| | | | | methylation | 311405 | 48 |

**Table S6.** Win-tie-loss table comparing different data integration approaches in terms of 1-norm. Paired Wilcoxon test is exploited to compare different methods, where results are paired considering the different amputation percentages, their random "repetitions" (i.e. 10 randomly amputated datasets for each amputation percentage) and by grouping the nine types of considered cancers.

| | 1-norm | | |
|---|---|---|---|
| Methods | wins | ties | losses |
| miss-SNF one | 3 | 0 | 0 |
| miss-SNF equidistant | 2 | 0 | 1 |
| NEMO | 1 | 0 | 2 |
| MOFA+ | 0 | 0 | 3 |

**Table S7.** Win-tie-loss table comparing different data integration approaches in terms of graph similarity. Paired Wilcoxon test is exploited to compare different methods, where results are paired considering the different amputation percentages, their random "repetitions" (i.e. 10 randomly amputated datasets for each amputation percentage) and considered cancers.

| | Similarity (3-RWK) | | |
|---|---|---|---|
| Methods | wins | ties | losses |
| miss-SNF one | 3 | 0 | 0 |
| miss-SNF equidistant | 2 | 0 | 1 |
| MOFA+ | 1 | 0 | 2 |
| NEMO | 0 | 0 | 3 |

**Table S8.** Win-tie-loss table comparing data integration approaches considering different clustering algorithms using NMI. Paired Wilcoxon test is exploited to compare different experiments, where results are paired considering the different amputation percentages and their random "repetitions" (i.e. 10 randomly amputated datasets for each amputation percentage), the evaluated number of clusters and cancer.

| | NMI | | |
|---|---|---|---|
| Methods | wins | ties | losses |
| NEMO - sp (SNFtool) | 14 | 0 | 0 |
| NEMO - sp (sClust) | 13 | 0 | 1 |
| NEMO - k-means (similarities) | 12 | 0 | 2 |
| one - k-means (similarities) | 11 | 0 | 3 |
| one - sp (sClust) | 10 | 0 | 4 |
| one - pam (similarities) | 9 | 0 | 5 |
| equi - k-means (similarities) | 8 | 0 | 6 |
| equi - sp (sClust) | 7 | 0 | 7 |
| NEMO - pam (similarities) | 6 | 0 | 8 |
| equi - pam (similarities) | 5 | 0 | 9 |
| MOFA+ - k-means (factors) | 4 | 0 | 10 |
| MOFA+ - sp (sClust) | 3 | 0 | 11 |
| MOFA+ - pam (factors) | 2 | 0 | 12 |
| MOFA+ - k-means (similarities) | 0 | 1 | 13 |
| MOFA+ - pam (similarities) | 0 | 1 | 13 |

**Table S9.** Win-tie-loss table comparing data integration approaches considering different clustering algorithms using NVI. Paired Wilcoxon test is exploited to compare different experiments, where results are paired considering the different amputation percentages and their random "repetitions" (i.e. 10 randomly amputated datasets for each amputation percentage), the evaluated number of clusters and cancer.

| | NVI | | |
|---|---|---|---|
| Methods | wins | ties | losses |
| NEMO - sp (SNFtool) | 14 | 0 | 0 |
| NEMO - sp (sClust) | 13 | 0 | 1 |
| NEMO - k-means (similarities) | 12 | 0 | 2 |
| one - k-means (similarities) | 11 | 0 | 3 |
| one - sp (sClust) | 10 | 0 | 4 |
| one - pam (similarities) | 9 | 0 | 5 |
| equi - k-means (similarities) | 8 | 0 | 6 |
| equi - sp (sClust) | 7 | 0 | 7 |
| NEMO - pam (similarities) | 6 | 0 | 8 |
| equi - pam (similarities) | 5 | 0 | 9 |
| MOFA+ - k-means (factors) | 4 | 0 | 10 |
| MOFA+ - sp (sClust) | 3 | 0 | 11 |
| MOFA+ - pam (factors) | 2 | 0 | 12 |
| MOFA+ - k-means (similarities) | 0 | 1 | 13 |
| MOFA+ - pam (similarities) | 0 | 1 | 13 |

**Table S10.** Win-tie-loss table comparing data integration approaches considering different clustering algorithms using Adjusted Rand Index. Paired Wilcoxon test is exploited to compare different experiments, where results are paired considering the different amputation percentages and their random "repetitions" (i.e. 10 randomly amputated datasets for each amputation percentage), the evaluated number of clusters and cancer.

| | Adjusted Rand Index | | |
|---|---|---|---|
| Methods | wins | ties | losses |
| NEMO - sp (SNFtool) | 14 | 0 | 0 |
| NEMO - sp (sClust) | 13 | 0 | 1 |
| NEMO - k-means (similarities) | 12 | 0 | 2 |
| one - k-means (similarities) | 11 | 0 | 3 |
| one - sp (sClust) | 10 | 0 | 4 |
| one - pam (similarities) | 9 | 0 | 5 |
| equi - k-means (similarities) | 8 | 0 | 6 |
| NEMO - pam (similarities) | 7 | 0 | 7 |
| equi - pam (similarities) | 6 | 0 | 8 |
| equi - sp (sClust) | 5 | 0 | 9 |
| MOFA+ - k-means (factors) | 4 | 0 | 10 |
| MOFA+ - k-means (similarities) | 3 | 0 | 11 |
| MOFA+ - pam (factors) | 0 | 2 | 12 |
| MOFA+ - pam (similarities) | 0 | 2 | 12 |
| MOFA+ - sp (sClust) | 0 | 2 | 12 |

**Table S11.** Win-tie-loss table comparing the number of enriched variables for the partitions obtained by the partial data fusion+clustering approaches. Wilcoxon test is exploited to compare the evaluated methods, pairing the different amputation percentages and their random "repetitions" (i.e. 10 randomly amputated datasets for each amputation percentage), the evaluated number of clusters and cancer.

| | Number of enriched clinical variables | | |
|---|---|---|---|
| Methods | wins | ties | losses |
| one - k-means (similarities) | 14 | 0 | 0 |
| one - sp (sClust) | 13 | 0 | 1 |
| one - pam (similarities) | 12 | 0 | 2 |
| NEMO - k-means (similarities) | 11 | 0 | 3 |
| NEMO - sp (SNFtool) | 10 | 0 | 4 |
| equi - k-means (similarities) | 9 | 0 | 5 |
| NEMO - sp (sClust) | 7 | 1 | 6 |
| NEMO - pam (similarities) | 6 | 2 | 6 |
| equi - pam (similarities) | 6 | 1 | 7 |
| equi - sp (sClust) | 5 | 0 | 9 |
| MOFA+ - k-means (factors) | 3 | 1 | 10 |
| MOFA+ - pam (factors) | 3 | 1 | 10 |
| MOFA+ - pam (similarities) | 1 | 1 | 12 |
| MOFA+ - sp (sClust) | 1 | 1 | 12 |
| MOFA+ - k-means (similarities) | 0 | 0 | 14 |

**Table S12.** Win-tie-loss table comparing the frequency of clusterings showing differential survival. Wilcoxon test is exploited to compare the evaluated methods, pairing the different amputation percentages, the evaluated number of clusters and cancer.

| | Frequency of significant logrank tests | | |
|---|---|---|---|
| Methods | wins | ties | losses |
| one - k-means (similarities) | 14 | 0 | 0 |
| one - sp (sClust) | 11 | 2 | 1 |
| one - pam (similarities) | 10 | 3 | 1 |
| NEMO - sp (SNFtool) | 10 | 3 | 1 |
| NEMO - k-means (similarities) | 10 | 2 | 2 |
| equi - k-means (similarities) | 9 | 0 | 5 |
| equi - pam (similarities) | 6 | 2 | 6 |
| equi - sp (sClust) | 6 | 2 | 6 |
| NEMO - sp (sClust) | 6 | 2 | 6 |
| NEMO - pam (similarities) | 5 | 0 | 9 |
| MOFA+ - k-means (factors) | 2 | 2 | 10 |
| MOFA+ - pam (factors) | 1 | 3 | 10 |
| MOFA+ - pam (similarities) | 1 | 3 | 10 |
| MOFA+ - sp (sClust) | 1 | 2 | 11 |
| MOFA+ - k-means (similarities) | 0 | 0 | 14 |

**Table S13.** Win-tie-loss table comparing data integration approaches considering different clustering algorithms using NMI after the selection of the optimal number of clusters. The left part of the table shows the results using the clin-surv method to select the number of clusters while on the right the results exploiting the eigengap approach are reported. Paired Wilcoxon test is leveraged to compare different experiments, where results are paired considering the amputation percentages, their random "repetitions" (i.e. 10 randomly amputated datasets for each amputation percentage) and cancers.

| | NMI clin-surv method | | | | NMI eigengap method | | |
|---|---|---|---|---|---|---|---|
| Methods | wins | ties | losses | Methods | wins | ties | losses |
| NEMO - sp (SNFtool) | 14 | 0 | 0 | NEMO - sp (SNFtool) | 14 | 0 | 0 |
| NEMO - sp (sClust) | 13 | 0 | 1 | NEMO - sp (sClust) | 13 | 0 | 1 |
| NEMO - k-means (similarities) | 12 | 0 | 2 | NEMO - k-means (similarities) | 12 | 0 | 2 |
| one - k-means (similarities) | 11 | 0 | 3 | one - k-means (similarities) | 11 | 0 | 3 |
| one - sp (sClust) | 10 | 0 | 4 | one - pam (similarities) | 9 | 1 | 4 |
| one - pam (similarities) | 9 | 0 | 5 | one - sp (sClust) | 9 | 1 | 4 |
| equi - k-means (similarities) | 8 | 0 | 6 | equi - k-means (similarities) | 8 | 0 | 6 |
| NEMO - pam (similarities) | 6 | 1 | 7 | equi - sp (sClust) | 7 | 0 | 7 |
| equi - sp (sClust) | 5 | 2 | 7 | MOFA+ - k-means (factors) | 5 | 1 | 8 |
| equi - pam (similarities) | 5 | 1 | 8 | NEMO - pam (similarities) | 4 | 2 | 8 |
| MOFA+ - k-means (factors) | 4 | 0 | 10 | MOFA+ - sp (sClust) | 3 | 2 | 9 |
| MOFA+ - sp (sClust) | 3 | 0 | 11 | MOFA+ - pam (factors) | 2 | 1 | 11 |
| MOFA+ - pam (factors) | 2 | 0 | 12 | equi - pam (similarities) | 1 | 3 | 10 |
| MOFA+ - pam (similarities) | 1 | 0 | 13 | MOFA+ - pam (similarities) | 0 | 2 | 12 |
| MOFA+ - k-means (similarities) | 0 | 0 | 14 | MOFA+ - k-means (similarities) | 0 | 1 | 13 |

**Table S14.** Win-tie-loss table comparing data integration approaches considering different clustering algorithms using NVI after the selection of the optimal number of clusters. The left part of the table shows the results using the clin-surv method to select the number of clusters while on the right the results exploiting the eigengap approach are reported. Paired Wilcoxon test is leveraged to compare different experiments, where results are paired considering the amputation percentages, their random "repetitions" (i.e. 10 randomly amputated datasets for each amputation percentage) and cancers.

| | NVI clin-surv method | | | | NVI eigengap method | | |
|---|---|---|---|---|---|---|---|
| Methods | wins | ties | losses | Methods | wins | ties | losses |
| NEMO - sp (SNFtool) | 14 | 0 | 0 | NEMO - sp (SNFtool) | 14 | 0 | 0 |
| NEMO - sp (sClust) | 13 | 0 | 1 | NEMO - sp (sClust) | 13 | 0 | 1 |
| NEMO - k-means (similarities) | 12 | 0 | 2 | NEMO - k-means (similarities) | 12 | 0 | 2 |
| one - k-means (similarities) | 11 | 0 | 3 | one - k-means (similarities) | 11 | 0 | 3 |
| one - sp (sClust) | 10 | 0 | 4 | one - pam (similarities) | 9 | 1 | 4 |
| one - pam (similarities) | 9 | 0 | 5 | one - sp (sClust) | 9 | 1 | 4 |
| equi - k-means (similarities) | 8 | 0 | 6 | equi - k-means (similarities) | 8 | 0 | 6 |
| NEMO - pam (similarities) | 6 | 1 | 7 | equi - sp (sClust) | 7 | 0 | 7 |
| equi - sp (sClust) | 5 | 2 | 7 | MOFA+ - k-means (factors) | 5 | 1 | 8 |
| equi - pam (similarities) | 5 | 1 | 8 | NEMO - pam (similarities) | 4 | 2 | 8 |
| MOFA+ - k-means (factors) | 4 | 0 | 10 | MOFA+ - sp (sClust) | 3 | 2 | 9 |
| MOFA+ - sp (sClust) | 3 | 0 | 11 | MOFA+ - pam (factors) | 2 | 1 | 11 |
| MOFA+ - pam (factors) | 2 | 0 | 12 | equi - pam (similarities) | 1 | 3 | 10 |
| MOFA+ - pam (similarities) | 1 | 0 | 13 | MOFA+ - pam (similarities) | 0 | 2 | 12 |
| MOFA+ - k-means (similarities) | 0 | 0 | 14 | MOFA+ - k-means (similarities) | 0 | 1 | 13 |

**Table S15.** Win-tie-loss table comparing data integration approaches considering different clustering algorithms using Adjusted Rand Index (ARI) after the selection of the optimal number of clusters. The left part of the table shows the results using the clin-surv method to select the number of clusters while on the right the results exploiting the eigengap approach are reported. Paired Wilcoxon test is leveraged to compare different experiments, where results are paired considering the amputation percentages, their random "repetitions" (i.e. 10 randomly amputated datasets for each amputation percentage) and cancers.

| | ARI clin-surv method | | | | ARI eigengap method | | |
|---|---|---|---|---|---|---|---|
| Methods | wins | ties | losses | Methods | wins | ties | losses |
| NEMO - sp (SNFtool) | 14 | 0 | 0 | NEMO - sp (SNFtool) | 14 | 0 | 0 |
| NEMO - sp (sClust) | 13 | 0 | 1 | NEMO - sp (sClust) | 13 | 0 | 1 |
| NEMO - k-means (similarities) | 12 | 0 | 2 | NEMO - k-means (similarities) | 12 | 0 | 2 |
| one - k-means (similarities) | 11 | 0 | 3 | one - k-means (similarities) | 11 | 0 | 3 |
| one - sp (sClust) | 10 | 0 | 4 | one - pam (similarities) | 10 | 0 | 4 |
| one - pam (similarities) | 9 | 0 | 5 | one - sp (sClust) | 9 | 0 | 5 |
| NEMO - pam (similarities) | 8 | 0 | 6 | equi - k-means (similarities) | 8 | 0 | 6 |
| equi - k-means (similarities) | 7 | 0 | 7 | NEMO - pam (similarities) | 7 | 0 | 7 |
| equi - pam (similarities) | 6 | 0 | 8 | equi - pam (similarities) | 5 | 1 | 8 |
| MOFA+ - k-means (factors) | 3 | 2 | 9 | equi - sp (sClust) | 5 | 1 | 8 |
| MOFA+ - k-means (similarities) | 3 | 2 | 9 | MOFA+ - k-means (similarities) | 4 | 0 | 10 |
| equi - sp (sClust) | 3 | 2 | 9 | MOFA+ - k-means (factors) | 3 | 0 | 11 |
| MOFA+ - sp (sClust) | 2 | 0 | 12 | MOFA+ - pam (factors) | 1 | 1 | 12 |
| MOFA+ - pam (similarities) | 1 | 0 | 13 | MOFA+ - pam (similarities) | 1 | 1 | 12 |
| MOFA+ - pam (factors) | 0 | 0 | 14 | MOFA+ - sp (sClust) | 0 | 0 | 14 |

**Table S16.** Win-tie-loss table comparing partial data fusion + clustering algorithms in terms of the the number of enriched clinical variables after the selection of the optimal number of clusters is performed on the complete dataset. Wilcoxon test is exploited to compare the evaluated methods, pairing the different amputation percentages, their random "repetitions" (i.e. 10 randomly amputated datasets for each amputation percentage) and cancers. Results using clin-surv (eigengap) method for optimal number of clusters selection are on the left (right) subtable.

| Number of enriched clinical variables | | | | | | | |
|---|---|---|---|---|---|---|---|
| | clin-surv method | | | | eigengap method | | |
| Methods | wins | ties | losses | Methods | wins | ties | losses |
| one - k-means (similarities) | 14 | 0 | 0 | NEMO - k-means (similarities) | 14 | 0 | 0 |
| NEMO - k-means (similarities) | 12 | 1 | 1 | one - k-means (similarities) | 12 | 1 | 1 |
| NEMO - sp (SNFtool) | 12 | 1 | 1 | one - sp (sClust) | 10 | 3 | 1 |
| one - pam (similarities) | 9 | 2 | 3 | NEMO - sp (sClust) | 10 | 2 | 2 |
| NEMO - sp (sClust) | 9 | 2 | 3 | one - pam (similarities) | 9 | 3 | 2 |
| one - sp (sClust) | 9 | 2 | 3 | NEMO - sp (SNFtool) | 9 | 1 | 4 |
| equi - k-means (similarities) | 8 | 0 | 6 | equi - k-means (similarities) | 7 | 1 | 6 |
| NEMO - pam (similarities) | 7 | 0 | 7 | equi - pam (similarities) | 7 | 1 | 6 |
| equi - pam (similarities) | 6 | 0 | 8 | NEMO - pam (similarities) | 5 | 1 | 8 |
| equi - sp (sClust) | 5 | 0 | 9 | equi - sp (sClust) | 4 | 2 | 8 |
| MOFA+ - pam (factors) | 3 | 1 | 10 | MOFA+ - pam (factors) | 2 | 3 | 9 |
| MOFA+ - pam (similarities) | 2 | 2 | 10 | MOFA+ - k-means (factors) | 1 | 3 | 10 |
| MOFA+ - k-means (factors) | 2 | 1 | 11 | MOFA+ - pam (similarities) | 1 | 3 | 10 |
| MOFA+ - sp (sClust) | 1 | 0 | 13 | MOFA+ - sp (sClust) | 1 | 2 | 11 |
| MOFA+ - k-means (similarities) | 0 | 0 | 14 | MOFA+ - k-means (similarities) | 0 | 0 | 14 |

**Table S17.** Win-tie-loss table comparing data integration approaches considering different clustering algorithms using the frequency of significant logrank tests after selecting the optimal number of clusters. Wilcoxon test is exploited to compare the evaluated methods, pairing the different amputation percentages and cancers. The left part of the table shows the results using the clin-surv while on the right the results exploiting the eigengap approach are reported.

| Frequency of significant logrank tests | | | | | | | |
|---|---|---|---|---|---|---|---|
| | clin-surv method | | | | eigengap method | | |
| Methods | wins | ties | losses | Methods | wins | ties | losses |
| one - k-means (similarities) | 11 | 3 | 0 | NEMO - k-means (similarities) | 11 | 3 | 0 |
| one - pam (similarities) | 10 | 4 | 0 | one - k-means (similarities) | 10 | 4 | 0 |
| NEMO - k-means (similarities) | 9 | 5 | 0 | NEMO - sp (sClust) | 10 | 4 | 0 |
| NEMO - sp (SNFtool) | 9 | 5 | 0 | NEMO - sp (SNFtool) | 10 | 4 | 0 |
| one - sp (sClust) | 8 | 4 | 2 | one - pam (similarities) | 7 | 6 | 1 |
| NEMO - sp (sClust) | 5 | 8 | 1 | equi - k-means (similarities) | 6 | 4 | 4 |
| NEMO - pam (similarities) | 3 | 7 | 4 | one - sp (sClust) | 6 | 4 | 4 |
| MOFA+ - k-means (factors) | 3 | 6 | 5 | equi - pam (similarities) | 5 | 5 | 4 |
| equi - k-means (similarities) | 2 | 7 | 5 | equi - sp (sClust) | 2 | 7 | 5 |
| equi - sp (sClust) | 2 | 7 | 5 | MOFA+ - pam (factors) | 1 | 6 | 7 |
| equi - pam (similarities) | 2 | 6 | 6 | MOFA+ - k-means (factors) | 1 | 5 | 8 |
| MOFA+ - pam (factors) | 1 | 7 | 6 | MOFA+ - pam (similarities) | 1 | 5 | 8 |
| MOFA+ - pam (similarities) | 1 | 5 | 8 | NEMO - pam (similarities) | 1 | 5 | 8 |
| MOFA+ - sp (sClust) | 1 | 2 | 11 | MOFA+ - sp (sClust) | 1 | 4 | 9 |
| MOFA+ - k-means (similarities) | 0 | 0 | 14 | MOFA+ - k-means (similarities) | 0 | 0 | 14 |
