## Supplementary Figures 1 for "miss-SNF: a multimodal patient similarity network integration approach to handle completely missing data sources"

[1]Dipartimento di Informatica "Giovanni Degli Antoni", Università degli Studi di Milano, Via Giovanni Celoria 18, 20133, Italy, [2]European Commission, Joint Research Centre (JRC), Ispra, Italy, [3]Department of Biomedical Sciences, Humanitas University, Via Rita Levi Montalcini 4, 20072, Pieve Emanuele (MI), Italy, [4]Department of Pathology, IRCCS Humanitas Clinical and Research Hospital, Via Alessandro Manzoni 56, 20089, Rozzano (MI), Italy, [5]Environmental Genomics and Systems Biology Division, Lawrence Berkeley National Laboratory, Berkeley, Ca, USA and [6]ELLIS - European Laboratory for Learning and Intelligent Systems, Milan, Italy

### S10. Supplementary Figures

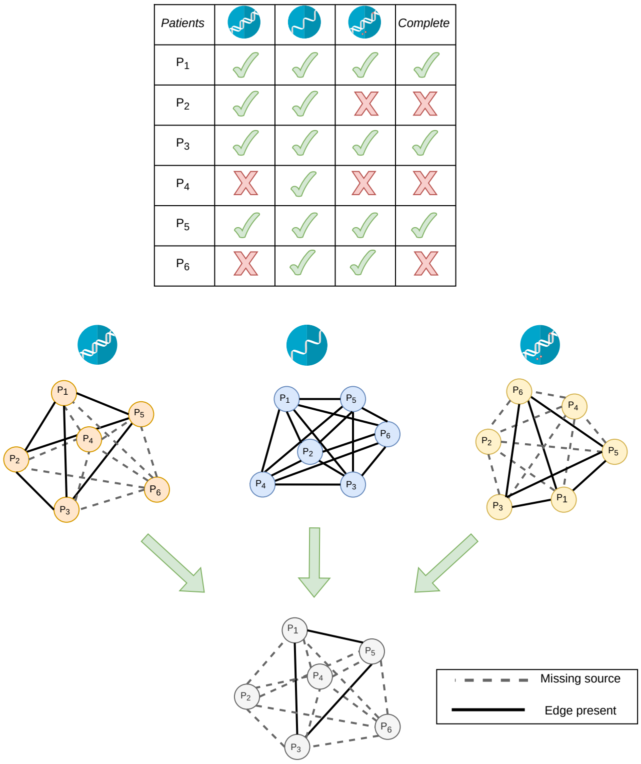


Fig. S6: Schematic example of partial dataset. Partial datasets are multimodal datasets characterized by the presence of samples having one or more data sources completely, or nearly completely, missing. Three different omics data sources are shown (i.e. genome, transcriptome and methylome) and missing modalities are represented by a $X$. The last column "Complete" indicates if the sample has all data sources, otherwise it is a "partial sample". The corresponding similarity matrices and the final integrated matrix are shown below, where missing data sources lead to incomplete unimodal networks that prevent the computation of similarities in the integrated network.

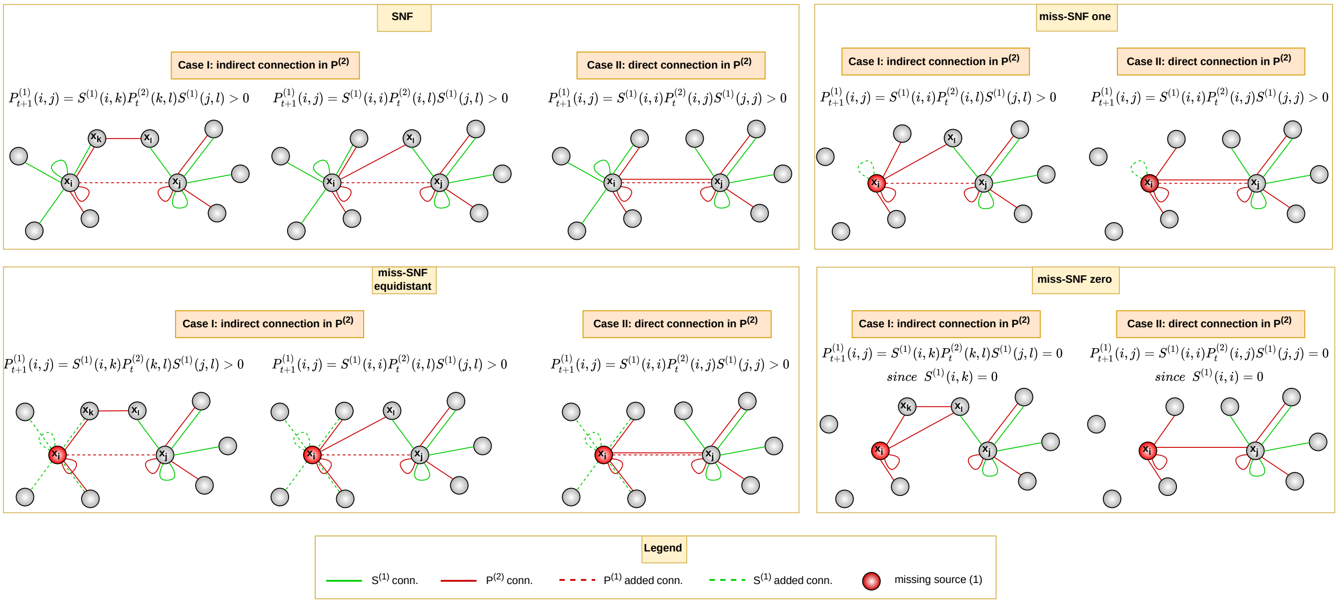


Fig. S7: SNF and miss-SNF diffusion process between two data sources focusing on nodes $x_i$ and $x_j$. We depict the diffusion process between nodes $x_i$ and $x_j$ considering two data sources represented by green (source 1) and red (source 2) edges in a toy graph, along with algorithm update formulas. The diffusion process updates the global similarity matrix $\mathbf{P}^{(1)}$ of data source 1 using information coming from the local similarity matrix $\mathbf{S}^{(1)}$ of the same data source (green edges) and the global similarity matrices $\mathbf{P}^{(2)}$ of the other data sources (red edges). In the SNF algorithm (upper left box), a novel edge (dotted line) can be added between nodes $x_i$ and $x_j$ if (I) their local neighbourhoods (e.g. $x_k$ and $x_l$) are connected and/or (II) they present a direct connection in the global similarity matrix which can be modified during the diffusion process. In miss-SNF *one* (upper right box), we set in the matrices $\mathbf{W}^{(1)}$, $\mathbf{P}^{(1)}$, $\mathbf{S}^{(1)}$ having missing patients, the similarity of the missing patient $x_i$ with himself to 1 and all the other similarities to zero, thus cutting the neighbourhood edges of $x_i$ with the exception of the self-loop. The passage of information is still allowed (I) if there is a common neighbour $x_l$ between $x_i$ in data source 2 and $x_j$ in the local similarity matrix of the missing data source 1 or (II) if $x_i$ and $x_j$ are directly connected in the source 2. In miss-SNF *equidistant* (lower left box), we set the similarity of the patient $x_i$ in the missing data source 1 to 0.5, while all the other similarities are set to the same small value. The patient is equally similar to all the other patients but more similar to himself and the diffusion process is the same depicted for SNF. In miss-SNF *zero* (lower right box), we cut all the edges of $x_i$ in data source 1 (i.e. $\mathbf{S}^{(1)}(i,k) = 0$) halting the passage of information between $x_i$ and $x_j$ because the result of the update formula is always 0.

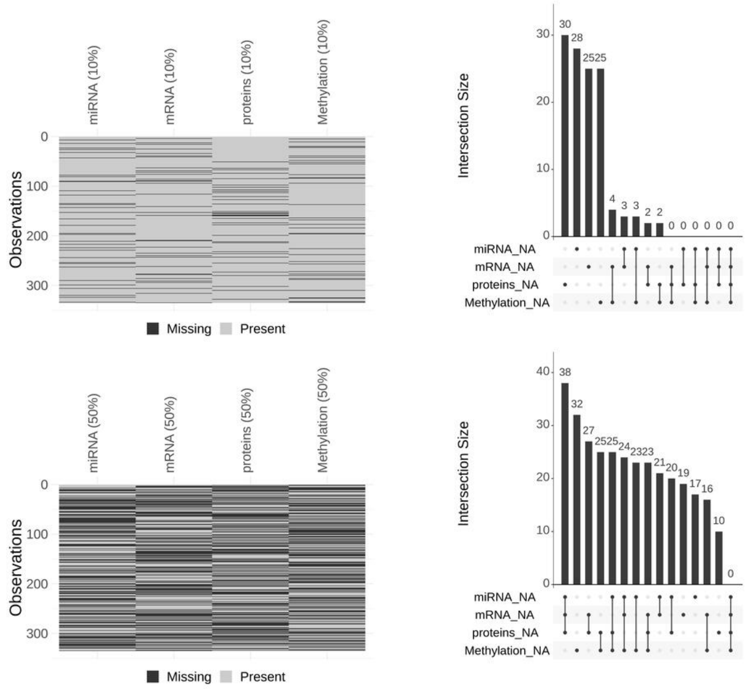


Fig. S8: Missing pattern for BLCA amputated dataset. Upper figure shows 10% of completely missing samples; Lower figure refers to 50% of completely missing samples. For both figures, the left panel shows for each sample in the dataset which data source is missing (black color) and which is present (grey color). Right panel shows a barplot with the number of samples that have a specific combination of data sources completely missing, where each combination is represented by the presence of black dots below the corresponding bar.

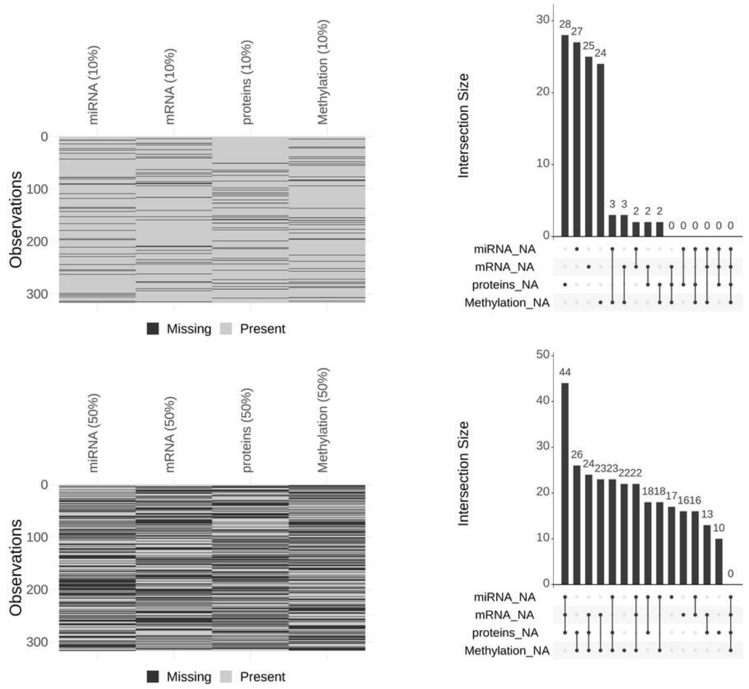


Fig. S9: Missing pattern for BRCA1 amputated dataset. Upper figure shows 10% of completely missing samples; Lower figure refers to 50% of completely missing samples. For both figures, the left panel shows for each sample in the dataset which data source is missing (black color) and which is present (grey color). Right panel shows a barplot with the number of samples that have a specific combination of data sources completely missing, where each combination is represented by the presence of black dots below the corresponding bar.

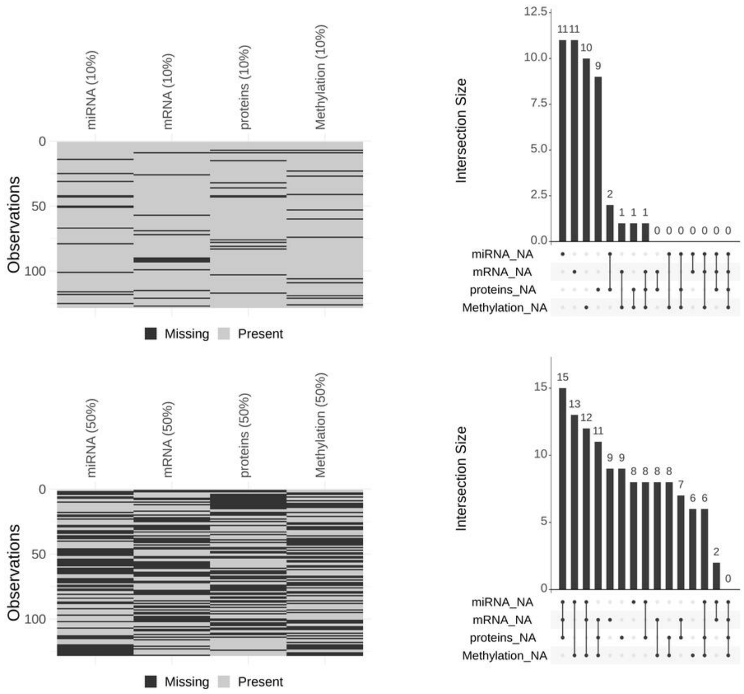


Fig. S10: Missing pattern for BRCA2 amputated dataset. Upper figure shows 10% of completely missing samples; Lower figure refers to 50% of completely missing samples. For both figures, the left panel shows for each sample in the dataset which data source is missing (black color) and which is present (grey color). Right panel shows a barplot with the number of samples that have a specific combination of data sources completely missing, where each combination is represented by the presence of black dots below the corresponding bar.

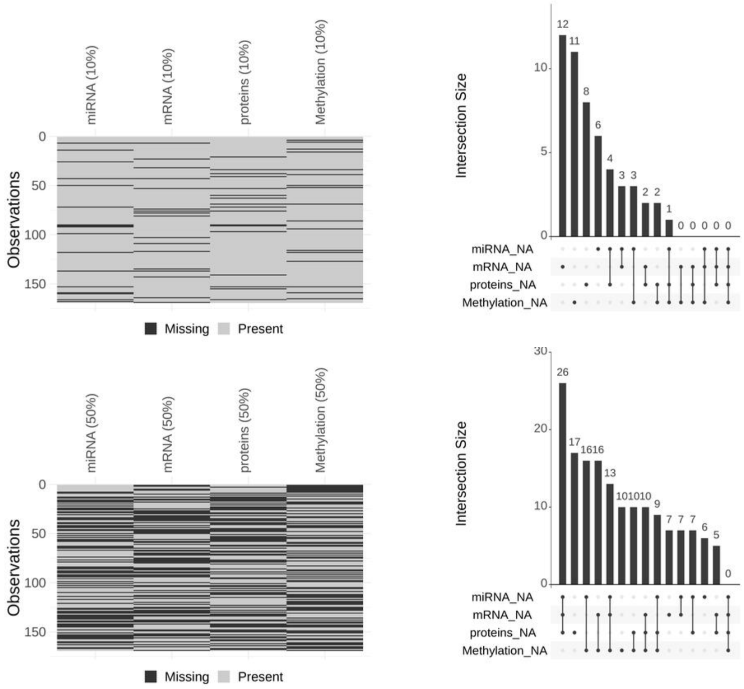


Fig. S11: Missing pattern for KIRC amputated dataset. Upper figure shows 10% of completely missing samples; Lower figure refers to 50% of completely missing samples. For both figures, the left panel shows for each sample in the dataset which data source is missing (black color) and which is present (grey color). Right panel shows a barplot with the number of samples that have a specific combination of data sources completely missing, where each combination is represented by the presence of black dots below the corresponding bar.

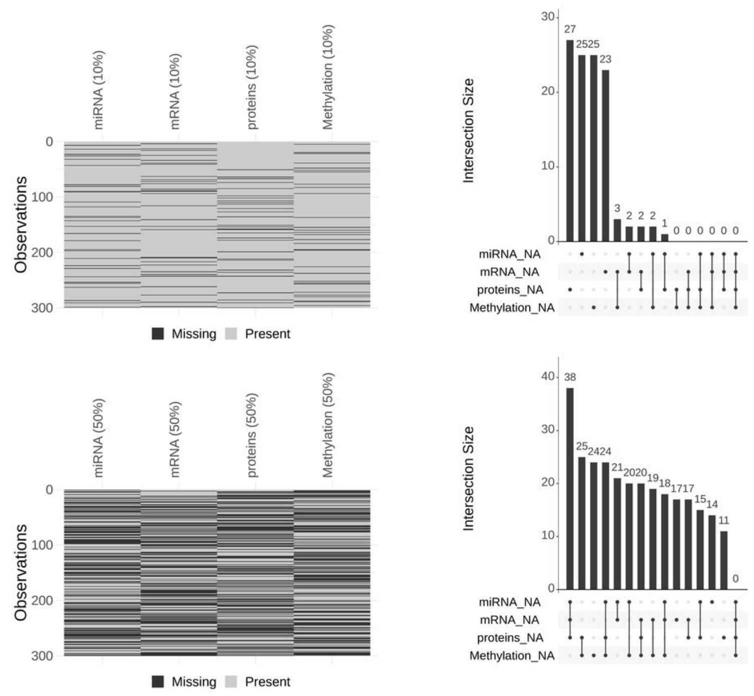


Fig. S12: Missing pattern for LUAD amputated dataset. Upper figure shows 10% of completely missing samples; Lower figure refers to 50% of completely missing samples. For both figures, the left panel shows for each sample in the dataset which data source is missing (black color) and which is present (grey color). Right panel shows a barplot with the number of samples that have a specific combination of data sources completely missing, where each combination is represented by the presence of black dots below the corresponding bar.

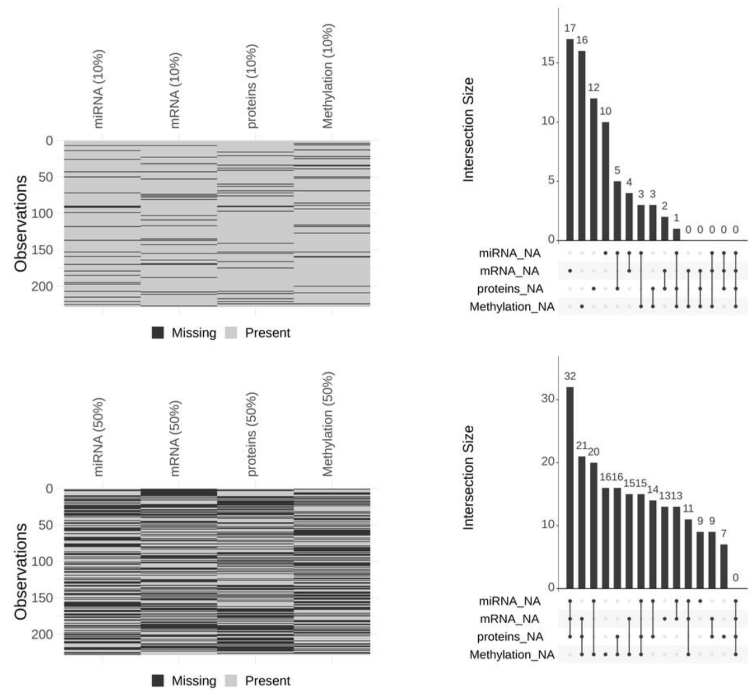


Fig. S13: Missing pattern for LUSC amputated dataset. Upper figure shows 10% of completely missing samples; Lower figure refers to 50% of completely missing samples. For both figures, the left panel shows for each sample in the dataset which data source is missing (black color) and which is present (grey color). Right panel shows a barplot with the number of samples that have a specific combination of data sources completely missing, where each combination is represented by the presence of black dots below the corresponding bar.

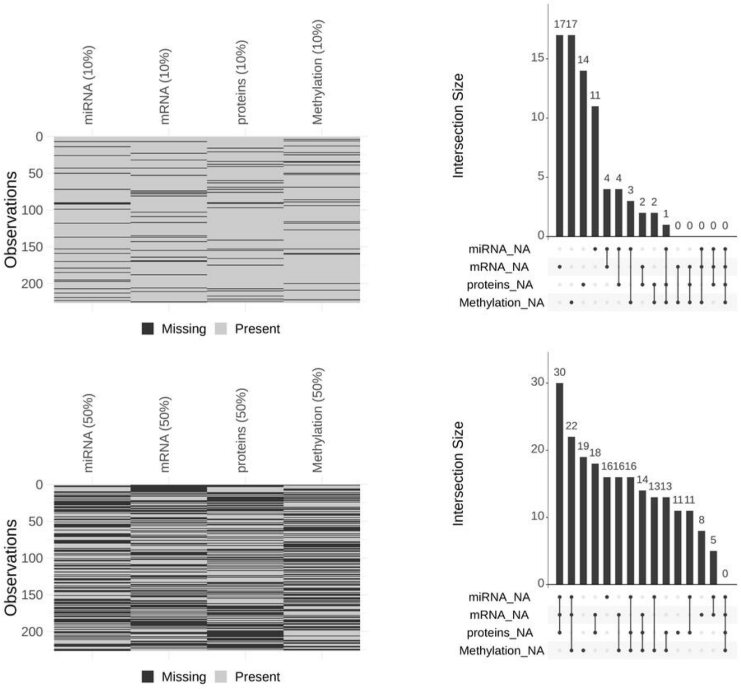


Fig. S14: Missing pattern for OV amputated dataset. Upper figure shows 10% of completely missing samples; Lower figure refers to 50% of completely missing samples. For both figures, the left panel shows for each sample in the dataset which data source is missing (black color) and which is present (grey color). Right panel shows a barplot with the number of samples that have a specific combination of data sources completely missing, where each combination is represented by the presence of black dots below the corresponding bar.

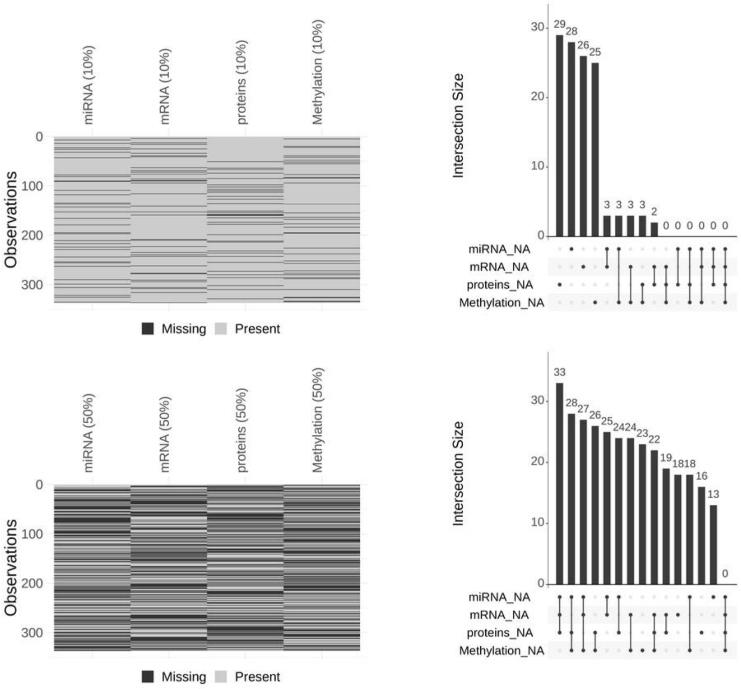


Fig. S15: Missing pattern for PRAD amputated dataset. Upper figure shows 10% of completely missing samples; Lower figure refers to 50% of completely missing samples. For both figures, the left panel shows for each sample in the dataset which data source is missing (black color) and which is present (grey color). Right panel shows a barplot with the number of samples that have a specific combination of data sources completely missing, where each combination is represented by the presence of black dots below the corresponding bar.

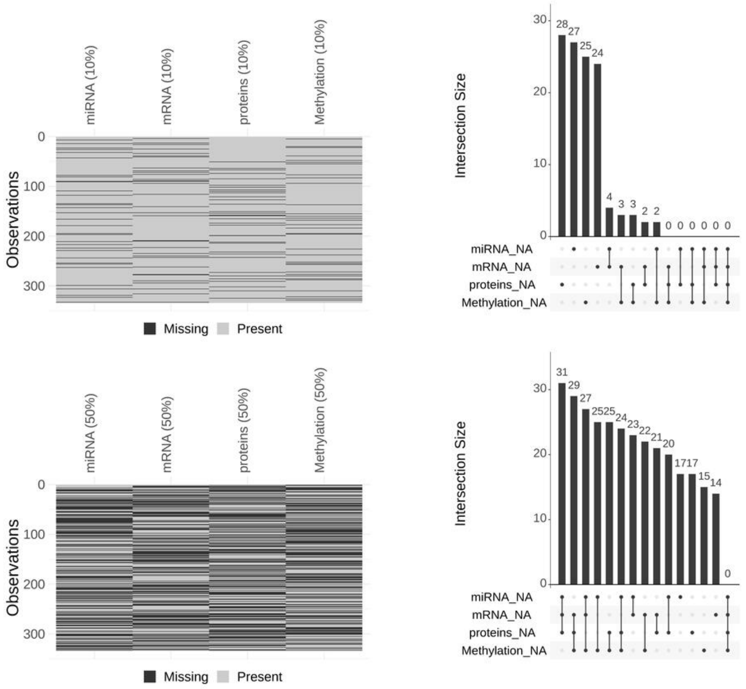


Fig. S16: Missing pattern for SKCM amputated dataset. Upper figure shows 10% of completely missing samples; Lower figure refers to 50% of completely missing samples. For both figures, the left panel shows for each sample in the dataset which data source is missing (black color) and which is present (grey color). Right panel shows a barplot with the number of samples that have a specific combination of data sources completely missing, where each combination is represented by the presence of black dots below the corresponding bar.

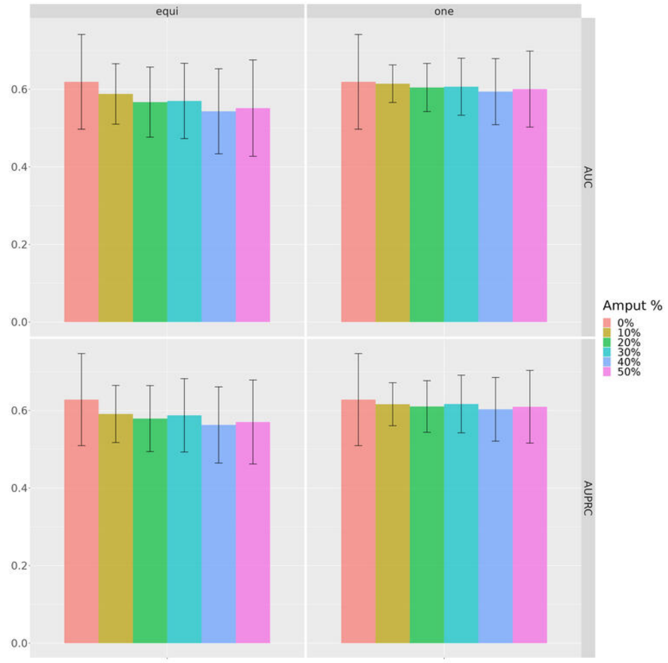


(a) Prediction of OS event

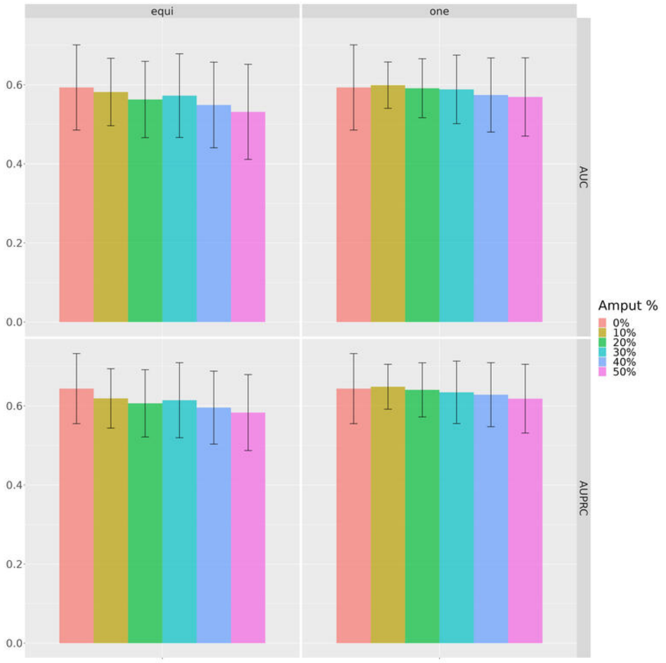


(b) Prediction of PFI event

Fig. S17: Classification results using SNF on complete dataset and miss-SNF with amputated datasets. (a) shows performance for Overall Survival (OS) prediction and (b) reports the performance for Progression Free Interval (PFI). In each plot, the mean performance and pooled standard deviation for different miss-SNF initializations (columns) and metrics (rows) are shown, considering various amputation percentages depicted with distinct colors.
