## Supplementary Figures 2 for "miss-SNF: a multimodal patient similarity network integration approach to handle completely missing data sources"

[1]Dipartimento di Informatica "Giovanni Degli Antoni", Università degli Studi di Milano, Via Giovanni Celoria 18, 20133, Italy, [2]European Commission, Joint Research Centre (JRC), Ispra, Italy, [3]Department of Biomedical Sciences, Humanitas University, Via Rita Levi Montalcini 4, 20072, Pieve Emanuele (MI), Italy, [4]Department of Pathology, IRCCS Humanitas Clinical and Research Hospital, Via Alessandro Manzoni 56, 20089, Rozzano (MI), Italy, [5]Environmental Genomics and Systems Biology Division, Lawrence Berkeley National Laboratory, Berkeley, Ca, USA and [6]ELLIS - European Laboratory for Learning and Intelligent Systems, Milan, Italy

### S10. Supplementary Figures

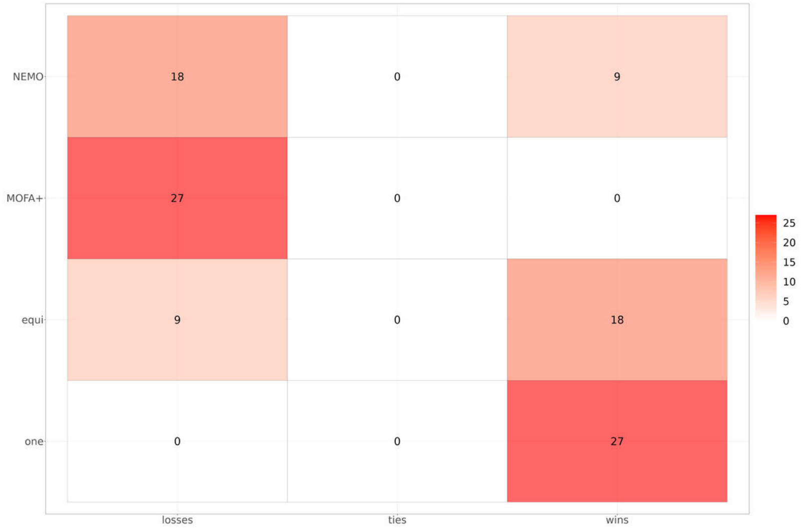


Fig. S18: Win-tie-loss table summarizing the statistical comparison between different data fusion approaches (miss-SNF one and equidistant, MOFA+, NEMO) in the reconstruction of partial datasets. Each element of the win-tie-loss counts the number of losses (left column), ties (center column), and wins (right column) for one of the data fusion algorithms (rows). The wins/ties/losses are counted by using the paired Wilcoxon test ($\alpha = 0.05$) to compare the distributions of the $h = 10$ 1-norm distance values comparing a complete dataset to each of the 10 amputated datasets obtained at a fixed percentage of partial samples. Cancers are considered separately leading to 27 comparisons for each algorithm.

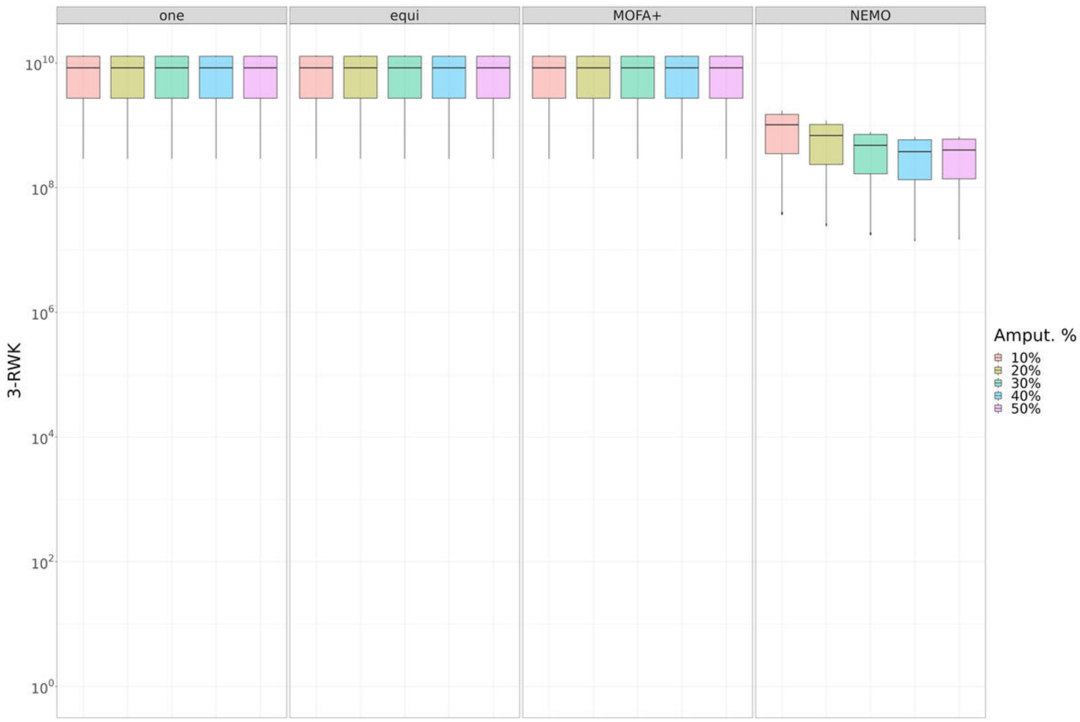


Fig. S19: Topological comparison between the fused PSNs obtained on the complete and partial datasets by different data integration approaches (miss-SNF one and equidistant, MOFA+, NEMO). Each boxplot represents the distribution of the topological similarities for all the nine TCGA datasets. For each dataset we consider the $h = 10$ similarities between the complete dataset and each of the $h = 10$ amputated datasets (randomly generated with the same percentage of partial cases). Similarities between graphs are computed using the 3-step Random Walk Kernel.

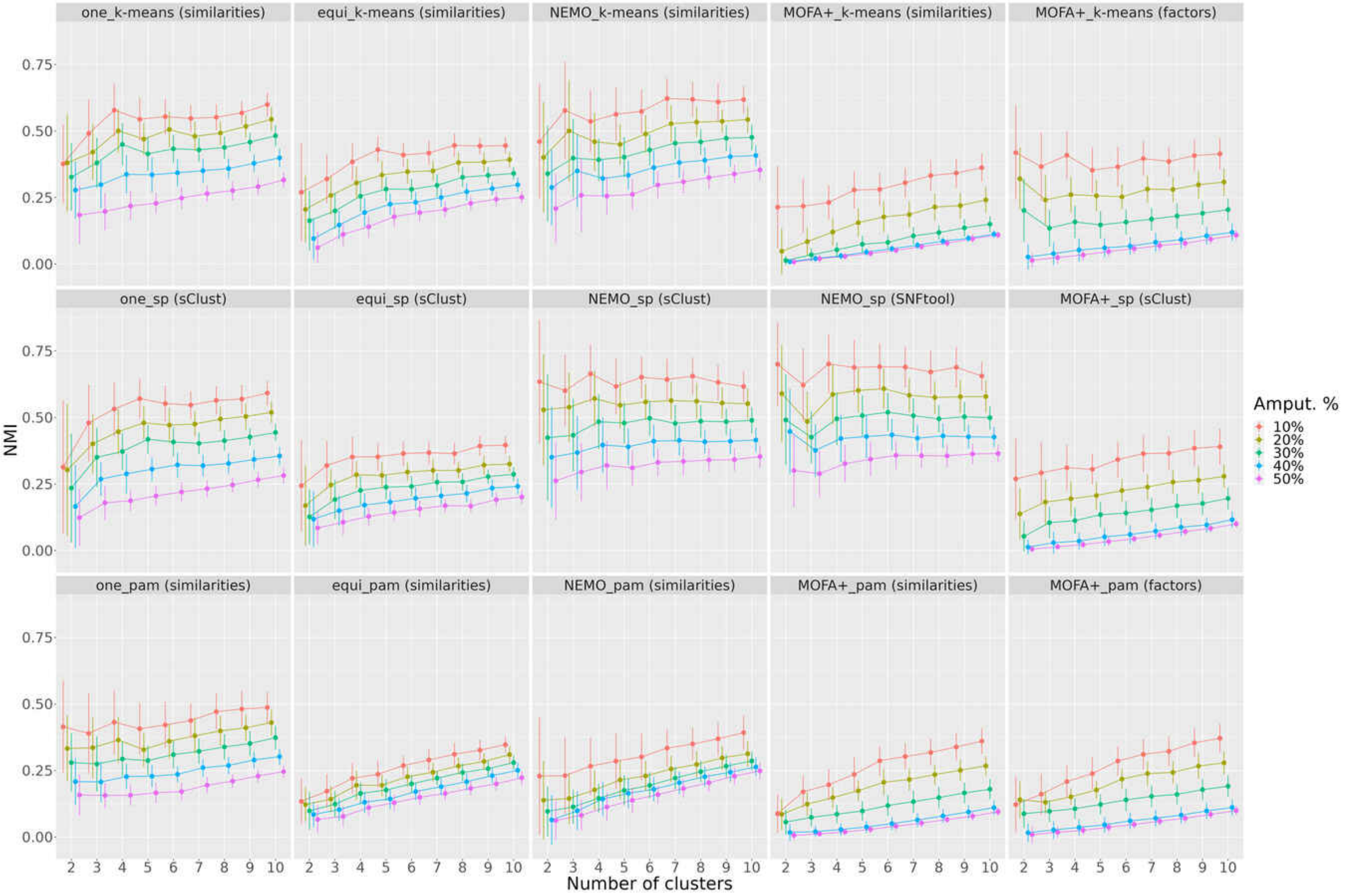


Fig. S20: Comparison of data integration approaches considering different clustering algorithms using NMI. The plot shows for each evaluated number of clusters $nc = [2 .. 10]$ the value of the NMI between clusterings obtained from complete and amputated datasets, where amputation percentages are in different colors. Each panel shows the results of the combination of a data integration approach with a clustering method. Points in the curves are the mean of NMI values obtained from different random repetitions of each amputation percentage and cancers, with their corresponding pooled standard deviation.

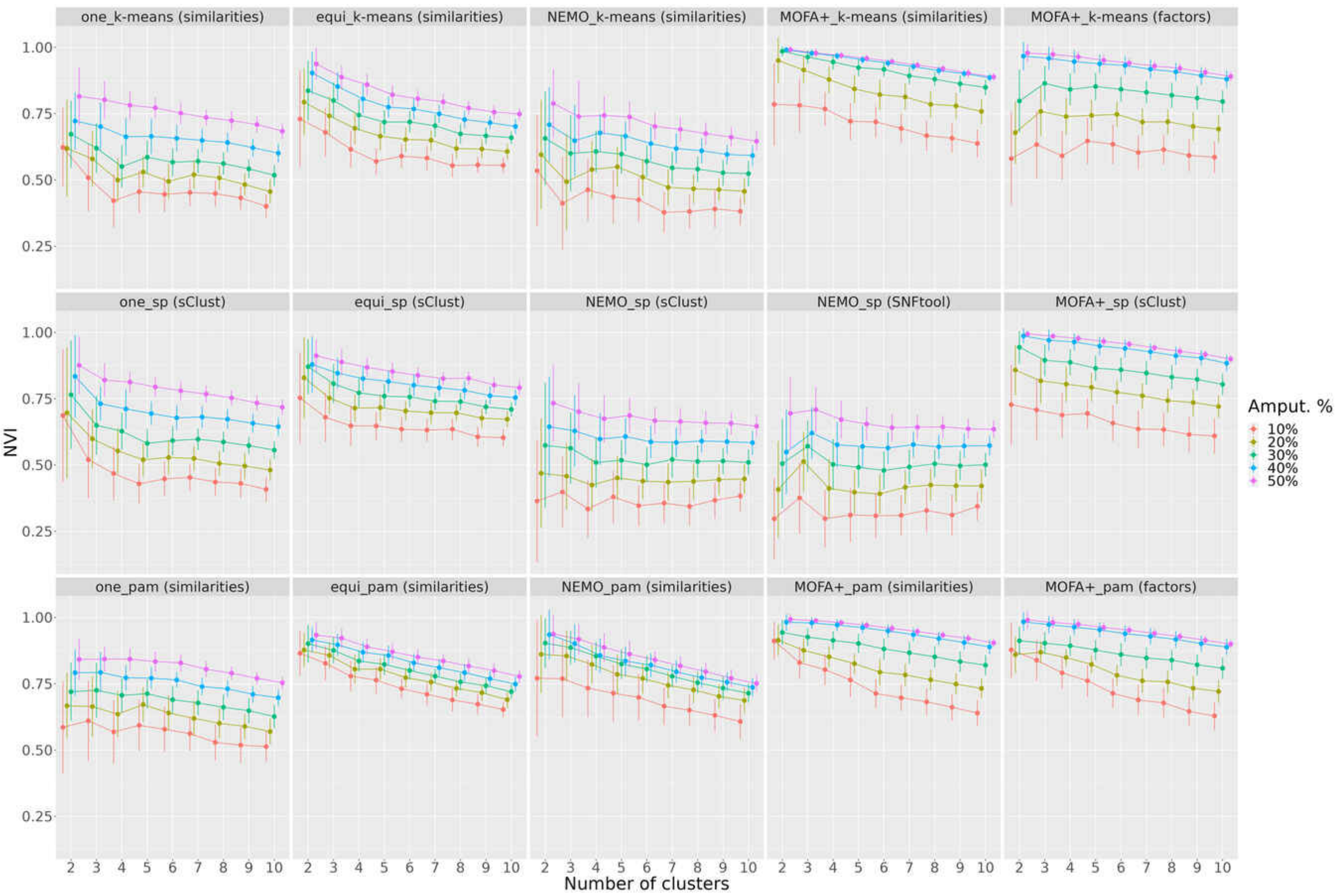


Fig. S21: Comparison of data integration approaches considering different clustering algorithms using NVI. The plot shows for each evaluated number of clusters $nc = [2 \; .. \; 10]$ the value of the NVI computed when clusterings obtained from complete and amputated datasets are considered. The results from amputated datasets having different amputation percentages are represented by colors. Each panel shows the results of the combination of a data integration approach (miss-SNF one, equidistant, NEMO, MOFA+) with a clustering method (k-means, SP and PAM), whose names are separated by an underscore. Points in the curves are the mean of NVI values obtained from different random repetitions of each amputation percentage and cancers, with their corresponding pooled standard deviation.

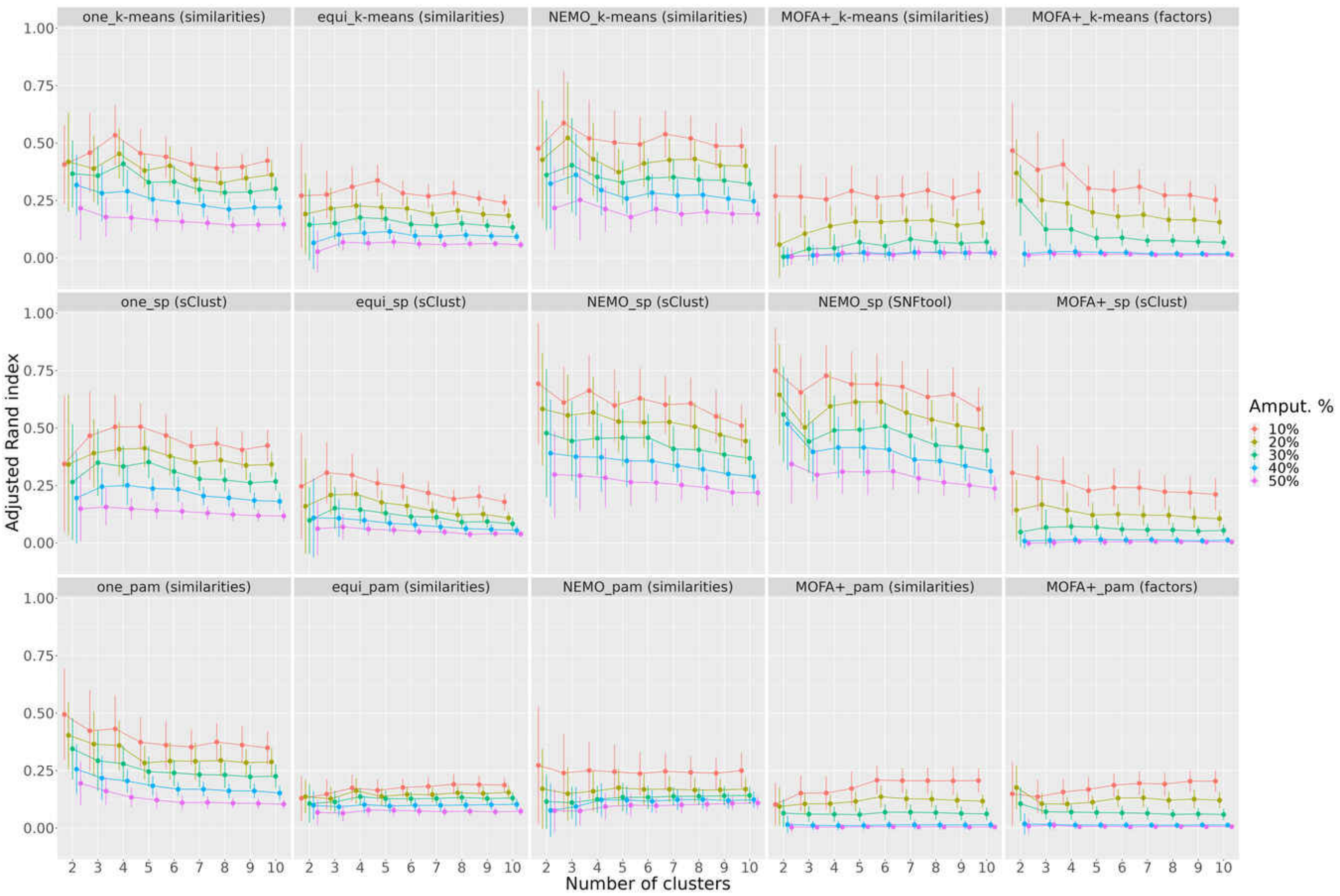


Fig. S22: Comparison of data integration approaches considering different clustering algorithms using Adjusted Rand Index. The plot shows for each evaluated number of clusters $nc = [2 .. 10]$ the value of the Adjusted Rand Index computed when clusterings obtained from complete and amputated datasets are considered. The results from amputated datasets having different amputation percentages are represented by colors. Each panel shows the results of the combination of a data integration approach (miss-SNF one, equidistant, NEMO, MOFA+) with a clustering method (k-means, SP and PAM), whose names are separated by an underscore. Points in the curves are the mean of Adjusted Rand Index values obtained from different random repetitions of each amputation percentage and cancers, with their corresponding pooled standard deviation.

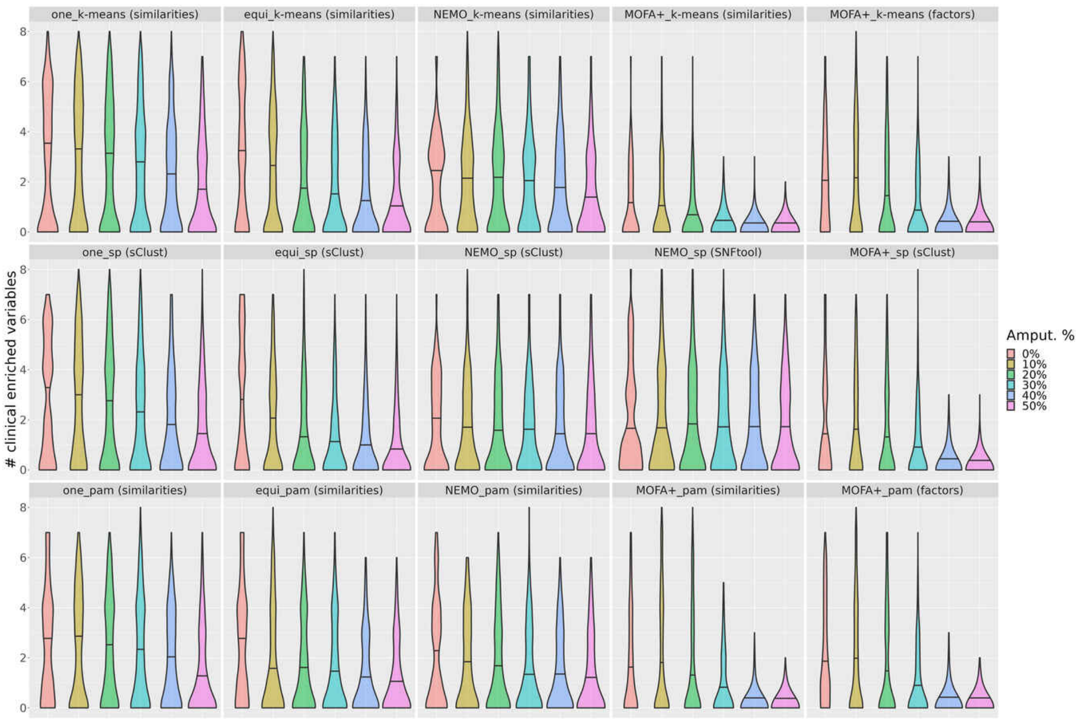


Fig. S23: Comparison of data integration approaches considering different clustering algorithms using the number of enriched clinical variables. The violin plot shows the distribution of the number of enriched clinical variables (Bonferroni adjusted p-value $< 0.05$) considering the results coming from each evaluated number of clusters, random repetitions of each amputation percentage and cancers. The horizontal line in each violin represents the median of the density estimate. Amputation percentages are depicted by different colors and each panel shows the results of a data integration approach with a clustering method.

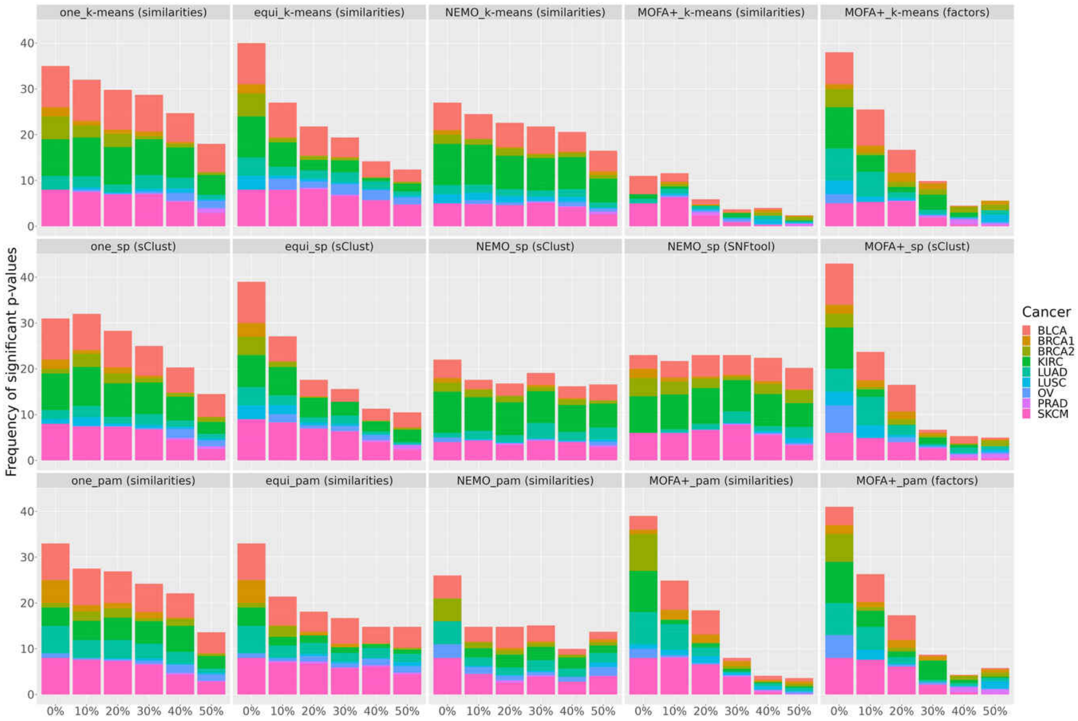


Fig. S24: Comparison of data integration approaches considering different clustering algorithms using the frequency of significant logrank tests. The stacked bar plot shows the frequency of significant logrank tests (p-value $< 0.05$) for the different amputation percentages on the x-axis; the various cancers are depicted by different colors. Each panel shows the results of a data integration approach with a clustering method. This plot is considering all the evaluated numbers of clusters.

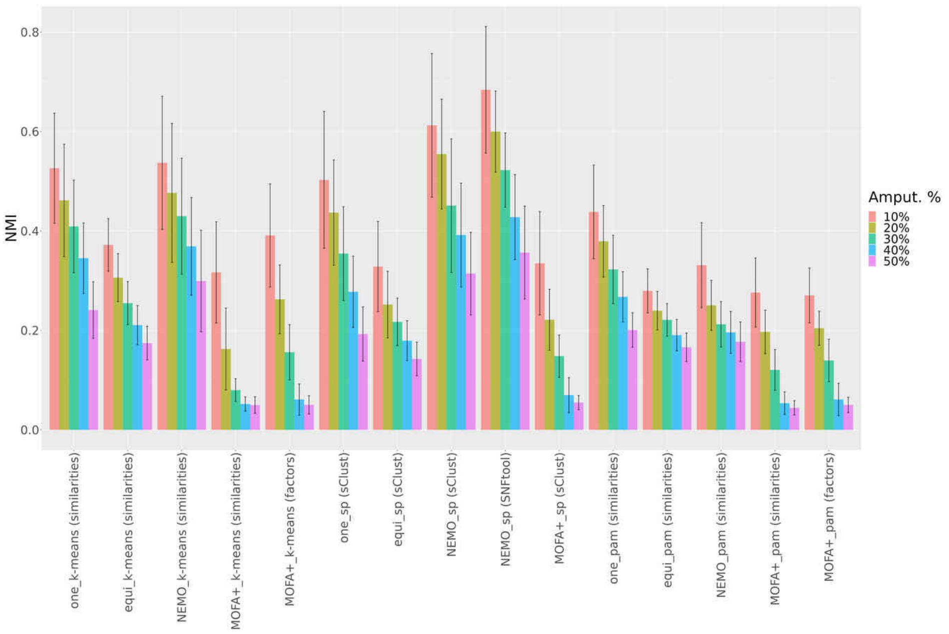


(a) Selection exploiting clin-surv method

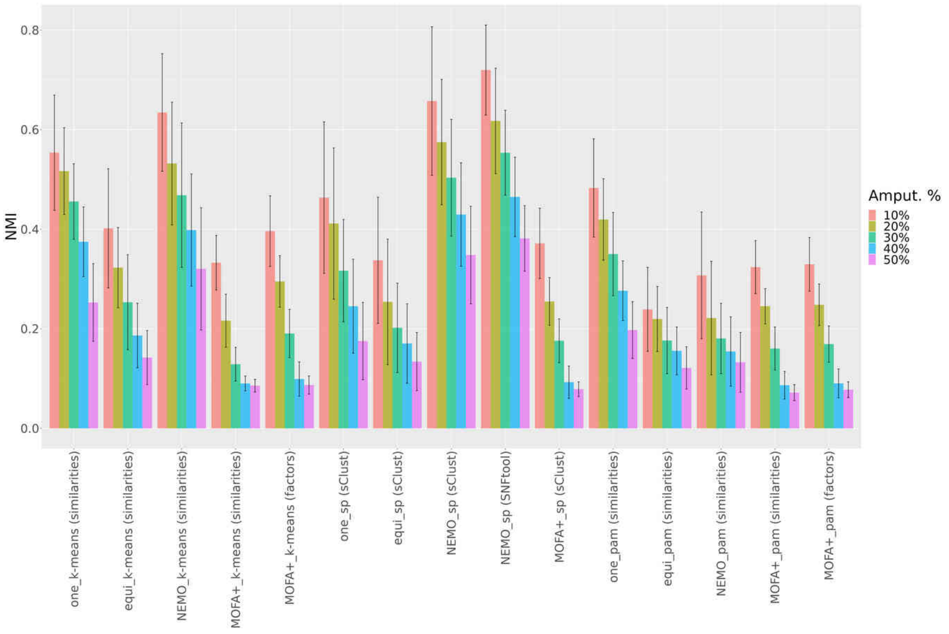


(b) Selection exploiting eigengap method

Fig. S25: Comparison of data integration approaches and clustering algorithms using NMI after selection of the optimal number of clusters. The plots show the value of the NMI computed when clusterings obtained from complete and amputated datasets are considered after the selection of the optimal number of clusters leveraging (a) the clin-surv method and (b) the modified eigengap approach. The results of the evaluated data integration approaches and clustering algorithms are represented on the x-axis while amputation percentages are depicted by different colors. Each bar shows the mean of NMI values obtained from different random repetitions of each amputation percentage and cancers, with its corresponding pooled standard deviation.

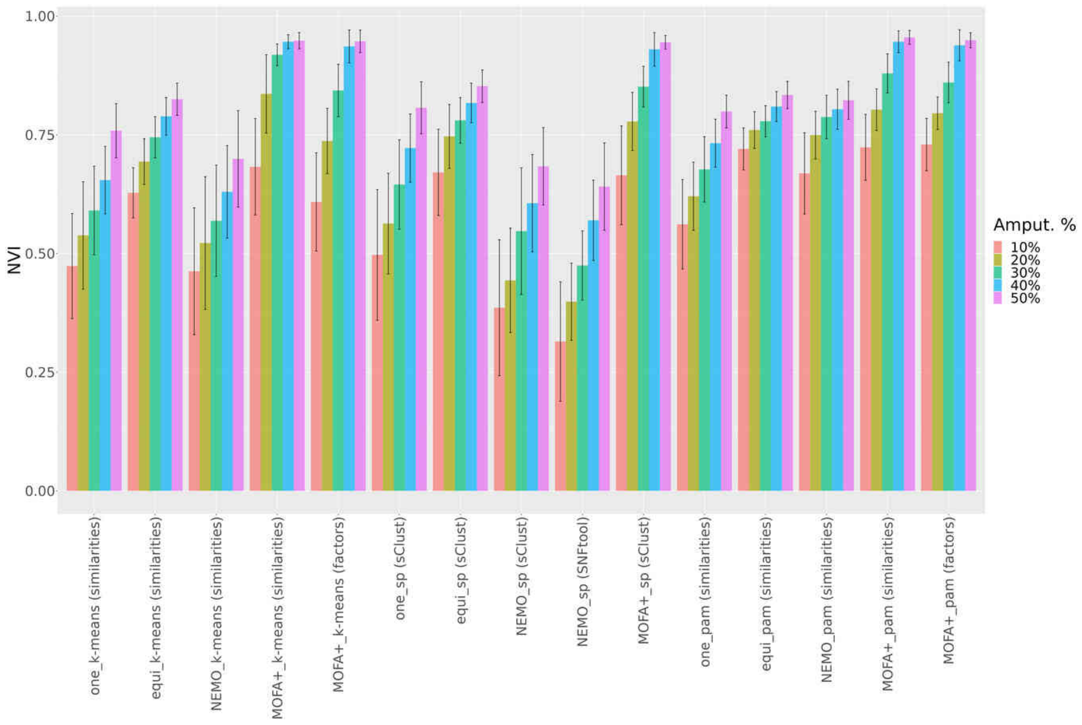


(a) Selection exploiting clin-surv method

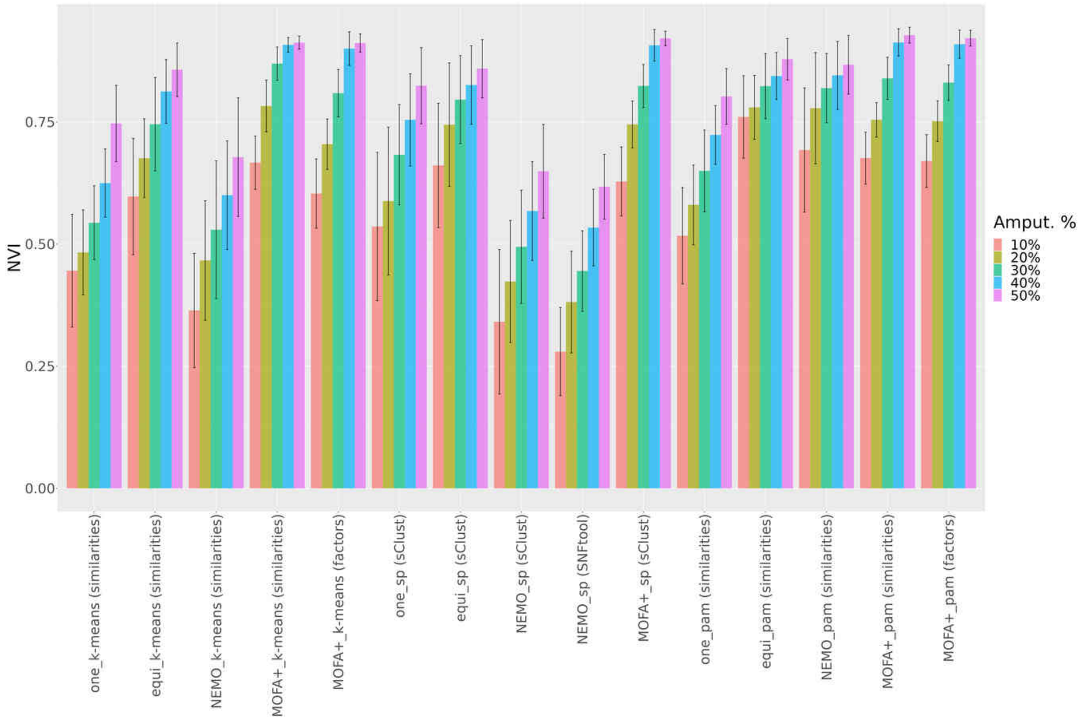


(b) Selection exploiting eigengap method

Fig. S26: Comparison of data integration approaches and clustering algorithms using NVI after selection of the optimal number of clusters. The plots show the value of the NVI computed when clusterings obtained from complete and amputated datasets are considered after the selection of the optimal number of clusters leveraging (a) the clin-surv method and (b) the modified eigengap approach. The results of the evaluated data integration approaches (miss-SNF one, equidistant, NEMO, MOFA+) and clustering algorithms (k-means, SP and PAM), having names separated by an underscore, are represented on the x-axis while different amputation percentages are depicted by different colors. Each bar shows the mean of NVI values obtained from different random repetitions of each amputation percentage and cancers, with its corresponding pooled standard deviation.

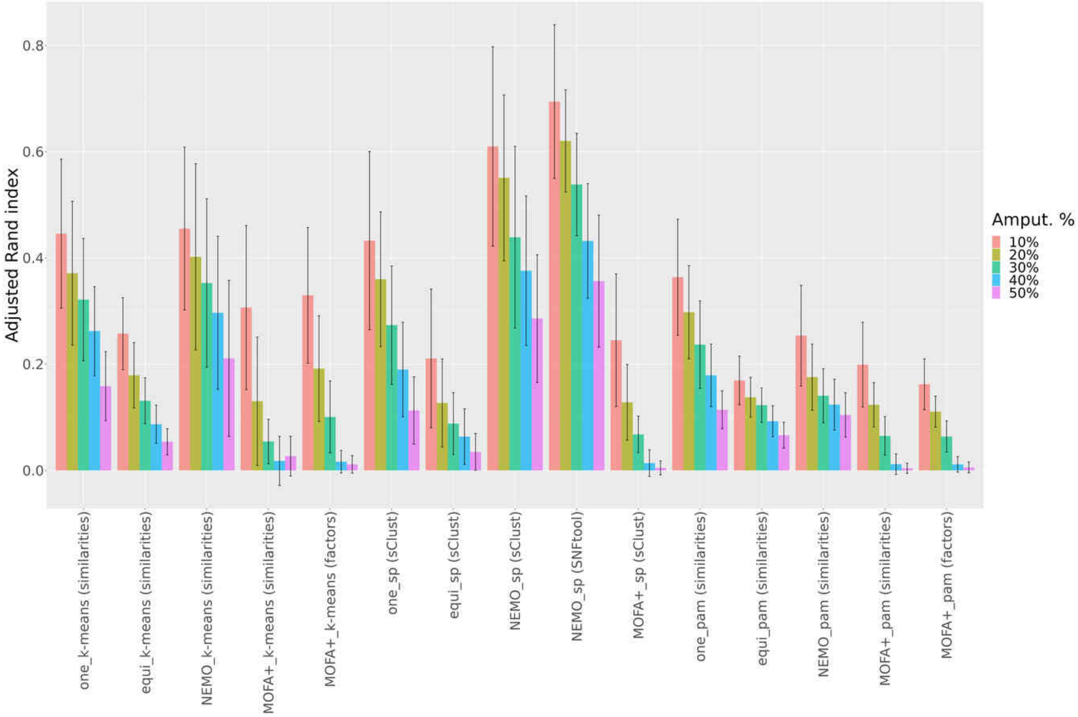


(a) Selection exploiting clin-surv method

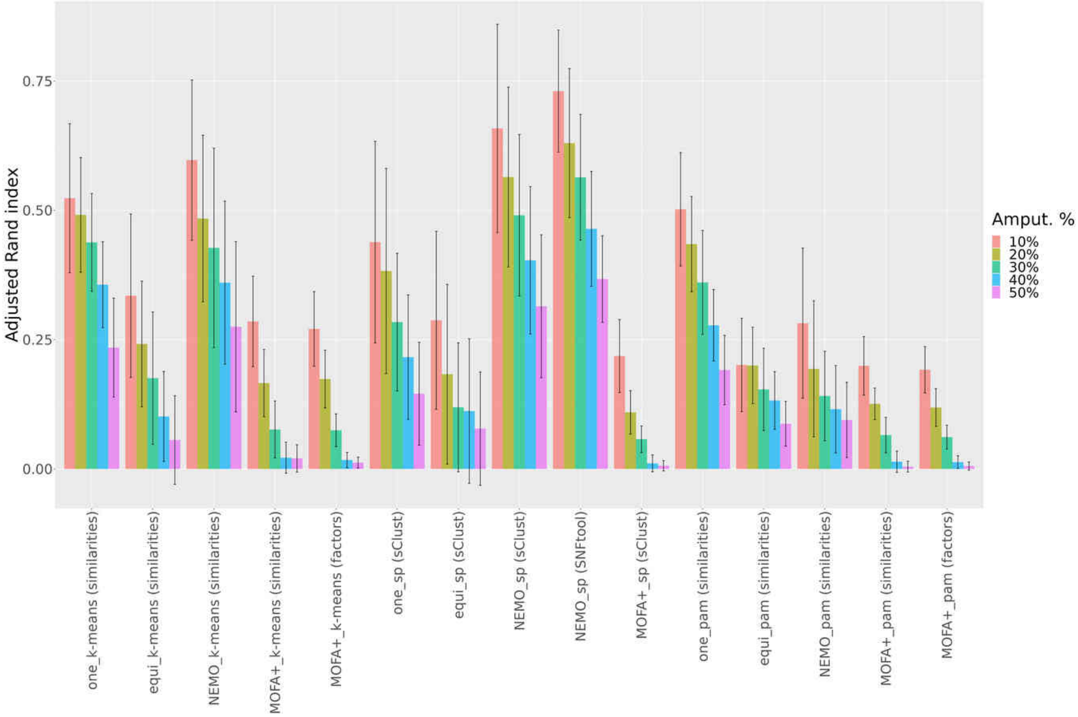


(b) Selection exploiting eigengap method

Fig. S27: Comparison of data integration approaches and clustering algorithms using Adjusted Rand Index after selection of the optimal number of clusters. The plots show the value of the Adjusted Rand Index computed when clusterings obtained from complete and amputated datasets are considered after the selection of the optimal number of clusters leveraging (a) the clin-surv method and (b) the modified eigengap approach. The results of the evaluated data integration approaches (miss-SNF one, equidistant, NEMO, MOFA+) and clustering algorithms (k-means, SP and PAM), having names separated by an underscore, are represented on the x-axis while different amputation percentages are depicted by different colors. Each bar shows the mean of Adjusted Rand Index values obtained from different random repetitions of each amputation percentage and cancers, with its corresponding pooled standard deviation

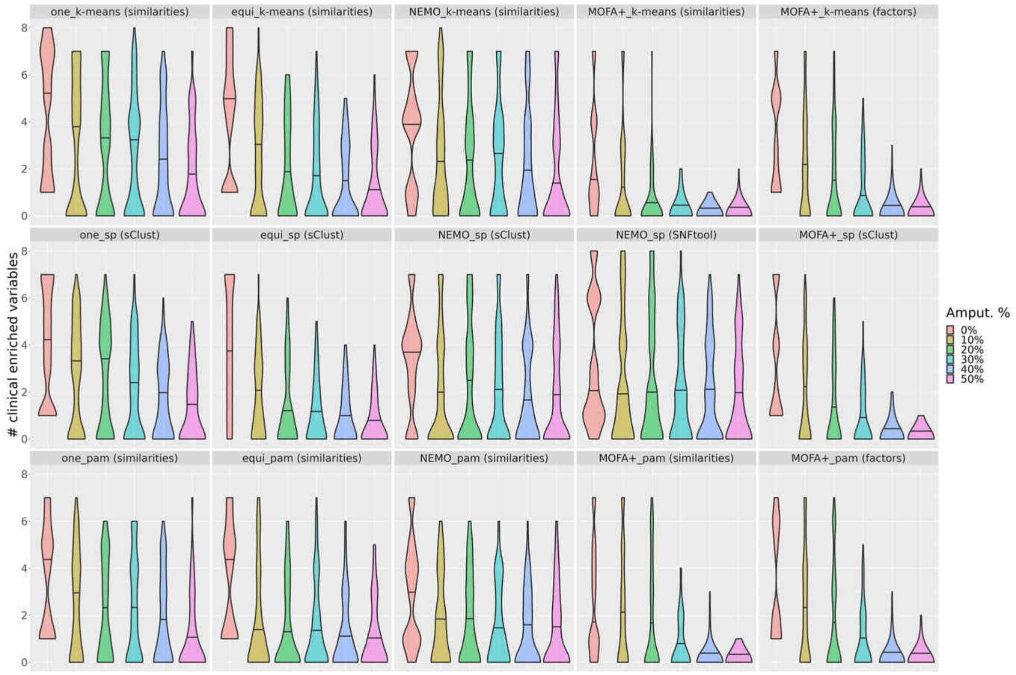


(a) Selection exploiting clin-surv method

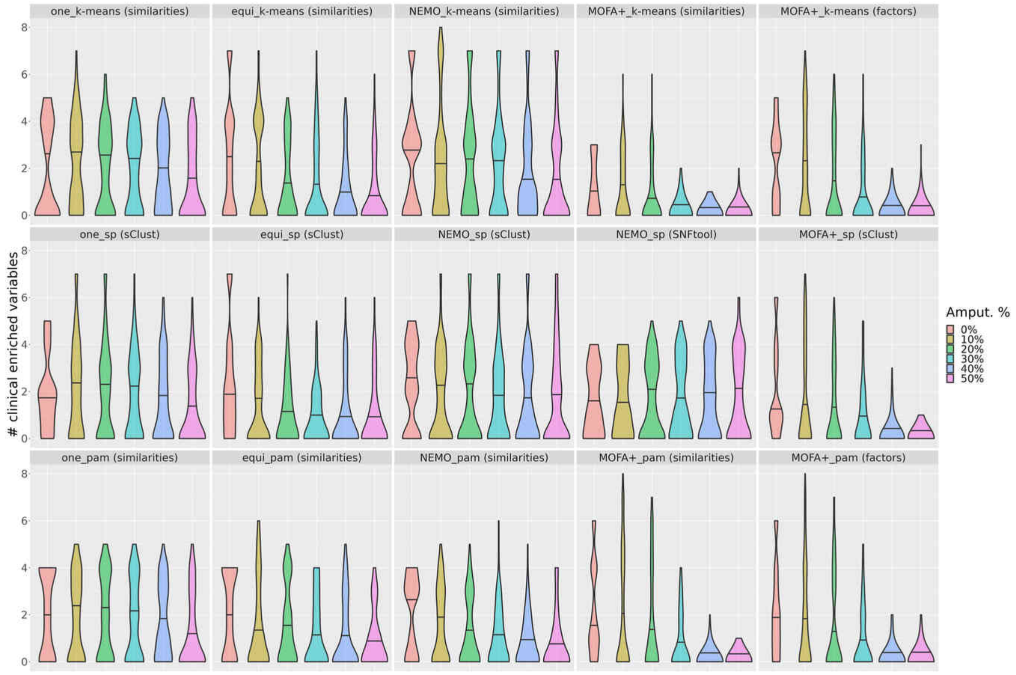


(b) Selection exploiting eigengap method

Fig. S28: Comparison of data integration approaches considering different clustering algorithms using the number of enriched clinical variables and the optimized number of clusters. The violin plot shows the distribution of the number of enriched clinical variables (Bonferroni adjusted p-value $< 0.05$) considering the results coming from the random repetitions of each amputation percentage and cancers. (a) depicts the results when clin-surv method is used to obtain the optimal number of clusters, while (b) reports the results using the eigengap method. The horizontal line in each violin represents the median of the density estimate. The results from amputated datasets having different amputation percentages are depicted by different colors and each panel shows the results of the combination of a data integration approach (miss-SNF one, equidistant, NEMO, MOFA+) with a clustering method (k-means, SP and PAM), whose names are separated by an underscore.

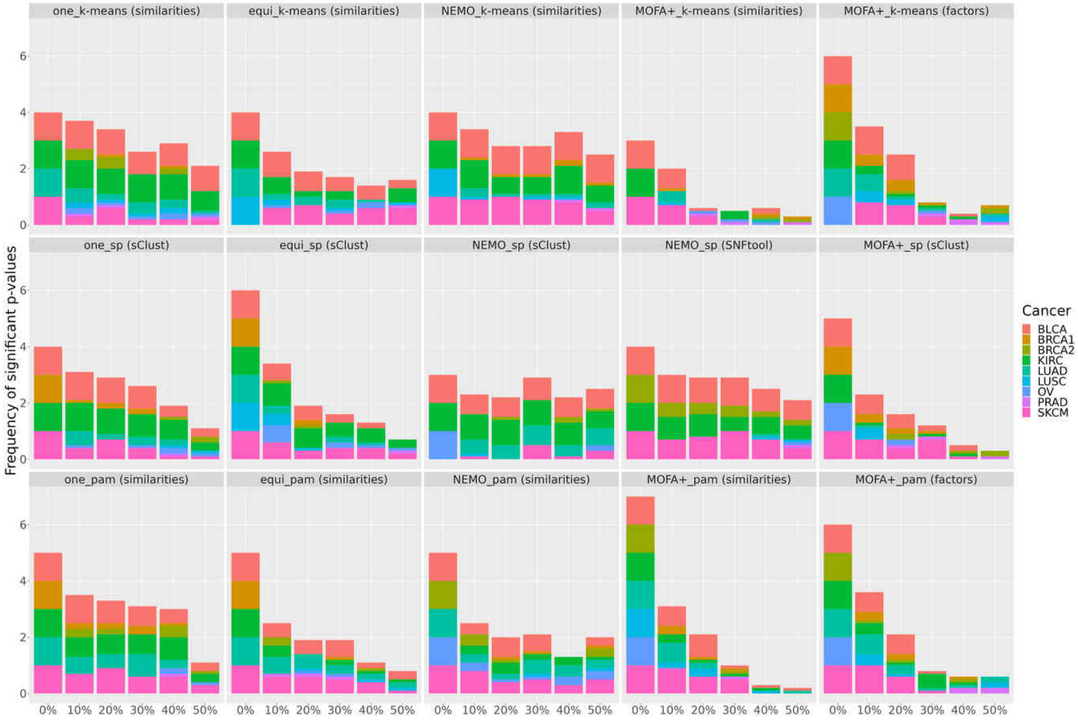


(a) Selection exploiting clin-surv method

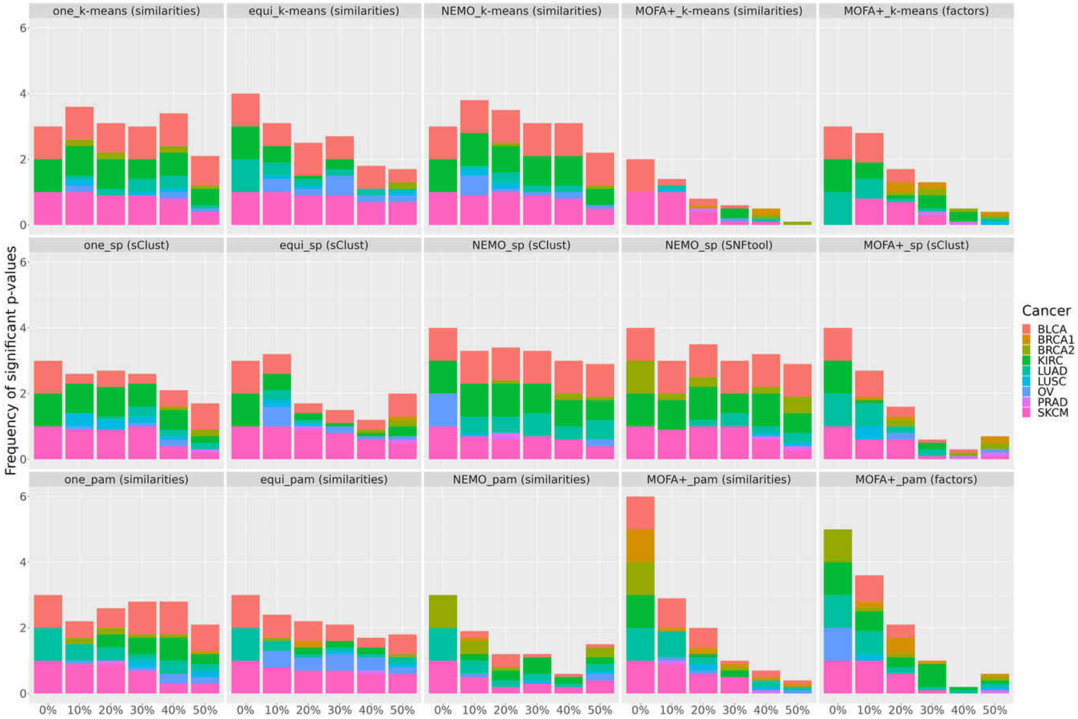


(b) Selection exploiting eigengap method

Fig. S29: Comparison of data integration approaches considering different clustering algorithms using the frequency of significant logrank tests after selection of the optimal number of clusters. The stacked bar plot shows the frequency of significant logrank tests (p-value $< 0.05$) for the different amputation percentages on the x-axis and the various cancers depicted by different colors. Plot (a) reports the results where clin-surv method is applied to select the optimal number of clusters, while plot (b) shows the results for the modified eigengap approach. Each panel represents the results of the combination of a data integration approach (miss-SNF one, equidistant, NEMO, MOFA+) with a clustering method (k-means, SP and PAM), whose names are separated by an underscore.
